# Multimodal transcriptional control of pattern formation in embryonic development

**DOI:** 10.1101/335919

**Authors:** Nicholas C Lammers, Vahe Galstyan, Armando Reimer, Sean A Medin, Chris H Wiggins, Hernan G Garcia

## Abstract

Predicting developmental outcomes from regulatory DNA sequence and transcription factor patterns remains an open challenge in physical biology. Using stripe 2 of the *even-skipped* gene in *Drosophila* embryos as a case study, we dissect the regulatory forces underpinning a key step along the developmental decision-making cascade: the generation of cytoplasmic mRNA patterns via the control of transcription in individual cells. Using live imaging and computational approaches, we found that the transcriptional burst frequency is modulated across the stripe to control the mRNA production rate. However, we discovered that bursting alone cannot quantitatively recapitulate the formation of the stripe, and that control of the *window of time* over which each nucleus transcribes *even-skipped* plays a critical role in stripe formation. Theoretical modeling revealed that these regulatory strategies—bursting and the time window—obey different kinds of regulatory logic, suggesting that the stripe is shaped by the interplay of two distinct underlying molecular processes.

## Introduction

During embryonic development, tightly choreographed patterns of gene expression—shallow gradients, sharp steps, narrow stripes—specify cell fates (Gilbert, 2010). The correct positioning, sharpness, and amplitude of these patterns of cytoplasmic mRNA and protein ensure the reliable determination of animal body plans (Peter and Davidson, 2015). Yet, despite decades of work mapping the gene regulatory networks that drive development and extensive efforts to dissect the regulatory logic of the enhancer elements that dictate the behavior of these networks, the precise prediction of how gene expression patterns and developmental outcomes are driven by transcription factor concentrations remains a central challenge in the field (Vincent et al., 2016).

Predicting developmental outcomes demands a quantitative understanding of the flow of information along the central dogma: how input transcription factors dictate the output rate of mRNA production, how this rate of mRNA production dictates cytoplasmic patterns of mRNA, and how these mRNA patterns lead to protein patterns that feed back into the gene regulatory network. While the connection between transcription factor concentration and output mRNA production rate has been the subject of active research over the last three decades (Lawrence et al., 1987; Driever and Nusslein-Volhard, 1988; Small et al., 1991; Struhl et al., 1992; Jiang and Levine, 1993; Gray et al., 1994; Jaeger et al., 2004; Segal et al., 2008; Levine et al., 2014; Garcia et al., 2016; Vincent et al., 2016; Sayal et al., 2016), the connection between this output rate and the resulting cytoplasmic patterns of mRNA has remained largely unexplored. For example, a stripe of cytoplasmic mRNA within an embryo could arise as a result of radically different transcriptional dynamics at the single-nucleus level (Figure 1A). Specifically, if individual nuclei along this stripe modulate their average RNA polymerase loading rate, then graded control of the mean rate of transcription results: nuclei in the middle of the stripe transcribe at a higher average rate than nuclei on the stripe boundaries (Figure 1B). We identify this graded transcriptional control strategy with the analog control of gene expression. Alternatively, transcription factors could exert control over the length of time a nucleus is transcriptionally active (Figure 1C). In this binary control scheme—akin to an on/off switch that dictates whether a nucleus is transcriptionally active or quiescent—individual nuclei transcribe at the same average rate regardless of their position along the stripe, but for different lengths of time. Finally, some nuclei might not engage in transcription at all during the formation of the pattern (Figure 1D). Here, a larger fraction of nuclei engage in mRNA production in the stripe center than in the boundaries. Any of these scenarios, or some combination thereof, can explain the formation of a cytoplasmic mRNA pattern.

**Figure 1.**
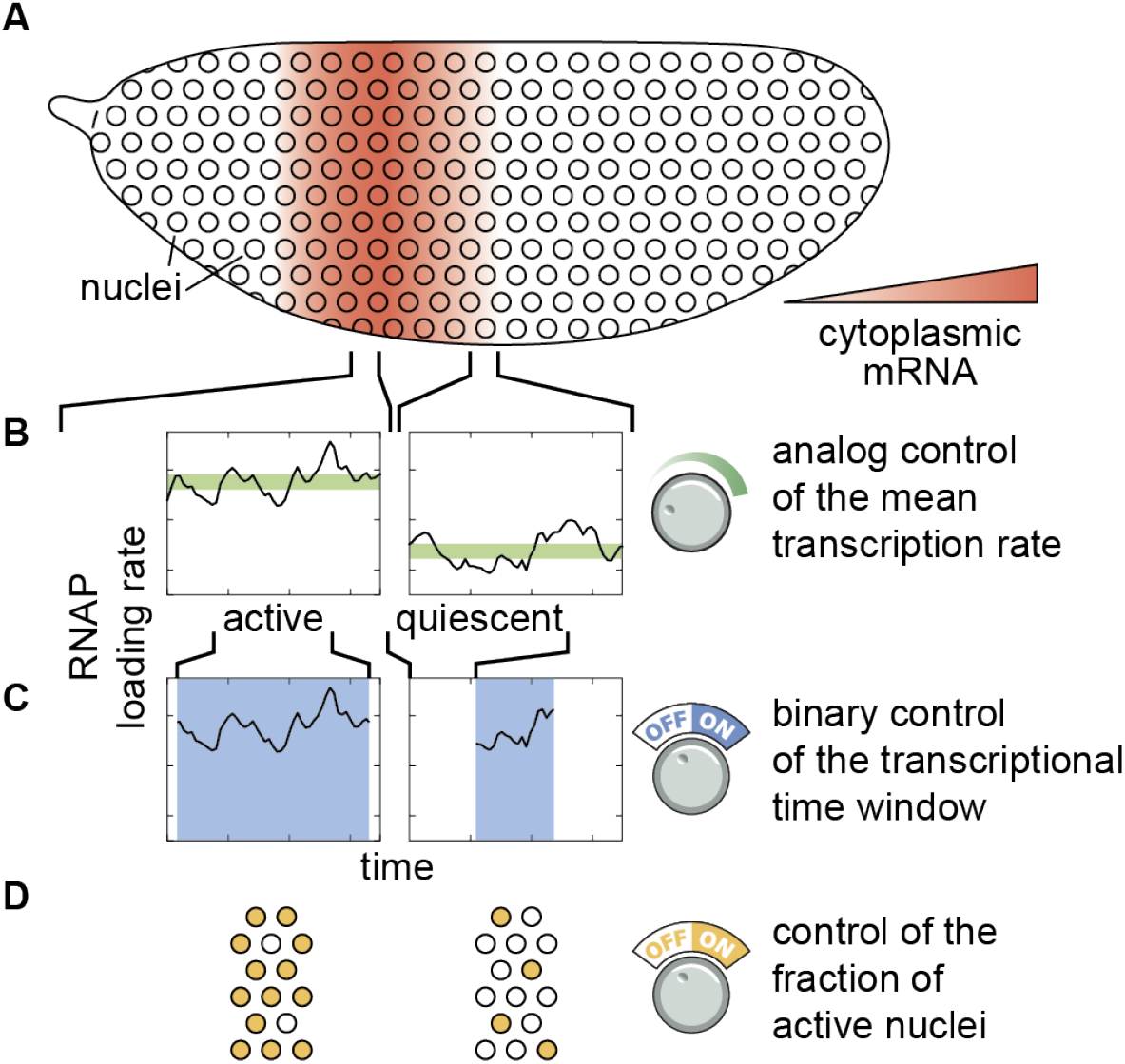
Models of pattern formation by single-cell transcriptional activity. **(A)** Cytoplasmic mRNA patterns could arise from transcription factors exerting control over **(B)** the mean transcription rate, **(C)** the transcriptional time window dictating when a nucleus is transcriptionally active or quiescent, **(D)** the fraction of active nuclei, or some combination thereof.

In order to uncover the quantitative contribution of these three regulatory strategies to pattern formation, and to determine whether other regulatory strategies are at play, it is necessary to measure the rate of RNA polymerase loading in individual nuclei, in real time, in a living embryo. However, to date, most studies have relied on fìxed-tissue techniques such as mRNA FISH and immunofluorescence in order to obtain snapshots of the cytoplasmic distributions of mRNA and protein as development progresses Jaeger et al., 2004; Fakhouri et al., 2010; Parker et al., 2011; Estrada et al., 2016; Crocker et al., 2016; Verd et al., 2017; Park et al., 2018). Such techniques are virtually silent regarding the regulation of single-cell gene expression over time, and are thus ill-suited to the study of how spatiotemporal variations in transcriptional dynamics give rise to patterns of cytoplasmic mRNA.

In this work, we investigated how single-cell transcriptional activity leads to the formation of stripe 2 of the widely studied *even-skipped* (*eve*) gene in the developing fruit fly embryo (Small et al., 1992; Arnosti et al., 1996). We combined single-cell live imaging with theoretical modeling in order to study transcriptional activity at the single-cell level in real time, seeking a quantitative connection between the spatiotemporal regulation of transcription and the formation of cytoplasmic patterns of mRNA. Consistent with previous studies, we found that the rate of mRNA production is elevated in the center of the stripe (Bothma et al., 2014). Strikingly, however, we discovered that this analog control is alone insufficient to quantitatively recapitulate the formation of the stripe; binary control of the transcriptional time window (Figure 1C) is also necessary. Furthermore, we developed novel computational approaches to uncover the molecular underpinning of each regulatory strategy. We employed a memory-adjusted hidden Markov model (mHMM) to uncover variations in transcriptional dynamics in individual nuclei across space and time (Suter et al., 2011; Molina et al., 2013; Corrigan et al., 2016). We showed that, consistent with previous results, transcription factors control the rate of transcription by altering the frequency of transcriptional bursts (Fukaya et al., 2016; Zoller et al., 2018). Finally, we utilized logistic regressions to correlate *eve* stripe 2 transcriptional dynamics with changes in input transcription factor concentrations. This analysis revealed that the transcriptional time window adheres to different regulatory logic than transcriptional bursting: while repressor levels alone were sufficient to explain the early silencing of nucei in the anterior and posterior stripe flanks, the control of bursting among transcriptionally engaged nuclei depends upon the input concentrations of both activators and repressors. Thus, our findings point to the presence of two distinct regulatory mechanisms that control transcription and gene expression patterns in early development, showcasing the potential for theoretical modelling and biological numeracy to yield novel biological insights when coupled with precise and quantitative experimental observation.

## Results

### Predicting cytoplasmic mRNA distributions from transcriptional activity

To predict how the transcriptional activity of individual nuclei dictates the formation of cytoplasmic patterns of mRNA, we began with a simple model that considers the balance between the rate of mRNA synthesis and degradation

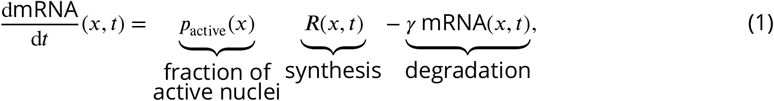

where mRNA(*x*,*t*) indicates the mRNA concentration at position *x* along the embryo at time *t*, *R*(*x*,*t*) corresponds to the mRNA synthesis rate averaged over multiple nuclei within the same position *x*, *p*_active_(*x*) is the fraction of active nuclei (corresponding to the regulatory strategy shown in Figure 1D) and *γ* is the degradation rate (see Appendix 1 for details of this derivation).

In order to examine the quantitative consequences of the three potential regulatory strategies (Figure 1B-D), we adopted widespread assumptions in the modeling of transcriptional regulation (Phillips et al., 2013). First, we assumed that the degradation rate *γ* is a constant and not under any kind of spatiotemporal control. Comparisons between model predictions and empirically measured levels of cytoplasmic mRNA suggest that this assumption is reasonable (see Appendix 2). Second, we posited that at each position throughout the embryo the synthesis rate *R*(*x*, *t*) does not vary significantly in time such that it can be approximated by its time average *R*(*x*) = 〈*R*(*x*,*t*)〉. This assumption will be revised later in the text in order to account for the time-dependent regulation of the mean rate of transcription. Finally, we assumed that nuclei along the axis of the embryo start transcribing at time *t*_on_(*x*), and stop transcribing and enter a state of transcriptional quiescence at time *t*_off_(*x*). Under these assumptions, Equation 1 can be solved analytically, resulting in

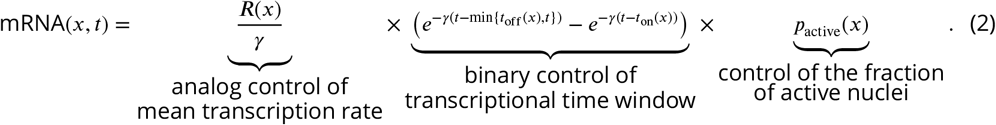

This equation makes precise predictions about how each regulatory strategy contributes to the formation of the cytoplasmic mRNA pattern. Thus, measuring how each quantity is regulated across the stripe allows us to predict their relative contributions to pattern formation.

### Binary control of the transcriptional time window is the primary driver of stripe formation

In order to test the simple model of pattern formation put forward in Equation 2, we quantified transcription of stripe 2 of *eve* in the fruit fly. We imaged the transcription of an *eve* stripe 2 reporter using the MS2 system (Garcia et al., 2013; Lucas et al., 2013; Bothma et al., 2014). Transcripts of a reporter gene driven by the *eve* stripe 2 enhancer and the *eve* promoter contain repeats of a DNA sequence that, when transcribed, form stem loops (Bertrand et al., 1998). These stem loops are recognized by maternally provided MS2 coat protein fused to GFP (Figure 2A). As a result, sites of nascent transcript formation appear as fluorescent puncta within individual nuclei (Figure 2B and Video 1). This fluorescence can be calibrated using single-molecule FISH in order to estimate the number of RNA polymerase molecules actively transcribing the gene (see Materials and Methods and Garcia et al. (2013)). By aligning multiple embryos (Figure2-Figure Supplement 1), we obtained the average number of actively transcribing RNA polymerase molecules as a function of time and position throughout the embryo (Figure 2C).

**Figure 2.**
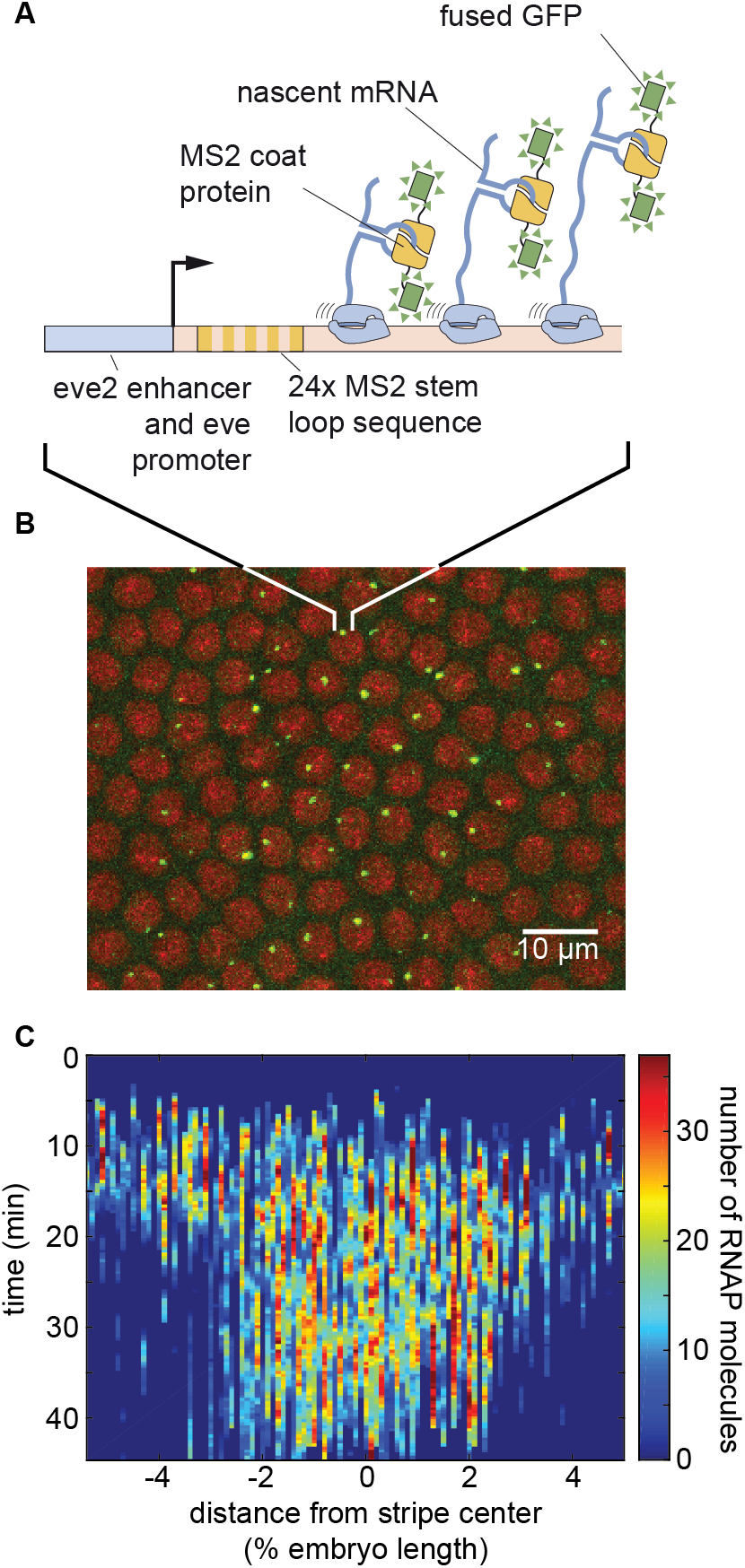
Measuring transcriptional dynamics of *eve* stripe 2 formation using the MS2 system. **(A)** MS2 stem loops introduced in an *eve* stripe 2 reporter gene are bound by MS2 coat protein fused to GFP. **(B)** Sites of nascent transcript formation appear as green fluorescent puncta whose intensity reports on the number of actively transcribing RNA polymerase molecules. Nuclei are visualized through a fusion of RFP to Histone. **(C)** Mean number of RNA polymerase molecules actively transcribing the gene as a function of space and time. (C, data averaged over 11 embryos). **Figure 2–Figure supplement 1. Aligning stripes from multiple embryos.** **Figure 2–Figure supplement 2. Integrating MS2 Spots.**

Using the MS2 system, we quantified each potential regulatory strategy and determined its predicted contribution to pattern formation according to our model in Equation 2. We first used our data to estimate the time-averaged rate of RNA polymerase loading, *R*(*x*) (see Appendix 2 for details). We found that this rate is modulated along the axis of the embryo (Figure 3A and B; see also Video 2, Figure 3–Figure Supplement 1 and Materials and Methods): whereas in the center of the stripe RNA polymerase molecules are loaded at a rate of approximately 16 molecules/min, this loading rate decreases to about 8 molecules/min at the boundaries.

**Figure 3.**
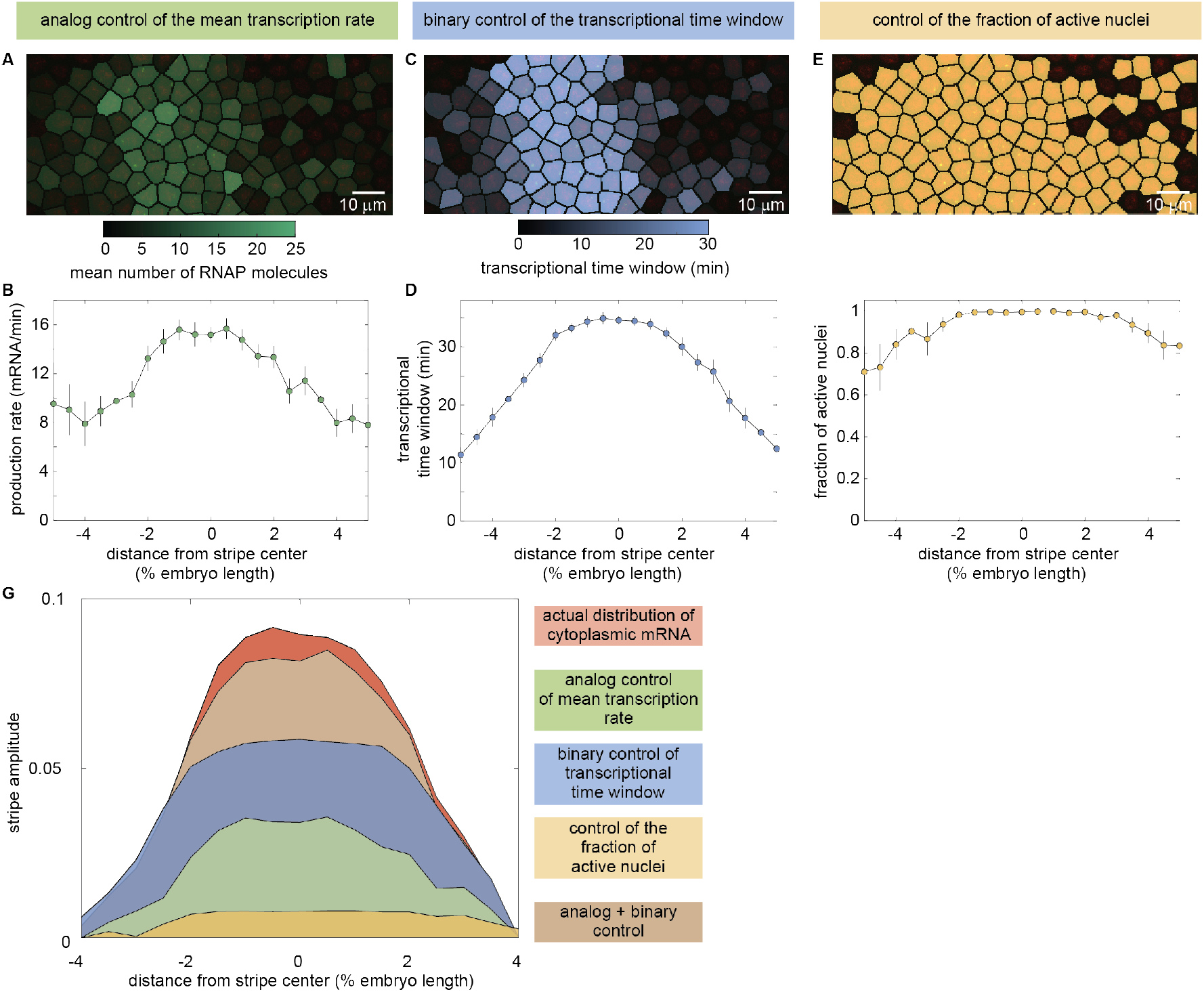
Regulatory strategies for pattern formation in *eve* stripe 2. **(A,B)** Time-averaged rate of mRNA production, **(C,D)** transcriptional time window, and **(E,F)** fraction of active nuclei as a function of position along the embryo. **(G)** Amplitude of the cytoplasmic mRNA distribution compared to the contributions to stripe formation of the analog control of the mean transcription rate, the binary control of the transcriptional time window, and the control of the fraction of active nuclei. The combined contribution from the analog and binary strategies is also shown. See Figure 3–Figure Supplement 3 for details of how depicted profiles were derived from raw data. (A,C,E, representative snapshots of an individual embryo 40 min into nuclear cycle 14; B,D,F, average over 11 embryos, error bars indicate bootstrap estimate of the standard error of the mean). **Figure 3–Figure supplement 1. Mean transcriptional activity over time.** **Figure 3–Figure supplement 2. Regulation of the transcriptional time window.** **Figure 3–Figure supplement 3. Definition of stripe amplitude.** **Figure 3–Figure supplement 4. Joint effect of mean rate, binary control, and fraction of active nuclei.**

Our data also revealed that the transcriptional time window is modulated along the stripe (Figure 3–Figure Supplement 2A). Whereas the time at which each nucleus becomes transcriptionally active, *t*_on_(*x*), was constant across the stripe, with all nuclei becoming active 9 ± 4 min after the previous anaphase (Figure 3–Figure Supplement 2B), the time at which nuclei stop transcribing and become quiescent, *t*_off_(*x*), showed a strong modulation along the embryo’s axis (Figure 3–Figure Supplement 2C). As a result, the time window over which each punctum is engaged in transcription, Δ*t* = *t*_off_ – *t*_on_, is sharply modulated along the stripe (Figure 3C and D and Video 3), with nuclei in the stripe center transcribing for >30 min and nuclei on the boundaries only transcribing for approximately 10 min.

Finally, our analysis also revealed the magnitude of the modulation of the fraction of active nuclei along the stripe. Most nuclei along the stripe were engaged in transcription. In the stripe center, around 90% of nuclei transcribed at some point during the nuclei cycle. This number reduced to about 70% at the boundaries (Figure 3E and F and Video 4).

The analysis in Figure 3A-F reveals that each of the three regulatory strategies identified in Figure 1 is at play in the embryo, and that they all have the potential to contribute to pattern formation. However, these measurements alone cannot inform us on *how much* each of these strategies contributes to the cytoplasmic mRNA pattern. To quantify the degree to which each regulatory strategy contributes to the formation of *eve* stripe 2, we employed the model described in Equation 2.

Figure 3G indicates the quantitative contribution of each regulatory strategy (each term on the right-hand side of Equation 2) to the formation of this cytoplasmic pattern. The cytoplasmic pattern of mRNA, corresponding to left-hand side of Equation 2, was obtained from our live-imaging data (see Appendix 2 for details). Regulation of the fraction of active nuclei along the embryo (Figure 3G, yellow) contributes negligibly to this mRNA pattern. In contrast, both the analog regulation of the mean rate (Figure 3G, green) and the binary control of the transcriptional time window (Figure 3G, blue) make significant contributions to the overall pattern, with binary control playing the dominant role. We thus concluded that the joint effect of these two strategies (Figure 3G, brown) is sufficient to quantitatively recapitulate the stripe of cytoplasmic mRNA from single-cell transcriptional activity.

### Mean transcription rate is dictated by bursting through modulation of the rate of promoter turn on

Are the binary and analog control strategies driven by distinct molecular mechanisms, or are they different manifestations of the same underlying process? To uncover the molecular mechanism behind the analog control of the mean rate of transcription, we analyzed the transcriptional activity of individual nuclei. Previous work demonstrated that the rate of gene expression at individual loci within the *eve* stripe 2 pattern is highly stochastic (Bothma et al., 2014). Indeed, as shown in Figure 4A, our data revealed punctuated peaks and troughs in the number of active RNA polymerase molecules. These features have been related to the rate of RNA polymerase loading at the *eve* promoter by assuming that promoter loading is “burst-like”, with the promoter loading RNA polymerase molecules onto the gene at a constant rate over discrete periods of time (Bothma et al., 2014). This and other evidence from live imaging (Bothma et al., 2014; Fukaya et al., 2016; Desponds et al., 2016), as well as data from fixed-tissue approaches (Pare et al., 2009; Little et al., 2013; Xu et al., 2015; Zoller et al., 2018), support a minimal two-state model of promoter switching (Figure 4B): promoters switch stochastically between ON and OFF states with rates *k*_on_ and *k*_off_. In this model, promoters in the ON state engage in mRNA production at rate *r*.

**Figure 4.**
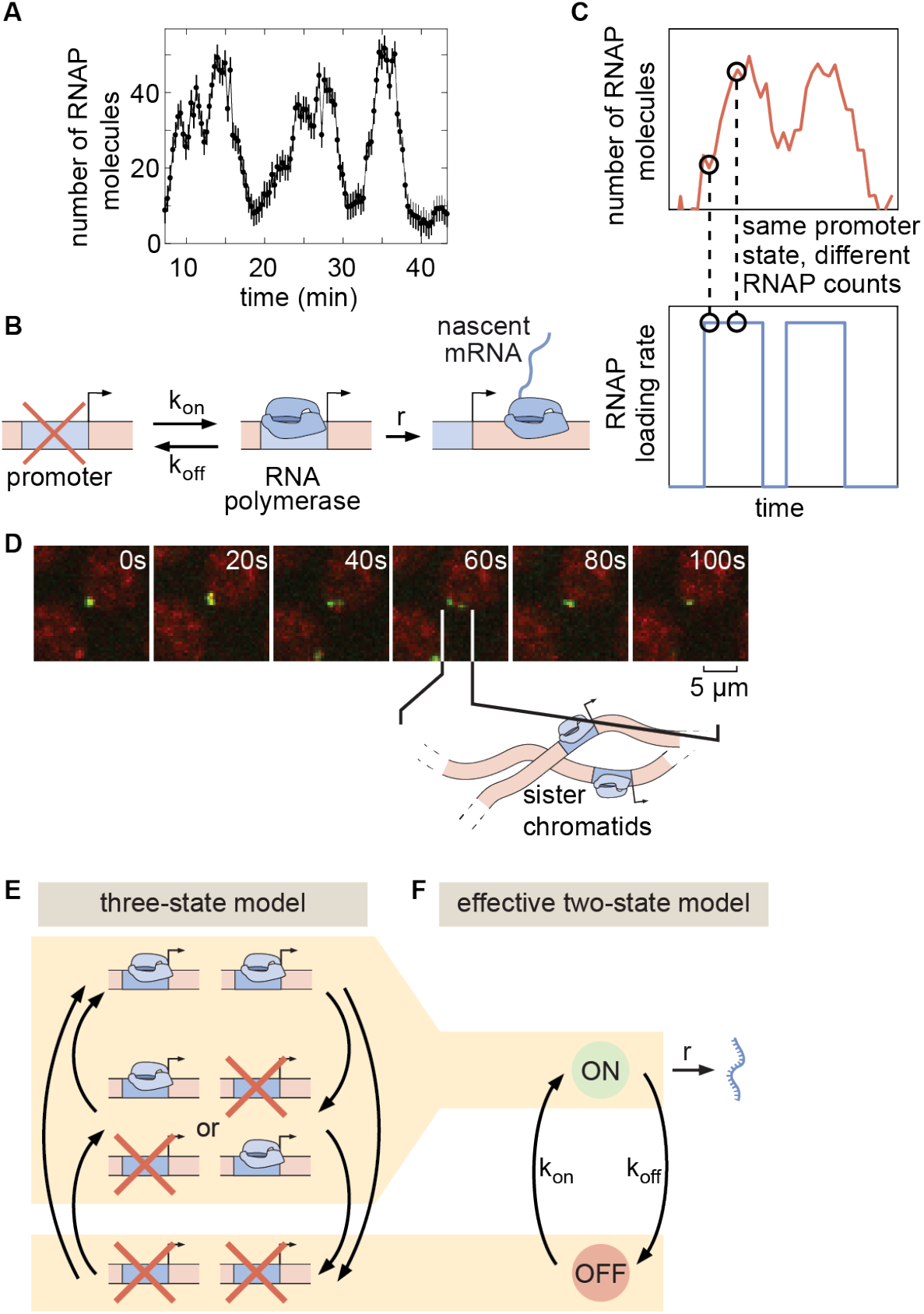
Transcriptional bursting in *eve* stripe 2. **(A)** Single-nucleus measurements reveal that nuclei transcribe in bursts. **(B)** Two-state model of bursting of a single promoter. **(C)** The same hidden rate of RNA polymerase loading (bottom) can correspond to different observable numbers of RNA polymerase molecules on the gene (top), such that standard Hidden Markov model approaches cannot be used to infer the hidden promoter state. **(D)** Fluorescent puncta are composed of two distinct transcriptional loci within a diffraction-limited spot, each corresponding to a sister chromatid. **(E)** Three-state model of promoter switching within a fluorescent punctum that accounts for the combined action of both sister chromatids. **(F)** Effective two-state model of transcriptional bursting. (A, error bars obtained from estimation background fluorescent fluctuations; Materials and Methods and Garcia et al. (2013).)

In the bursting model, the mean rate of transcription is given by the product of the fraction of time spent in the ON state with the transcription rate in this active state (Peccoud and Ycart, 1995; Kepler and Elston, 2001; Sasai and Wolynes, 2003; Sanchez and Kondev, 2008; Sanchez et al., 2011; Xu et al., 2016)

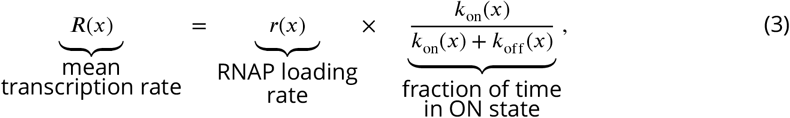

where all parameters are allowed to vary as a function of position along the embryo, *x* (see Appendix 1 for details of this derivation). Thus, within this framework, the observed modulation of the mean rate of transcription across the stripe (Figure 3G, green) implies that one or more of these bursting parameters is subject to spatially controlled regulation. However, the mean rate trend alone is not sufficient to identify *which* of the three bursting parameters (*k*_on_, *k*_off_, and *r*) is being regulated by the input transcription factors in order to control the average transcription rate.

Typically, the *in vivo* molecular mechanism of transcription factor action is inferred from measurements of transcriptional noise obtained through snapshots of dead and fixed embryos or cells using theoretical models (Zenklusen et al., 2008; So et al., 2011; Little et al., 2013; Jones et al., 2014; Senecal et al., 2014; Xu et al., 2015; Padovan-Merhar et al., 2015; Skinner et al., 2016; Bartman et al., 2016; Zoller et al., 2018; Hendy et al., 2017). In contrast, MS2-based live imaging can directly inform on the dynamics of transcriptional bursting in real time. The MS2 approach, however, reports on the total number of actively transcribing RNA polymerase molecules and not on the instantaneous rate of RNA polymerase loading at the promoter, which is the relevant quantity for estimating *k*_on_, *k*_off_, and *r*. To date, approaches for extracting bursting parameters from such data in multicellular organisms have mainly relied on the manual analysis of single-nucleus transcriptional dynamics (Bothma et al., 2014; Fukaya et al., 2016) or autocorrelation-based methods that infer mean bursting parameters across ensembles of traces (Larson et al., 2011; Coulon et al., 2014; Desponds et al., 2016). A computational method for inferring the rates of RNA polymerase loading (Figure 4C, bottom) from the total number of actively transcribing RNA polymerase molecules in single cells (Figure 4C, top) is thus needed to obtain the bursting parameters.

Hidden Markov models (HMMs) are widely used to uncover the dynamics of a system as it transitions through states that are not directly accessible to the observer (Bronson et al., 2009). However, our observable (the MS2 signal) does not correspond to the hidden variable of interest (the promoter state) in a one-to-one fashion (compare Figure 4C top and bottom). Instead, the observable MS2 signal reflects the net effect of promoter switching over a period equal to the time that an RNA polymerase molecule takes to transcribe the whole gene. Thus, instantaneous fluorescence does not just depend on the current promoter state; it exhibits a dependence on how active the promoter has been over a preceding window of time, which effectively constitutes a memory for recent promoter states (Choubey et al., 2015; Xu et al., 2016; Corrigan et al., 2016; Choubey, 2018; Choubey et al., 2018). Classic HMM approaches cannot account for this kind of system memory.

In order to model the process of transcription and extract the kinetic parameters of promoter switching, we augmented classic HMMs to account for memory (details about implementation of the method are given in Appendix 3). Similar approaches were recently introduced to study transcriptional dynamics in cell culture and tissue samples (Suter et al., 2011; Molina et al., 2013; Zechner et al., 2014; Zoller et al., 2015; Hey et al., 2015; Bronstein et al., 2015; Corrigan et al., 2016; Featherstone et al., 2016). We used simulated data to establish that mHMM reliably extracts the kinetic parameters of transcriptional bursting from live-imaging data (Appendix 4), providing an ideal tool for dissecting the contributions from individual bursting parameters to observed patterns of transcriptional activity across space and time.

Before applying our model to real-time transcriptional data, we had to account for the rapid replication of the *D. melanogaster* genome at the beginning of each nuclear cycle (Rabinowitz, 1941; Shermoen et al., 2010), which leads to the presence of two distinct *eve* loci within each fluorescent spot (Figure 4D and Video 5). The first evidence of resolved chromatids appears as early as 8 minutes into nuclear cycle 14 (Appendix 5–Figure 1)—coincident with the average onset time of transcription (Figure 3–Figure Supplement 2B). Moreover, our analysis indicates that replication of the relevant portion of the genome likely occurs in all eve-expressing nuclei by no later than 10 minutes following mitosis (Appendix 5–Figure 2). Thus, we conclude that the vast majority of our data feature two distinct *eve* loci within each diffraction-limited transcription spot. Moreover, while the distance between sister loci varies over time (see, *e.g*. Figure 4D), they nonetheless stay in relatively close proximity to ensure their proper segregation from each other at the next mitosis (Senaratne et al., 2016) such that the fluorescent intensity signals extracted from our data reflect the integral over both loci (Figure 2–Figure Supplement 2). As a result, if we assume that each locus can be well-represented by a two-state model of transcriptional bursting, then an effective three-state model is needed to capture *eve* dynamics (Figure 4E). For ease of exposition, we present our main results in the context of an effective two-state model, in which, as detailed in Appendix 1, the system is considered to be in the ON state so long as *either* chromatid is bursting (Figure 4F). Note that none of our conclusions below are affected by this choice of an effective model as shown in Appendix 6 where we present full results for the three-state model.

A typical experimental trace for a nucleus in the core of the stripe is shown in Figure 5A, along with its best fit, which corresponds to the mHMM-inferred promoter trajectory in Figure 5B. Our ability to infer the instantaneous promoter state in individual nuclei throughout development is further illustrated in Figure 5C and Video 6. These data revealed that, as development progresses and the stripe sharpens, the *eve* promoter continuously fluctuates between the ON and OFF states on a time scale of approximately 1-2 minutes.

**Figure 5.**
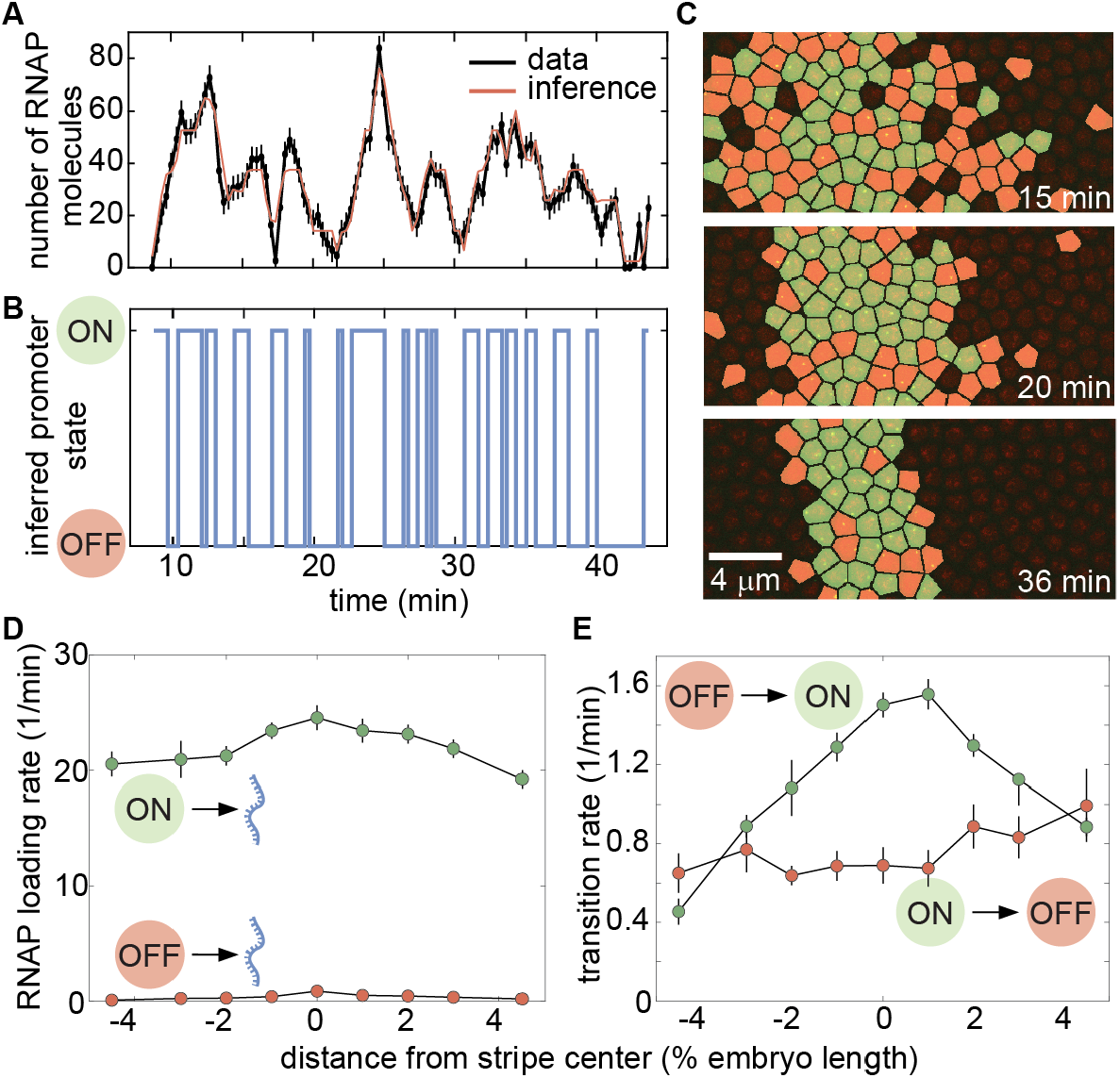
Inferring bursting dynamics using a memory-adjusted Hidden Markov model. **(A)** Representative experimental trace along with its best fit and **(B)** its most likely corresponding promoter state trajectory. **(C)** Instantaneous visualization of promoter state in individual cells throughout development through the false coloring of nuclei by promoter state (colors as in B). **(D)** The rate of initiation for each transcriptional state is not significantly modulated along the embryo. **(E)** Our mHMM reveals that the transition rate between the OFF and ON state (equivalent to the burst frequency) is up-regulated in the stripe center. (A, error bars obtained from estimation of background fluorescent fluctuations, as described in Materials and Methods and Garcia et al. (2013); D,E, error bars indicate the magnitude of the difference between the first and third quartiles of mHMM inference results for bootstrap samples of experimental data taken across 11 embryos. See Materials and Methods for details.) **Figure 5–Figure supplement 1. Fraction of time spent in each transcriptional state.**

In order to infer time-averaged bursting parameter values, we grouped traces by position along the anterior-posterior axis. The rate of RNA polymerase loading, *r*, remained constant throughout the stripe (Figure 5D), suggesting that none of the transcription factors regulating *eve* stripe 2 act on this kinetic parameter. Similarly, we noted no significant spatial modulation of the rate of switching out of the ON state, *k*_off_ (Figure 5E). In contrast, the rate of switching into the ON state (also known as burst frequency), *k*_on_, was strongly up-regulated in the stripe center (Figure 5E). These observations suggested that, in order to enact analog control of the mean transcription rate, transcription factors act primarily on the rate of promoter turning on, consistent with previous results both in embryos (Xu et al., 2015; Desponds et al., 2016; Fukaya et al., 2016) and in single cells (So et al., 2011; Senecal et al., 2014; Padovan-Merhar et al., 2015; Bartman et al., 2016). This regulatory modality increases the fraction of time that loci near the stripe center spend in the ON state (Figure 5–Figure Supplement 1 and Zoller et al. (2018)).

### Binary control of the transcriptional time window is independent of transcriptional bursting

Having determined that analog control of the mean transcriptional rate is realized by the modulation of the burst frequency, *k*_on_, we next sought to uncover the molecular mechanism by which the binary regulation of the transcriptional time window is implemented. In one possible scenario, the onset of transcriptional quiescence at the end of the transcriptional time window would reflect a fundamental change to the molecular character of the transcriptional locus such that the bursting framework no longer applies. For instance, repressing transcription factors could induce an irreversible change in the local chromatin landscape that precludes further activator-mediated bursting, effectively silencing transcription (Figure 6A, top; Allis et al. (2015)). Alternatively, if the rates of promoter switching vary in time, then the time window could be explained without invoking an extra silenced state that is mechanistically distinct from the processes driving transcriptional bursting. In this scenario, one or multiple promoter-switching rates would change over time in order to progressively reduce the frequency, intensity, and/or duration of transcriptional bursts, abolishing all activity at the locus and leading to the observed quiescence. Such modulation could be achieved by downregulating *k*_on_, downregulating *r*, and/or upregulating *k*_off_ (Figure 6A, bottom).

**Figure 6.**
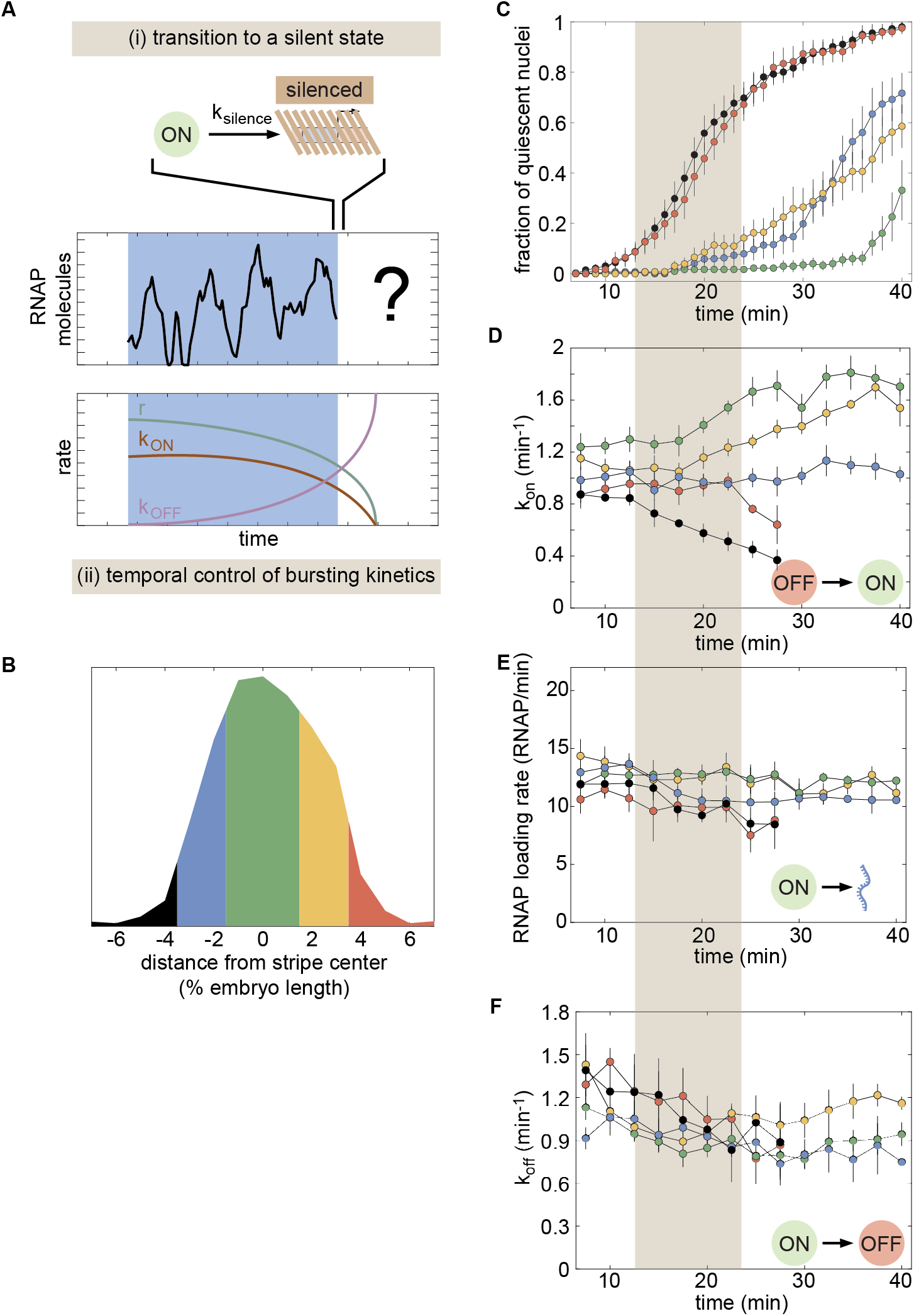
Investigating the molecular character of transcriptional quiescence. **(A)** Two hypotheses explaining promoter quiescence onset: (i) an irreversible transition into an alternative, transcriptionally silent state and (ii) the modulation of one or more bursting parameters over time. **(B)** Division of the stripe into five regions for our analysis of: **(C)** the fraction of quiescent nuclei, **(D)** the transition rate from OFF to ON, **(E)** the rate of RNA polymerase loading when the promoter is in the ON state, and **(F)** the transition rate from ON to OFF as a function of time and position along the stripe. Grey shaded region indicates the onset of transcriptional quiescence. (C, error bars indicate bootstrap estimate of the standard error of the mean. D-F, error bars indicate the magnitude of the difference between the first and third quartiles of mHMM inference results for bootstrapped samples of experimental data. See Materials and Methods for details)

In order to discriminate between these two possible scenarios, we split the stripe into the five regions shown in Figure 6B. For each region, we sought to determine whether the bursting dynamics varied over time in a manner that could explain the dynamics of entry into quiescence of individual nuclei (Figure 6C). To probe for this time-dependence in transcriptional bursting, we extended our mHMM method to obtain promoter-bursting parameters over discrete periods of time by performing inference on our live-imaging data using a sliding window (see Appendix 3 for details). Our inference revealed that the rate of promoter turn on, *k*_on_, varied significantly in time (Figure 6D). Specifically, *k*_on_ decreased in both the anterior and posterior stripe boundaries (black and red curves) as development progressed and the fraction of active nuclei decreased (grey shaded region), while loci in the stripe center (green and yellow curves) exhibited a significant *increase* in *k*_on_. Further, while relatively constant at most positions along the stripe, both the rate of RNA polymerase loading when in the ON state, *r*, and the rate of promoter turn off, *k*_off_, decreased slightly in (Figure 6E and F).

These findings confirmed our time-averaged inference results (Figure 5D and E) indicating that *k*_on_ was the primary kinetic pathway through which transcription factors influence *eve* stripe 2 transcription dynamics. Moreover, the coincidence of the decrease in *k*_on_ in flank nuclei with the onset of transcriptional quiescence (grey shaded region in Figure 6D) seemed to suggest that, at least in part, quiescence in the stripe flanks could be driven by the temporal modulation of bursting parameters (Figure 6A, bottom). However, other trends in our data results were not consistent with the view that a decrease in *k*_on_ drives transcriptional quiescence. Although 70% and 50% of nuclei in the regions directly anterior and posterior of the stripe center were quiescent by 40 min into the nuclear cycle (blue and yellow curves in Figure 6C), we detected no corresponding decrease in *k*_on_. In fact, *k*_on_ actually *increased* in some inner regions of the stripe (Figure 6D) —a trend that would increase overall transcriptional activity and would therefore go against the establishment of transcriptional quiescence.

The divergent outcomes observed in the central stripe regions, with the rate of transcriptional bursting remaining constant or increasing at *eve* loci within the engaged population of nuclei even as loci in neighboring nuclei turn off for good, runs counter to the hypothesis that quiescence is driven by the temporal modulation of the promoter switching parameters. It is conceivable that temporal changes in bursting parameters associated with the onset of quiescence occur too rapidly to be captured by our model. However, as discussed in Appendix 7, these changes would need to occur on the same time scale as bursting itself (1 to 3 min). Given that both the other temporal trends detected by our inference (Figure 6) and the shifts in the input transcription factors themselves (Appendix 8) unfold on significantly slower timescales (5-15 min), we concluded that while possible, a scenario were bursting dynamics are changing too quickly to detect is unlikely.

The contradictory trends observed in the stripe center suggested that entry into transcriptional quiescence might be akin to an irreversible transition into a silent state (Figure 6A, top), thus suggesting that binary control of the transcriptional time window and the transcriptional bursting driving the analog control of the mean transcription rate may arise from distinct molecular processes.

### Input-output analysis reveals distinct regulatory logic for bursting and the transcriptional time window

*eve* stripe 2 is mainly established by the combined action of two activators, Bicoid and Hunchback, and two repressors, Giant and Krüppel (Frasch and Levine, 1987; Small et al., 1992; Arnosti et al., 1996). If transcriptional bursting and the transcriptional time window are controlled by distinct molecular mechanisms, then distinct forms of regulatory logic may be at play. For example, the Bicoid and Hunchback activators could control transcriptional bursting, while the Giant and Krüppel repressors could dictate the entry into the quiescent state. In order to reveal the molecular logic controlling each regulatory strategy, we sought to correlate the fraction of nuclei that have entered the quiescent state (Figure 7A) and the fraction of nuclei in the bursting ON state (Figure 7B) with the corresponding spatiotemporal patterns in the input concentrations of these four transcription factors.

**Figure 7.**
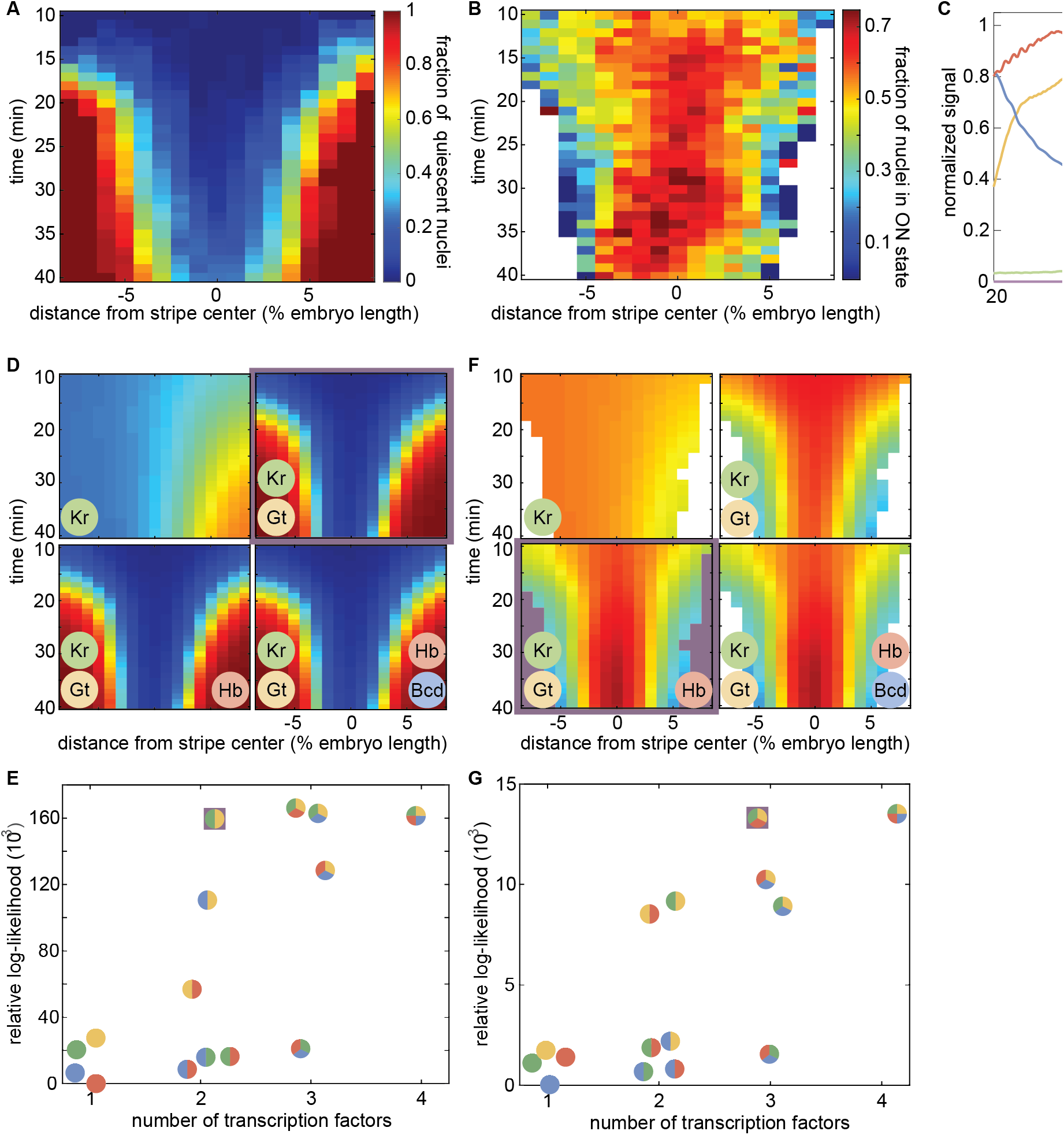
Probing the regulatory logic of bursting and the transcriptional time window. **(A)** Fraction of nuclei in the transcriptionally quiescent state and **(B)** fraction of nuclei in the bursting ON state as a function of time and position along the embryo. **(C)** Snapshot of input transcription factor levels and predicted *eve* mRNA levels of our “average” embryo. **(D)** Predicted fraction of quiescent nuclei for progressively more complex regression models. The simplest model with the highest likelihood is outlined in purple. **(E)** Model likelihood indicating that Krüppel and Giant levels are sufficient to recapitulate the fraction of quiescent nuclei in (D). **(F)** Predicted fraction of nuclei in the ON state. The simplest and most likely model is highlighted in purple. **(G)** Model scores reveal that Giant, Krüppel, and Hunchback recapitulate the bursting behavior in (F).

We measured Bicoid concentration profiles using a a well-established Bicoid-GFP fusion (Gregor et al., 2007) and obtained spatiotemporal concentration profiles for Krüppel, Giant, and Hunchback from published immunofluorescence data (Dubuis et al., 2013). We combined these data with our live-imaging data of *eve* stripe 2 transcriptional activity to generate an “average embryo” in which the concentration of all relevant inputs and the output transcriptional activity at each point in time and space were known (Figure 7C and Video 7). Building upon previous work (Ilsleyet al., 2013), we utilized logistic regressions to probe the regulatory role played by each of these four factors in the spatiotemporal control of transcriptional bursting and the transcriptional time window. Logistic regression is a widely used method of inferring predictive models in processes with binary outcomes. For example, in order to query the regulatory logic behind the control of the transcriptional time window, the model probes the impact of each transcription factor on the relative likelihood of a locus entering the quiescent state versus the likelihood of remaining transcriptionally engaged such that

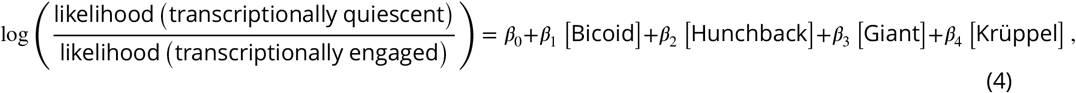

where the coefficients *β_n_* indicate the magnitude and nature (activating or repressing) of the transcription factor’s regulatory function. In estimating these coefficients, we used prior knowledge about the function of each transcription factor, requiring Bicoid and Hunchback to play activating roles, and Krüppel and Giant to play repressing roles (Small et al., 1991, 1992). We used an analogous model to investigate the regulatory logic controlling transcriptional bursting by inferring the factors that determine the relative likelihood that nuclei are in the bursting ON versus the OFF state, likelihood (ON state)/likelihood (OFF state).

Our analysis of the fraction of nuclei in the quiescent state revealed that no single transcription factor can explain quiescence dynamics (Figure 7D and E). However, a simple model in which increasing levels of the repressors Giant and Krüppel drive the onset of transcriptional quiescence in the anterior and posterior stripe flanks, respectively, recapitulated experimentally observed trends. The further addition of Hunchback and/or Bicoid had no impact on the model’s predictive power, suggesting that activator concentrations have no influence over the molecular processes responsible for silencing. Relaxing constraints on the functional role of each transcription factor-for instance, allowing the presumed activators to function as repressors-also provided no significant improvement over models presented here as shown in Appendix 8.

We next turned our attention to the relationship between transcription factor levels and the fraction of nuclei in the ON state (Figure 7B). Unlike the transcriptional time window, repressor levels alone could not recapitulate the observed bursting profile; Hunchback levels were also necessary in order to fully capture the spatiotemporal bursting dynamics (Figure 7E and G). Specifically, we linked a rise in Hunchback concentration to an observed rise in the fraction of nuclei in the ON state in the stripe center between 30 and 35 min into the nuclear cycle (Figure 7B and F).

Our input-output analysis thus revealed that bursting and the transcriptional time window exhibit significantly different forms of regulator logic: whereas repressor levels alone are sufficient to explain the transcriptional time window, the joint action of activators and repressors appears necessary to explain the observed patterns of transcriptional bursting. These results are consistent with the hypothesis that regulation of bursting and of the transcriptional time window occur via distinct molecular processes, therefore supporting a model in which the long-lived trancriptionally silent state observed in flank nuclei constitutes a distinct molecular state outside of the bursting model.

## Discussion

In *Drosophila* development, information encoded in a handful of maternally deposited protein gradients propagates through increasingly complex layers of interacting genes, culminating in the specification of the adult body plan. The prediction of this cascade of developmental outcomes requires a quantitative understanding of the mechanisms that facilitate the flow of information along the central dogma. Here, we utilized live imaging in conjunction with theoretical modelling to shed light on a critical link in this cascade: how the regulation of transcriptional activity at the single-nucleus level gives rise to a spatiotemporal pattern of cytoplasmic mRNA.

*A priori*, there are several distinct regulatory strategies at the single-cell level capable of generating spatially differentiated patterns of cytoplasmic mRNA (Figure 1), each with distinct implications for the nature of the underlying molecular processes at play. Several recent studies have revealed that the average rate of transcription is mainly modulated across the embryo by tuning the frequency of transcriptional bursting (Lionnet et al., 2011; Bothma et al., 2014; Xu et al., 2015; Desponds et al., 2016; Fukaya et al., 2016; Zoller et al., 2018). Yet it has remained unclear whether this modulation of the rate of transcription (and thereby mRNA production) is the dominant modality by which input concentrations of transcription factors drive the formation of patterns of gene expression, or if, instead, it is simply the most readily apparent mechanism among multiple distinct control strategies.

In this work, we derived a simple theoretical model that predicts how the interplay between regulatory strategies at the single-cell level dictates the accumulation of cytoplasmic mRNA and the subsequent formation of a gene expression pattern (Equation 2), and tested its predictions experimentally by performing single-cell live-imaging measurements in developing embryos using the MS2 system. Our results revealed that the modulation of the mean rate of transcription alone is insufficient to recapitulate the formation of a sharp gene expression stripe (Figure 3F, green). We discovered that the window of time over which promoters engage in transcription is sharply controlled along the axis of the embryo (Figure 3C and D) and that the joint action of the analog control of the rate of transcription and the binary control of the duration of transcription is necessary and sufficient to quantitatively recapitulate most of the the full stripe profile (Figure 3F, brown).

Here, we used biological numeracy as a driver for biological discovery: our discovery of the key role of the binary control of the transcriptional time window in pattern formation was only made possible by going beyond the *qualitative* description of pattern formation and demanding a *quantitative* agreement between our theoretical predictions and the experimental data (Phillips, 2015). Further, our work emphasizes how the regulation of gene expression *timing* in development is as important as the regulation of the spatial extent of these patterns along the embryo. Thus, in order to make progress toward a quantitative and predictive picture of pattern formation in development, it is necessary to go beyond the widespread steady-state, *static* picture of pattern formation in development put forward by previous single-cell transcriptional activity studies that focused on the study of snapshots of fixed embryos (Pare et al., 2009; Little et al., 2013; Xu et al., 2015; Zoller et al., 2018) and embrace a *dynamical* description that acknowledges that development is a process that occurs outside of steady state (Berrocal et al., 2018).

To determine whether the same molecular mechanisms dictate the analog control of the mean transcription rate and the binary control of the transcriptional time window, we utilized a variety of theoretical and computational tools in conjunction with our live-imaging data. Specifically, to uncover how the mean rate of transcription is regulated across the stripe, we developed a mHMM that is capable of inferring the instantaneous activity state of individual gene loci from MS2 traces. We used this mHMM to infer average promoter-switching parameters across the stripe (Figure 5). In agreement with previous measurements of various gene expression patterns (Xu et al., 2015; Desponds et al., 2016; Fukaya et al., 2016; Zoller et al., 2018), our results revealed that the the burst frequency (*k*_on_) is the main bursting parameter regulated by the input transcription factors across *eve* stripe 2. This increase in *k*_on_ in the stripe center increases the fraction of time that nuclei spend in the active transcriptional state.

Importantly, our mHMM algorithm is not limited to the *eve* stripe 2 system and should prove useful to infer the underlying regulatory kinetics of any gene that is tagged using approaches such as the MS2 or PP7 systems in any organism (Larson et al., 2011; Hocine et al., 2012; Fukaya et al., 2016). Further, the method could be used to infer the state of the ribosome as mRNA is being translated into protein in novel single-molecule *in vivo* translation assays (Morisaki et al., 2016; Wang et al., 2016; Yan et al., 2016; Wu et al., 2016). Thus, we envision that our method will be useful for the broader biophysical analysis of *in vivo* cellular processes at the single-molecule level.

Having identified *k*_on_ as the primary kinetic mode by which transcription factors modulate the mean rate of expression across *eve* stripe 2, we next sought to probe the relationship between bursting and the transcriptional time window (Figure 6A). We adapted our mHMM to go beyond time-independent models of promoter switching to infer the regulation of these rates across both space and time. We observed striking temporal trends indicating that the burst frequency responds dynamically to time-varying transcription factor inputs. However, we noted a significant disconnect between temporal trends in the burst frequency and the onset of transcriptional quiescence. In particular, *k*_on_ either increased or remained constant near the stripe center even as a significant fraction of *eve* nuclei transitioned into quiescence (Figure 6C and D). We reasoned that the onset of transcriptional quiescence is likely not the result of a progressive reduction in burst frequency, amplitude, or duration, and that quiescence is instead driven by molecular processes that are distinct from those that regulate transcriptional bursting such as transcriptional silencing.

To test this hypothesis, we utilized a logistic regression framework and time-resolved data for the primary regulators of *eve* stripe 2 to query the regulatory logic exhibited by the time window and bursting, respectively (Appendix 8). Consistent with our time-resolved mHMM results, the two regulatory strategies responded to transcription factor concentrations in different ways. On the one hand, increasing levels of Giant and Krüppel were sufficient to explain the onset of transcriptional quiescence in the stripe flanks (Figure 7A and D). This observation points to a model in which repressor levels act unilaterally—without respect to coincident levels of activator proteins—to shut off transcription at loci in an (at least effectively) irreversible fashion. Conversely, the joint action of Giant, Krüppel, and Hunchback was necessary to recapitulate the observed pattern of transcriptional bursting (Figure 7B and F).

This difference in the regulatory logic observed for the two strategies dissected in this work suggests that control of the transcriptional time window and the modulation of the average transcription rate arise from two distinct, orthogonal molecular mechanisms. Further, the striking absence of a direct functional role for Bicoid in the regulation of either phenomenon suggests that, while Bicoid is almost certainly necessary for the expression of *eve* stripe 2 (Small et al., 1992), it does not play a direct role in dictating the magnitude or duration of *eve* stripe 2 transcription. In this interpretation, Bicoid functions like a general transcription factor, facilitating the transcription of *eve* 2 without directly conferring spatiotemporal information.

While the results of our input-output analysis provide valuable insights into the mechanisms driving the regulation of transcription of the *eve* stripe 2 enhancer, key questions remain about the molecular character of the underlying processes. For instance, while loci engaged in transcriptional bursting appear to continuously sense changes in transcription factor concentrations, it remains an open question whether loci continue to actively read out transcription factor concentrations following the onset of transcriptional quiescence. While the transition *appears* irreversible in our data, it is possible that quiescence is, in fact, reversible in principle, but simply not observed in practice because repressor levels increase over time in our region of interest. In such a scenario, the direct action of relatively short-lived repressor binding could function to silence the locus, and a reduction in repressor concentration would lead to a rapid reactivation of transcription. Alternatively, if repressor levels function more akin to a trigger to, say, induce a change in the chromatin state at the *eve* locus, this would imply that loci, once quiescent, cease to sense transcription factor concentrations and would fail to reactivate even if repressor levels decreased. Of course, intermediate cases could also be imagined.

In order to further test these and other molecular hypotheses, it will be critical to move beyond spatiotemporal averages for transcription factor inputs (Figure 7C) and, instead, use live singlenucleus measurements to directly correlate input transcription factor concentration dynamics with the corresponding transcriptional activity at the single-cell level (Holloway and Spirov, 2017). Experimentally, we recently demonstrated the simultaneous measurement of inputs and outputs in single nuclei of a living fly embryo using novel genetically encoded LlamaTags (Bothma et al., 2018). We believe that utilizing this novel technique, in conjunction with the theoretical methods presented here, to query the effects of targeted disruptions to transcription factor binding domains on regulatory enhancers will constitute a powerful assay for querying transcription factor function at the molecular level. Thus, there are clear experimental and theoretical paths to uncovering the detailed quantitative mechanisms behind the molecular control of transcriptional bursting and quiescence in development. Such a quantitative description is a necessary step toward a predictive understanding of developmental decision-making that makes it possible to calculate developmental outcomes from knowledge of the nature of the transcription factor interactions within gene regulatory networks.

## Materials and methods

### Cloning and transgenesis

This work employed the same *eve* stripe 2 reporter construct developed by Bothma et al. (2014). This construct contains the *even-skipped* (*eve*) stripe 2 enhancer and promoter region (spanning −1.7 kbp to +50 bp) upstream of the *yellow* reporter gene. 24 repeats of the MS2 stem loop sequence were incorporated into the 5’ end of the reporter gene.

### Sample preparation and data collection

Sample preparation followed procedures described in Bothma et al. (2014) and Garcia and Gregor (2018). In short, female virgins of yw;His-RFP;MCP-GFP (MCP, MS2 coat protein) were crossed to males bearing the reporter gene. Embryos were collected and mounted in halocarbon oil 27 between a semipermeable membrane (Lumox film, Starstedt) and a coverslip. Data collection was performed using a Leica SP8 Laser Scanning Confocal Microscope. Average laser power on the specimen (measured at the output of a 10x objective) was 35 *μ*W. Image resolution was 256 × 512 pixels, with a pixel size of 212 nm and a pixel dwell time of 1.2 *μ*s. The signal from each frame was accumulated over three repetitions. At each time point, a stack of 21 images separated by 500 nm were collected. Image stacks were collected at a time resolution of 21 seconds. The MCP-GFP and Histone-RFP were excited with a laser wavelength of 488 and 556 nm using a White Light Laser, respectively. Fluorescence was detected with two separate Hybrid Detectors (HyD) using the 498-546 nm and 566-669 nm spectral windows. Specimens were imaged for a minimum of 40 minutes into nuclear cleavage cycle 14.

### Image analysis

Image analysis of live embryo movies was performed based on the protocol found in Garcia et al. (2013) with modifications to the identification of transcriptional spots, which were segmented using the Trainable Weka Segmentation plugin for FIJI using the FastRandomForest algorithm (Schindelin et al., 2012; Schneider et al., 2012; Arganda-Carreras et al., 2017; Witten et al., 2016). In comparison with a previous algorithm based on Difference of Gaussians (Little et al., 2013; Garcia et al., 2013; Bothma et al., 2014, 2015), this alternative spot segmentation approach was found to be superior for the detection of dim transcription spots—a feature critical to establishing the precise timing of the cessation of activity at transcriptional loci.

### Data processing

Processed live-imaging movies were compiled from across 11 experiments (embryos) to form one master analysis set. While the position of *eve* stripe 2 along the anterior-posterior axis of the embryo was found to be consistent to within 1-2% of egg length, we sought to further reduce this embryo-to-embryo variation by defining new, “registered” AP axes for each experiment using the observed position and orientation of the mature stripe. To this end, an automated routine was developed to consistently establish the position and orientation of the *eve* stripe 2 center for each data set.

This routine, described graphically in Figure 2–Figure Supplement 1, used observed spatial patterns of fluorescence measured from 30 minutes into nuclear cycle 14—the approximate time at which the mature stripe is first established (Bothma et al., 2014)— to the time of last observation (≥40 min) to find the natural position and orientation of the mature stripe. Generally, the *eve* stripes run roughly perpendicular to the anterior-posterior (AP) axis of the embryo; however, the approach allowed for the possibility that the true orientation of the *eve* 2 stripe deviated from the orientation implied by manual estimates of the anterior posterior axis. Thus, a variety of orientations for the natural stripe axis were considered, ranging between ± 15 degrees from the line perpendicular to the stripe with the manually specified anterior posterior axis. For each orientation, a sliding window of 4% embryo length in width was used to find the position along the proposed orientation that captured the largest fraction of the total fluorescence emitted by the mature stripe. The orientation and position that maximized the amount of fluorescence captured within this window defined a line through the field of view that was taken as the stripe center. All anterior-posterior positions used for subsequent analyses were defined relative to this center line.

Once the stripe centers for each set were established, fluorescence traces were interpolated to 20s resolution, with all times shifted to lie upon a common reference time grid. Traces near the edge of the field of view or that exhibited uncharacteristically large changes in fluorescence over a time step were flagged through a variety of automated and manual filtering steps. When necessary, these traces were removed from subsequent analyses to guard against the influence of non-biological artifacts.

### mHMM inference

To account for finite RNA polymerase elongation times, a compound state Markov formalism was developed in which the underlying tw-promoter system—assumed to have three states (see Figure 4E,F)—was transformed into a system with 3^w^ compound gene states, where *w* indicates the number of time steps needed for an RNA polymerase molecule to traverse the full transcript (see Appendix 9). These compound gene states played the role of the “hidden” states within the traditional HMM formalism. See Appendix 3 for details regarding the model’s architecture. Following this transformation from promoter states to compound gene states, it was possible to employ a standard version of the expectation-maximization (EM) algorithm, implemented using custom-written scripts in Matlab, to estimate bursting parameters from subsets of experimental traces (Appendix 3). The scripts are available at the GarciaLab/mHMM GitHub repository. Bootstrap sampling was used to estimate the standard error in our parameter estimates. Subsets of 8,000 data points were used to generate time-averaged parameter estimates. In order to accurately capture the time-averaged dynamics across the entirety of nuclear cycle 14, the full length of each experimental trace was used for time averaged inference. Sample sizes for windowed inference varied due to data set limitations. When possible, samples of 4,000 points were used. Only data points falling within a 15 minute window centered about the time point of interest were included in windowed inference runs. Inference was not conducted for spatiotemporal regions for which fewer than 1,250 time points were available. A minimum of 10 bootstrap samples were used to estimate each parameter value reported in this work. Reported values represent the median taken across bootstrap samples.

### Input-output logistic regressions

The input-output analysis presented in Figure 7 utilized input transcription factor data from immunostaining experiments presented in Dubuis et al. (2013), as well as live measurements of a Bicoid-GFP fusion courtesy of Jonathan Liu and Elizabeth Eck. Logistic regression parameters were estimated in Matlab using the *fmincon* function. See Appendix 8 for further details.

### Bootstrap error calculation

Bootstrap resampling was used frequently throughout this work to estimate the standard error in a variety of reported quantities, from trends estimated directly from raw experimental data in Figure 1 to mHMM inference results presented in Figure 5 and Figure 6. In this procedure, multiple bootstrap replicates, 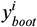 are generated by sampling with replacement from the pool of available experimental data, *Y* (see, *e.g*. Efron and Hastie (2016)). The parameter of interest (say, *t*_on_(*x*)) is then calculated for each replicate and the mean of these estimates is taken as the bootstrap estimate of the parameter value, 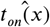, while the standard deviation across the pool of bootstrap parameter estimates is used to approximate the standard error in our estimate of *t_on_*(*x*). In our case, simply performing this procedure across the available pool of nuclei failed to account for biological variability that exists from embryo to embryo. To account for this, we introduced a hierarchical bootstrapping procedure. The first step in this procedure was to draw bootstrap samples from across the 11 embryos used in this study. Because these samples were taken with replacement, most bootstrap samples excluded some embryos out of the original set of 11 and included duplicates (or triplicates, etc.) of others. Each embryo-level bootstrap defined a subset of nuclei. The final set of nuclei used for parameter estimation was generated by performing another round of bootstrap sampling on this pool. Bootstrap averages and standard errors were then calculated as described above. This two-step procedure thus accounts for both embryo-to-embryo and nucleus-to-nucleus variability.

We note that the limited number of data points available for many spatiotemporal regions prevented us from performing this two-tiered bootstrap procedure in the case of our time-dependent mHMM inference (Figure 6D-F and Appendix 6–Figure 3D-E). In these cases, we used all available sets (essentially skipping the first bootstrap resampling step) and took bootstrap samples from amongst available nuclei as in step two of the procedure described above.

### Absolute calibration of MS2 signal

In order to frame our results with respect to units with a clear physical interpretation, we calibrated our fluorescence measurements in terms of absolute numbers of mRNA molecules. This calibration was also used to inform our Poisson loading sensitivities (Appendix 3). To calculate this calibration for our *eve* stripe 2 data, we relied on measurements reported by a previous study that utilized MS2 in conjunction with single molecule FISH to establish a calibration factor, *α*, between the integrated MS2 signal, *F*_MS2_, and the number of mRNA molecules produced at a single transcriptional locus, *N*_FISH_, (Garcia et al., 2013) given by

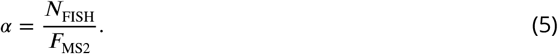

This calibration factor can be used to estimate the average contribution of a single mRNA molecule to the observed (instantaneous) fluorescent signal. While the values for the parameters in Equation 5 reported here pertain to the transcriptional output driven by the Bicoid activated P2 enhancer and promoter during nuclear cycle 13, the calibration should generalize to all measurements taken using the same microscope.

First, consider the total integrated fluorescence emitted by a single nascent mRNA while it is on the reporter gene,

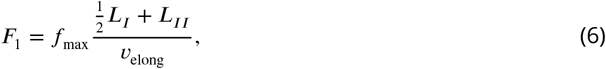

where *f*_max_ denotes the instantaneous fluorescence emitted by a nascent mRNA that has transcribed the full complement of MS2 loops, *L_I_* indicates the length of the MS2 loops, *L_II_* indicates the distance between the end of the MS2 loop cassette and the 3’ end of the gene, and *υ*_elong_ indicates the elongation rate of RNA polymerase molecules along the gene. We can solve for *f*_max_ using *α*, namely,

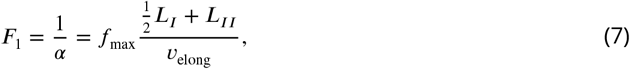

such that

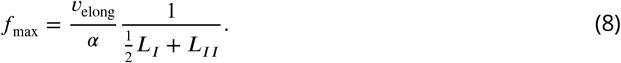

Here, we recognize that the cumulative fluorescence per RNA polymerase molecule is simply the inverse of the number of molecules per unit fluorescence (*α*). Now we have the pieces necessary to derive an expression for the *instantaneous* fluorescence of a single RNA polymerase molecule, that is,

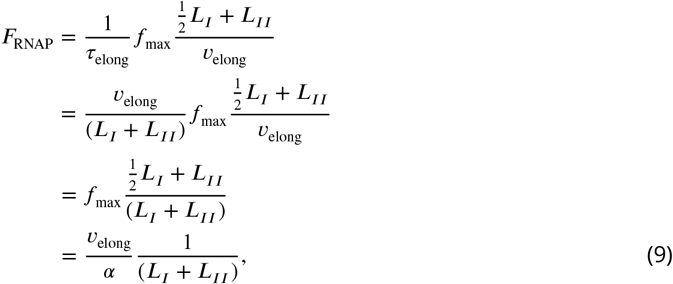

resulting in

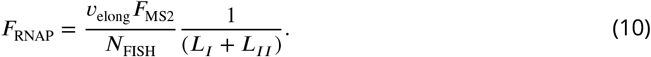

Measurements performed in Garcia et al. (2013) give *N*_FISH_ to be 220 (± 30) mRNA per nucleus and *υ*_elong_ to be 1.5 (± 0.14) kb/min. Experimental measurements on the P2 enhancer (courtesy of Elizabeth Eck, Maryam Kazemzadeh-Atoufi and Jonathan Liu) indicate that the total fluorescence per nucleus, *F*_MS2_, is 9,600 (±320) AU minutes. For the reporter gene used to take these measurements, *L_I_* and *L_II_* are 1.275 kb and 4.021 kb, respectively. Thus, we obtain

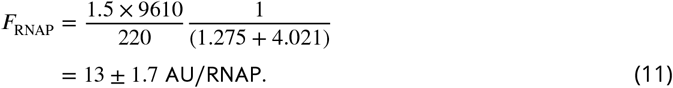

Though the error in our calibration is significant (>13%), the conversion from arbitrary units to numbers of nascent mRNA nonetheless provides useful intuition for the implications of our inference results, and none of our core results depend upon having access to a precise calibration of the observed signal in terms of absolute numbers of RNA polymerase molecules.

## Videos

**Video 1. Transcriptional activity of** *eve* **stripe 2 reported by MS2**. Raw MS2 signal where fluorescent puncta report on the number of actively transcribing RNA polymerase molecules.

**Video 2. Mean rate of transcription of** *eve* **stripe 2 reported by MS2**. Nuclei false colored by their mean transcriptional activity averaged over a 4 min time window as a function of time.

**Video 3. Transcriptional time window**. Nuclei along the stripe false colored after the duration of their transcriptional time window.

**Video 4. Fraction of active nuclei**. Nuclei along the stripe false colored according to whether they engaged in transcription at any time point during the nuclear cycle.

**Video 5. Fluorescent puncta contain sister chromatids**. Fluorescent puncta transiently separate to reveal the presence of sister chromatids as shown by the white circles throughout the movie.

**Video 6. Real-time inferred promoter states**. Real-time inference of effective promoter ON (green) and OFF (red) state in individual nuclei.

**Video 7. Average embryo containing all inputs and the output**. Average concentrations of Bicoid (blue), Hunchback (red), Krüppel (green) and Giant (yellow) combined with the average transcriptional activity of the *eve* reporter (purple). (Hunchback, Krüppel and Giant data obtained from Dubuis et al. (2013)).

## Acknowledgments

We thank Thomas Gregor and Lev Barinov for discussion about an initial implementation of the mHMM approach, Florian Jug for help with the spot segmentation using machine learning, and Elizabeth Eck, Maryam Kazemzadeh-Atoufi and Jonathan Liu for the P2 data used in the absolute MS2 calibration and for the Bicoid-GFP data used in the logistic regression. We are also grateful to Jack Bateman, Jane Kondev, Rob Phillips, Allyson Sgro, Donald Rio and Richard Harland for comments and discussion on the manuscript. HGG was supported by the Burroughs Wellcome Fund Career Award at the Scientific Interface, the Sloan Research Foundation, the Human Frontiers Science Program, the Searle Scholars Program, the Shurl & Kay Curci Foundation, the Hellman Foundation, the NIH Director’s New Innovator Award (DP2 OD024541-01), and an NSF CAREER Award (1652236). NL was supported by NIH Genomics and Computational Biology training grant 5T32HG000047-18. CW was supported by the NIH/NIC (U54 CA193313), CUNY (RFCUNY 40D14-A), and the NSF (IIS-1344668).

## Appendix 1

### Theoretical model to predict cytoplasmic mRNA levels given from *in vivo* measurements of transcriptional activity

#### Derivation details

Here we provide a more detailed treatment of the mathematical framework for connecting transcriptional activity in individual nuclei to levels of accumulated cytoplasmic mRNA. We begin with general expressions for the rate of mRNA production during the transcriptionally active and quiescent periods that dictate the transcriptional time window. When the promoter is actively transcribing (*t*_on_ ≤ *t* ≤ *t*_off_), the net rate of mRNA production is

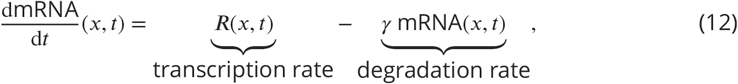

where *γ* is the mRNA degradation rate constant. For a promoter that has entered a transcriptionally quiescent state (*t* > *t*_off_), we have

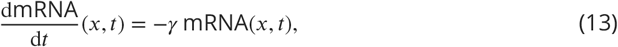

such that degradation is now the only contribution to the change of mRNA concentration in time. Note that, in these two equations, we have ignored the contribution of mRNA diffusion. Previous measurements have estimated a diffusion coefficient of mRNA of 0.09 *μ*m^2^/s (Halstead et al., 2015) and a typical mRNA degradation rate of 0.14 min^−1^ (Edgar et al., 1987). Given these numbers, we expect an *eve* mRNA molecule to diffuse approximately 6 μm, which corresponds to one nuclear diameter or 1% of the embryo length, before being degraded. Thus, given the overall width of the stripe mRNA profile of about 8% of the embryo length (Figure 3G), we expect diffusion to play a minimal role in stripe formation. Finally, note that we are also ignoring the delay between transcriptional initiation and the delivery of an mRNA molecule to the cytoplasm as a result of nuclear export. This delay would affect the *timing* of pattern formation, but would leave our conclusions about the relative role of transcriptional bursting and the regulation of the duration of the transcriptional time window unaffected.

To make progress, as in the main text, we make the simplifying assumption that the instantaneous rate of transcription can be well approximated by the time average at each position given by

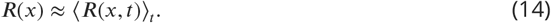

We now consider the role of *t*_on_(*x*) in dictating pattern formation by envisioning a scenario where transcription begins at time *t*_on_(*x*), but does not cease. In this scenario, the accumulated mRNA is given by

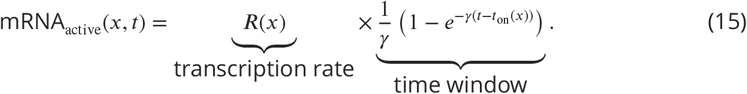

Note that if the system evolves for a long amount of time, the second term in the parenthetical in Equation 15 becomes vanishingly small (*γ*(*t* – *t*_on_(*x*)) ≫ 1) such that all time dependence drops out of the expression and we recover the familiar expression for mRNA levels in steady state

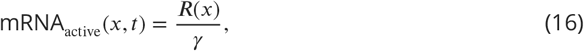

where mRNA production and degradation are balanced.

Next, consider the impact of regulating the timing with which nuclei cease transcriptional activity and become quiescent, *t*_off_. Here, when *t* > *t*_off_(*x*), the amount of mRNA produced during the period of activity is subsumed within a decaying exponential envelope such that

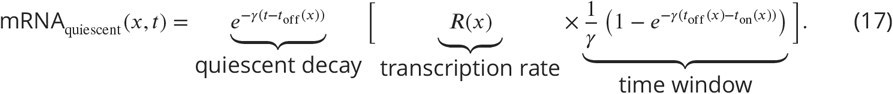

Equation 17 represents a scenario in which the accumulation of cytoplasmic mRNA results from the interplay between two distinct regulatory strategies: the modulation of when the transcription starts and stops (binary control of the transcription time window) and the average rate with which transcription occurs within this time window (analog control of transcriptional bursting). We refactor Equation 17 to reflect this distinction and consider the case when *t* > *t*_on_, giving

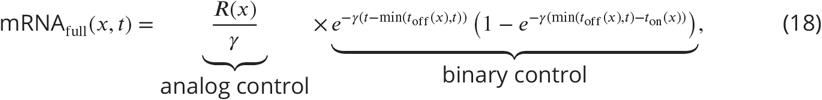

which can be simplified slightly to yield

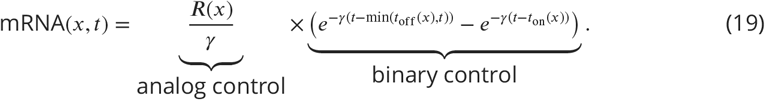

Finally, we account for the fact that only some *p*_active_(*x*) fraction of nuclei within each region *ever* engage in transcription leading to

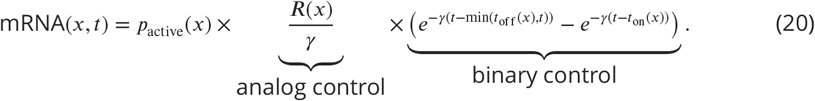

This equation constitutes the basis of our theoretical dissection of pattern formation by transcriptional bursting and the control of the transcriptional time window.

#### Accounting for multiple transcriptional states

In the main text, Equation 3 expresses the mean rate of mRNA production, *R*(*x*), as a function of the bursting parameters *k*_on_, *k*_off_, and *r*. We can combine this equation with Equation 20 to obtain an expression for the predicted amount of cytoplasmic mRNA that includes the burst parameters inferred by our mHMM

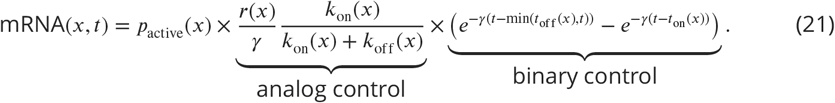

While we present our results in terms of an effective two-state model in the main text, the presence of two transcriptional loci within each observed fluorescent spot suggests that the system is more naturally described using a three-state kinetic model. Here, we extend the framework presented in Equation 21 to a scenario in which there are three distinct system states: 0 promoters on (0), 1 promoter on (1), and both promoters on (2) (see Figure 4). We begin with a general expression for this scenario that takes the contribution from the analog control term shown in Equation 20 to be a sum over the output of each of the 3 activity states, namely,

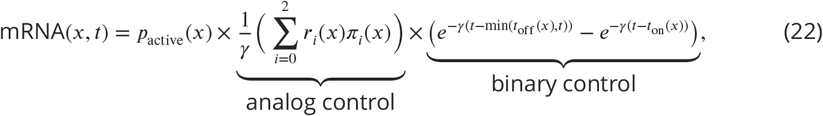

where *r_i_*(*x*) is the rate of RNA polymerase loading for state *i*, and *π_i_*(*x*) indicates the fraction of time spent in state *i*. Note that the independent effect of the duration of the transcription time window and of mRNA decay on cytoplasmic mRNA levels remain unchanged in the multi-state case.

The fractional occupancies of the activity states (*π_i_*(*x*) terms in Equation 22) are a function of the rates with which the promoter switches between activity states. In general, the fractional occupancy of each activity state, *π_i_*, may vary as a function of time; however we focus on their steady state values here, such that:

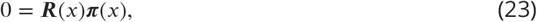

where *R*(*x*) is the transition rate matrix. Consistent with our inference results, we assume that no transitions are permitted between the high and low states (0 & 2). Thus, the transition rate matrix takes the following form:

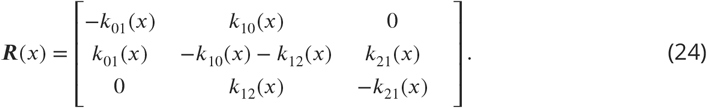

Together, Equation 23 and Equation 24 allow us to solve for the fractional occupancy of each activity state as a function of the transition rates that describe the system.

For the remainder of this derivation, we will drop the explicit *x* and *t* dependencies for ease of notation. Intuitively, the steady state (or stationary) distribution represents a limiting behavior of the system such that, upon reaching ***π***, no further shifts occur in the mean fraction of time spent in each activity state. Equation 23 leads to a system of three equations:

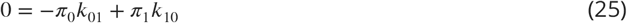

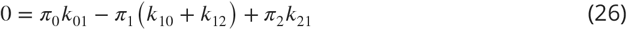

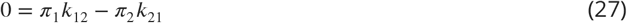

Before proceeding, we note that, since ***π*** is a probability distribution, we can eliminate one of our unknowns by enforcing normalization, that is,

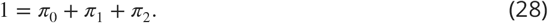

With this in mind, we can solve Equation 25 for *π*_1_ to find

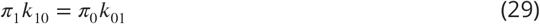

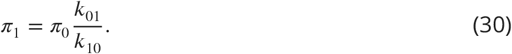

Next, we use the normalization condition to eliminate *π*_2_ from Equation 27:

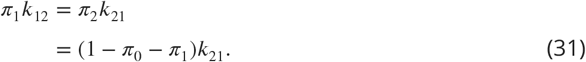

By combining this result with Equation 30, we obtain

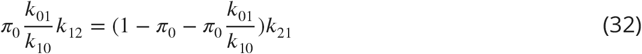

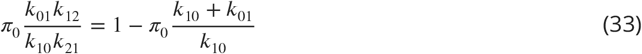

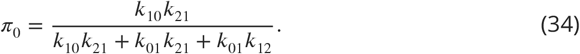

With Equation 34 in hand, it is then straightforward to solve for the remaining *π_i_* terms. First we obtain *π*_1_ by substituting Equation 34 into Equation 30:

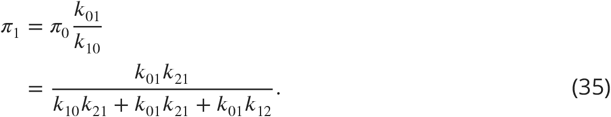

And finally *π*_2_:

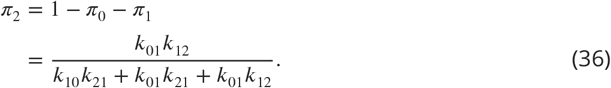

Thus, we arrived at the full expression for cytoplasmic mRNA levels in the 3-state case:

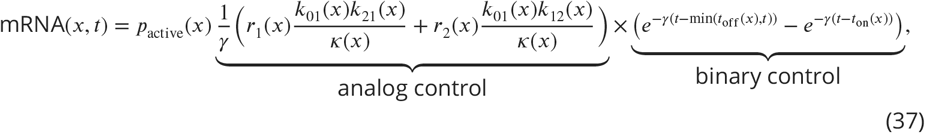

where, consistent with the 2-state case, we have taken *r*_0_(*x*) to be equal to zero and where *κ*(*x*) denotes the denominator in Equation 34, Equation 35 and Equation 36, namely,

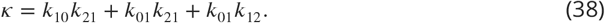

Thus, from Equation 37 we see that, while there are more terms comprising the analog control expression, the expression nonetheless takes on the same essential form as in Equation 20.

#### Mapping the three-state model into an effective two-state model

Here we provide expressions relating the effective two-state parameters presented in the main text to parameters from the full three-state model. As we have done throughout this work, we take the transition rates between states (0) and (2) of the 3-state model to be negligible (consistent with inference results, see Appendix 6). First, the on rate, 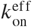 is directly equivalent to the transition rate between states (0) and (1), that is,

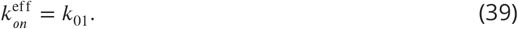

Similarly, since we do not observe from state (2) to state (0), 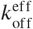 is equal to the transition rate from (1) to (0), weighted by the relative fraction of time the system spends in state (1) when it is in the effective ON state (1 or 2). Thus, we have:

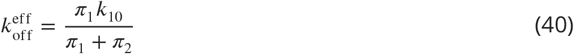

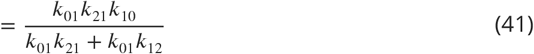

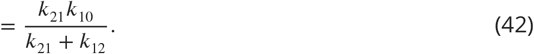

Finally, *r*^eff^ is the occupancy-weighted average of the initiation rates for states (1) and (2)

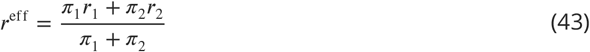

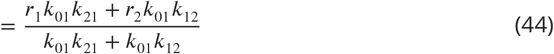

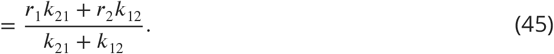

## Appendix 2

### Measuring the amount of produced mRNA

Here, we outline the approach that was used to estimate the total amount of mRNA produced by *eve* stripe 2 nuclei from MS2 traces. This approach, which is independent of the bursting parameter estimates returned by mHMM, was used to calculate the total cytoplasmic mRNA levels per nucleus shown in Figure 3G (red), as well as the “binary control” of the duration of the transcriptional time window contribution Figure 3G (blue).

#### Calculating full mRNA profiles

The observed fluorescent signal at transcriptional loci as a function of time, *F*(*t*), is linearly related to the number of actively transcribing RNA polymerase molecules. Thus, after a period equal to the amount of time needed for an RNA polymerase molecule to transcribe the gene, *τ*_elong_, the number of new mRNAs added to the cytoplasm will be proportional to *F*(*t*) (Bothma et al., 2014), that is,

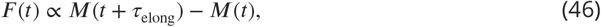

where *M*(*t*) indicates the total number of mRNA molecules that have been produced up to time *t*. We relate this fluorescence signal to absolute numbers of RNA polymerase molecules using the calibration procedure described in the Materials and Methods. However, only the *relative* amounts of mRNA present across the *eve* stripe 2 pattern are needed in order to calculate the relative contributions from the different regulatory strategies identified in Equation 2. Thus, we capture the calibration factor, along with all other proportionality constants, with a generic term *β*, with the expectation that *β* will drop out from all consequential stripe contribution calculations. Drawing from the derivation provided in the SI Methods of Bothma et al. (2014), we take the rate of mRNA production at time *t* to be approximately equal to the observed fluorescence at time 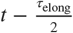

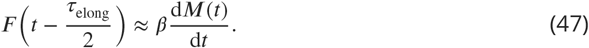

Here, the 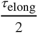 term accounts for the time lag between the number of transcribing nascent mRNA and the rate of mRNA release into the cytoplasm. For ease of notation, we will ignore this offset factor for the remainder of this section. We will also treat the relationship in Equation 47 as one of equality. For Figure 3G, the metric of interest is the amount of mRNA produced *per nucleus*. Thus for a given region along the axis of the embryo, the average observed fluorescence across *all N* nuclei (active, quiescent, and those that never engaged in transcription) within the region of interest was used as a proxy for the instantaneous rate of mRNA production per nucleus, given by

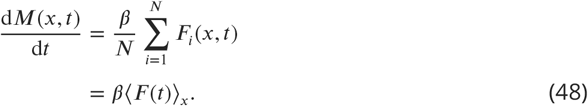

Here, *F_i_*(*x*,*t*) is the fluorescence of nucleus *i* at time *t*. The *x* subscript in Equation 48 indicates that the average is taken over all nuclei falling within the same anterior-posterior region within the *eve* stripe 2 pattern. Having obtained an expression for the rate of mRNA production as a function of space and time, we next sought to account for the degradation of mRNA over time. As indicated in the main text, we assumed a constant rate of mRNA decay, *γ*, over space and time. The next section in this appendix provides evidence for the validity of this assumption. For a constant mRNA decay rate, calculating the average concentration of mRNA amounts to taking a weighted sum over all preceding production rates for a position of interest, where the weight terms account for the effects of mRNA decay and are of the form *e*^−*γt*)^. Thus, we summed over all time points for each region of interest to estimate the total amount of cytoplasmic mRNA present on average, yielding the quantity on the left-hand side of Equation 2, namely,

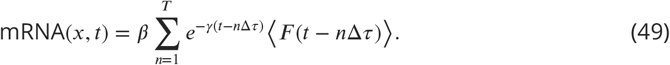

Here Δ*τ* is the experimental time resolution, and 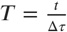 denotes the number of measurements taken through time *t*. The exponential term within the summand on the right-hand side captures the effects of mRNA decay (see Appendix 1). Finally, to calculate the normalized mRNA profile shown in Figure 3G (red), the estimates for the total accumulated mRNA per nucleus found using Equation 49 must be divided by the sum across all spatial regions considered

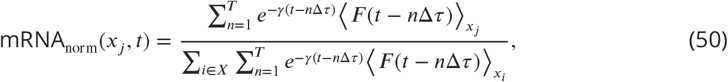

where the subscripts *i* and *j* outside the angled brackets denote the spatial region over which the sum is taken. Note that the proportionality constant *β* cancels in the final expression for mRNA_norm_. As a final step, we subtract the minimum across the AP region considered to remove any basal offset such that

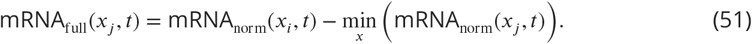

#### Calculating mRNA profiles due to the binary control of the transcriptional time window

The predicted profile due to binary control of the transcriptional time window alone (Figure 3G, blue) was calculated following the same procedure as for the full mRNA profile described above, save for the fact that, in this case, instantaneous fluorescent values for individual nuclei were converted to binary indicator variables (*f_i_*(*t*)) that were set equal to 1 if 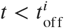 and 0 otherwise. Additionally, only nuclei that were active at *some point* during nuclear cycle 14 were included, to distinguish the effects of the transcriptional time window (Figure 1C) from the control of the fraction of active nuclei (Figure 1D). Thus, in this scenario, the “average rate” of mRNA production is equivalent to the fraction of nuclei engaged in transcriptional activity at a given point in time such that the rate of mRNA production is given by

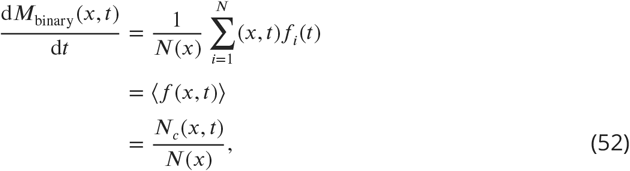

where *N_c_*(*t*) indicates the number of transcriptionally competent nuclei at time *t*. The binary equivalent to Equation 49 takes the form of a time-weighted sum of the fraction of active nuclei within a region

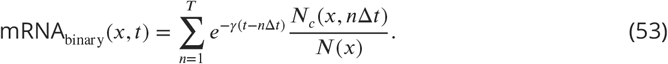

The steps for calculating the the normalized binary mRNA levels comprising the blue profile in Figure 3G from Equation 53 are identical to those shown for the full mRNA profile in Equation 50 and Equation 51 and are therefore not repeated here.

#### Comparison between predicted and measured cytoplasmic mRNA profiles

As a check for the validity of our approach to predicting levels of cytoplasmic mRNA from live imaging data (Equation 50 and Equation 51), we sought to compare our model’s predictions to existing mRNA FISH data for the endogenous *eve* stripe 2 (Fowlkes et al., 2008). For this comparison, we elected to use live imaging data for *eve* stripe 2 activity that was driven by a BAC containing the full *eve* locus (see Berrocal et al. (2018) for details). This was done to minimize potential differences with the activity of the endogenous gene. Most notably, the reporter construct used for the majority of this work does not contain an enhancer sequence that is responsible for driving *eve* expression late in nuclear cycle 14 Jiang et al., 1991).

Appendix 2 Figure 1 summarizes the results of this comparison. To account for uncertainty regarding the precise dorsal-ventral (DV) orientation of embryos within our live-imaging set, we compared our model’s predictions to mRNA measurements for a range of DV positions, encompassed by the green-shaded profile. We found a high degree of agreement between model predictions and reported levels of cytoplasmic mRNA. This conclusion is relatively insensitive to our assumptions regarding the average lifetime of *eve* mRNA as shown by the lines in the figure. While substantial uncertainties regarding the precise timing of the mRNA measurements prevented us from leveraging this comparison to, for instance, infer the rate of *eve* mRNA decay, we nonetheless concluded that it is sufficient to establish our modelling assumptions. In particular, the relative insensitivity of the distribution of cytoplasmic mRNA to the decay rate suggests that, while it is possible that the precise rate of mRNA decay is regulated across space or time, such phenomena—if they exist—would not impact the core conclusions presented in this work. Moreover, as discussed below, this paper’s findings are also relatively insensitive to our choice of decay rate *γ*, with the basic dynamics of stripe formation remaining consistent even in the limits near instantaneous and infinitely slow mRNA decay (Figure 2).

As alluded to above, several variables limited our ability to carry out a precise comparison between our model predictions and empirical measurements. The most significant of these was the lack of sufficiently precise temporal information for the empirical mRNA measurements. The authors used the percent invagination of cellular membranes through cellularization as a means to break individual embryos into rough temporal cohorts (Luengo Hendriks et al., 2006). We cross-referenced the invagination ranges for each temporal group with data provided by Dubuis et al. (2013) to obtain estimates for the range of times encapsulated by each of these cohorts. This calibration revealed that most time-averaged cohorts spanned too broad a range of times to allow for reasonable comparison. We elected to use the cohort comprised of embryos with ages ranging between 38 to 48 minutes into nuclear cycle 14 both because this range was much narrower than that spanned by the preceding cohort and because we had established that the stripe appeared to be relatively stable during this time period. An additional complication with establishing the precise timing of each cohort was the fact that the authors of Luengo Hendriks et al. (2006) measured invagination on the ventral surface of the embryo, while the authors in Dubuis et al. (2013) used the dorsal surface. However, while invagination is known to proceed more rapidly on the ventral side of the embryo,the authors in Luengo Hendriks et al. (2006) reported that this discrepancy is minimal up to the point where cell membrane extension has progressed to approximately 40% of its eventual full extent. The lower and upper bounds on the percent membrane invagination for the chosen cohort are 26% and 50% respectively. Thus, we expect the time estimate derived for the beginning of the period to be reasonably accurate, since Dorsal and Ventral membrane progression was reported to be comparable during this period. On the other hand, to the degree that Ventral invagination outpaces Dorsal invagination at the end of our period of interest, this would result in an over-estimation of ending time. Thus, it is possible that the true temporal window encompassed by the selected cohort is actually tighter than 10 minutes, since the ending time might in fact be earlier than 48 minutes into nuclear cycle 14. Given the relative stability of the stripe profile during this period of development, we do not expect this potential discrepancy to have a material impact on our conclusions.

**Appendix 2 Figure 1.**
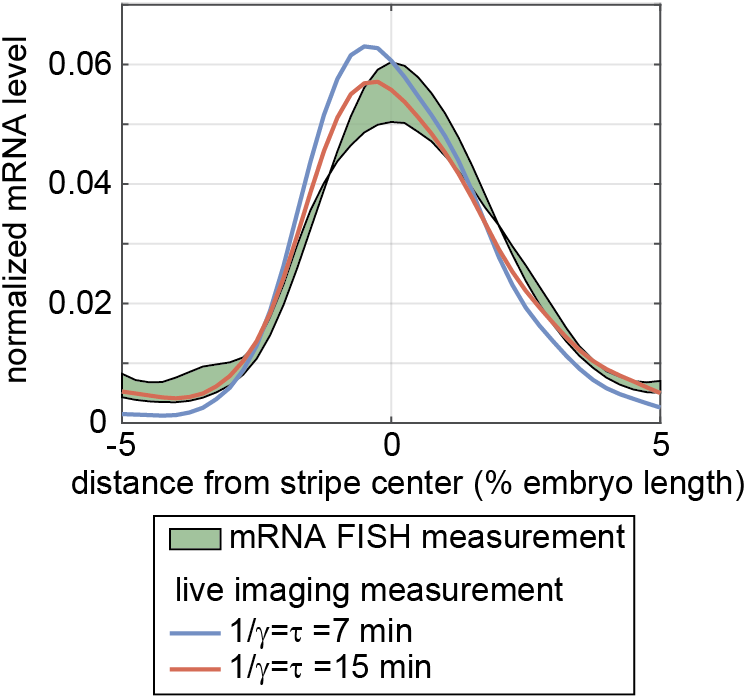
Comparison of predicted cytoplasmic mRNA by live-imaging measurements to direct measurements by FISH. In an effort to check the validity of our modelling assumptions, we compared the predictions of our mRNA model stemming from live-imaging measurements of stripe 2 of an *eve* reporter from a BAC containing the full *eve* locus to direct measurements of *eve* cytoplasmic mRNA levels using FISH (Luengo Hendriks et al., 2006). Here, the blue lines indicate our model’s predictions under two different assumptions for the rate of mRNA degradation, and the shaded green profile indicates the range of directly measured mRNA levels. Comparisons indicate a high degree of agreement between prediction and measurement, indicating that our modelling assumptions are justified.

#### Sensitivity of results to mRNA lifetime assumption

In the main text we assume a degradation rate for *eve* of 0.14 min^−1^ (corresponding to a lifetime of roughly *τ* = 7 min). Since, to our knowledge, the decay rate of *eve* mRNA has not been measured directly, we follow Bothma et al. (2014) and base this estimate off of measurements for another of the pair rule genes, *fushi tazu* (*ftz*, Edgar et al. (1987)). In this section, we examine the degree to which the apparent contributions of each regulatory strategy (Figure 1) change under different assumptions for *eve* mRNA lifetime. Rather than conducting an exhaustive survey, we instead focus primarily on two limiting cases: rapid mRNA decay (*τ* = 1 min) and *no* mRNA decay (*τ* = ∞).

Appendix 2–Figure 2 summarizes the results of our analysis. We find that, regardless of the assumed mRNA lifetime, our model predicts that *eve* stripe 2 is formed almost entirely via the interplay between the binary control of the transcriptional time window and the analog modulation of the mean rate of transcription (compare brown and red profiles in Appendix 2–Figure 2). However, we find that the relative importance of each factor depends, somewhat, on the assumed decay rate. In the case of rapid mRNA decay, as well as for the decay rate assumed in the main text, the time window (blue profile) is clearly the dominant factor in driving pattern formation (Appendix 2–Figure 2A and B). If we assume the true mRNA lifetime is 15 minutes, slightly more than double our best guess of 7 minutes, we find that the time window is still predicted to contribute slightly more to stripe formation, but that the two contributions are now of order with one another (Appendix 2–Figure 2C). Finally, in the limit where there is effectively no mRNA decay, the effects of the mean rate and time window are roughly equivalent (Appendix 2–Figure 2D). This result can be explained by the fact that the mean rate strategy is insensitive to the decay rate, whereas the effect of the time window is enhanced by the action of mRNA decay.

Thus, overall, we found that our model’s prediction that the control of the transcriptional time window plays a primary role in stripe formation holds for mRNA lifetimes less than or equal to 15 minutes, which is more than double the measured life time of *ftz* mRNA (Edgar et al., 1987). Perhaps more importantly, both factors are found to play a significant role, *irrespective* of mRNA decay rate, indicating that our central finding is robust to our assumption regarding mRNA decay dynamics.

**Appendix 2 Figure 2.**
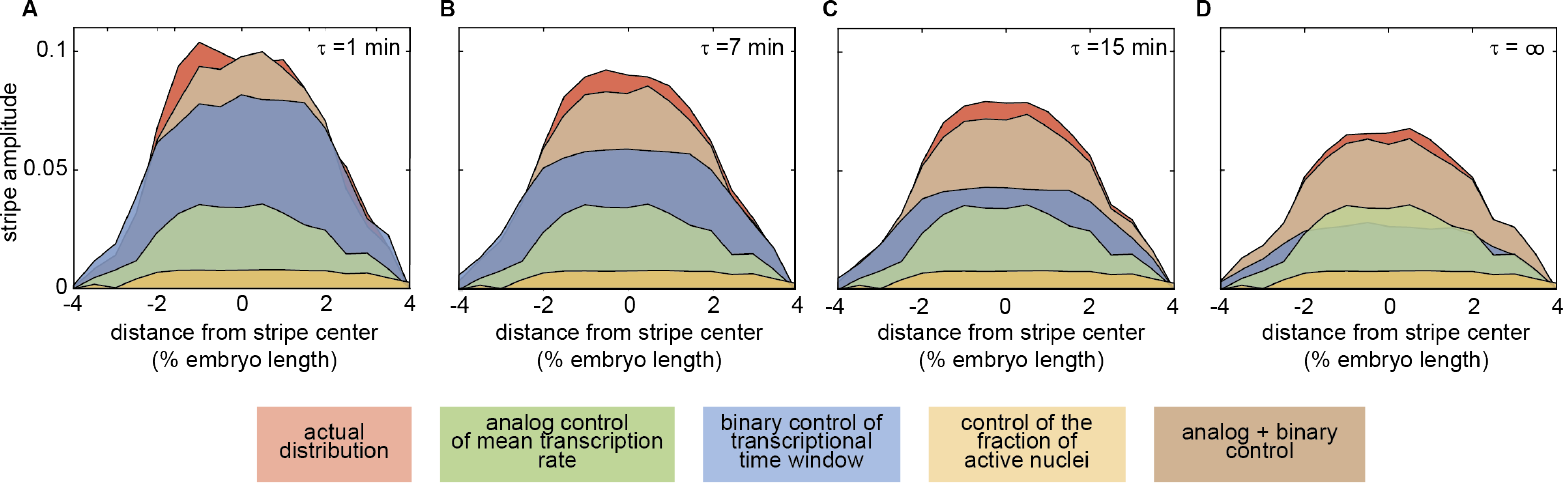
Sensitivity of regulatory strategy contribution to assumed mRNA lifetime. The average lifetime of *eve* mRNA is a significant assumed parameter in our model. This figure compares the predicted contributions of each regulatory strategy for the mRNA lifetime assumed in the main text (*τ* = 7 min) to limiting cases in which mRNA is assumed to decay almost instantaneously (*τ* = 1 min) on the one hand, and infinitely slowly on the other (*τ* = ∞). Even at these extremes, the central conclusion that the stripe is formed via the join action of mean rate modulation (green profile) and the time window (blue profile) remains intact. As expected, the relative contribution of the time window is sensitive to the assumed τ, yet even in the limit of no significant mRNA decay, its impact is still of order with the effect of mean rate modulation.

#### Control strategy contributions for *eve* BAC

A key question regarding the results in the main text is whether and to what degree the relative contributions of the regulatory control strategies we identified in Figure 1 and Figure 3 for the reporter containing only the *eve* stripe 2 enhancer hold true for the formation of the stripe in the endogenous context. While we cannot directly query activity at the endogenous *eve* locus, we were able to examine the dynamics of stripe formation for an *eve* BAC used in the companion paper to this manuscript (Berrocal et al., 2018). Since this BAC contains the full *eve* regulatory locus, it likely provides a better proxy for stripe formation in the endogenous context than the isolated *eve* 2 reporter. Appendix 2—Figure 3 shows the results of this analysis. As with the reporter construct used in the main text, we find that the stripe is formed primarily through the interplay between two regulatory strategies: the modulation of the average rate of production (green) and of the duration of transcriptional activity (blue). As with the reporter, the binary control of the transcriptional time window is the dominant driver of stripe formation (compare with Figure 1G). Interestingly, unlike the reporter construct, the full predicted profile (red profile) that accounts for the interplay between mRNA decay temporal fluctuations in the mean rate of mRNA production differs substantially from the simpler model (brown profile) that approximates mRNA production as constant over time. We speculate that this difference is attributable to the influence of the “late enhancer”—which is present in the *eve* BAC but *not* in the reporter—that takes over control of *eve* activity late in nc14. Further work will be necessary to fully elucidate the regulatory impact of this late element on the formation of the mature *eve* stripe pattern.

**Appendix 2 Figure 3.**
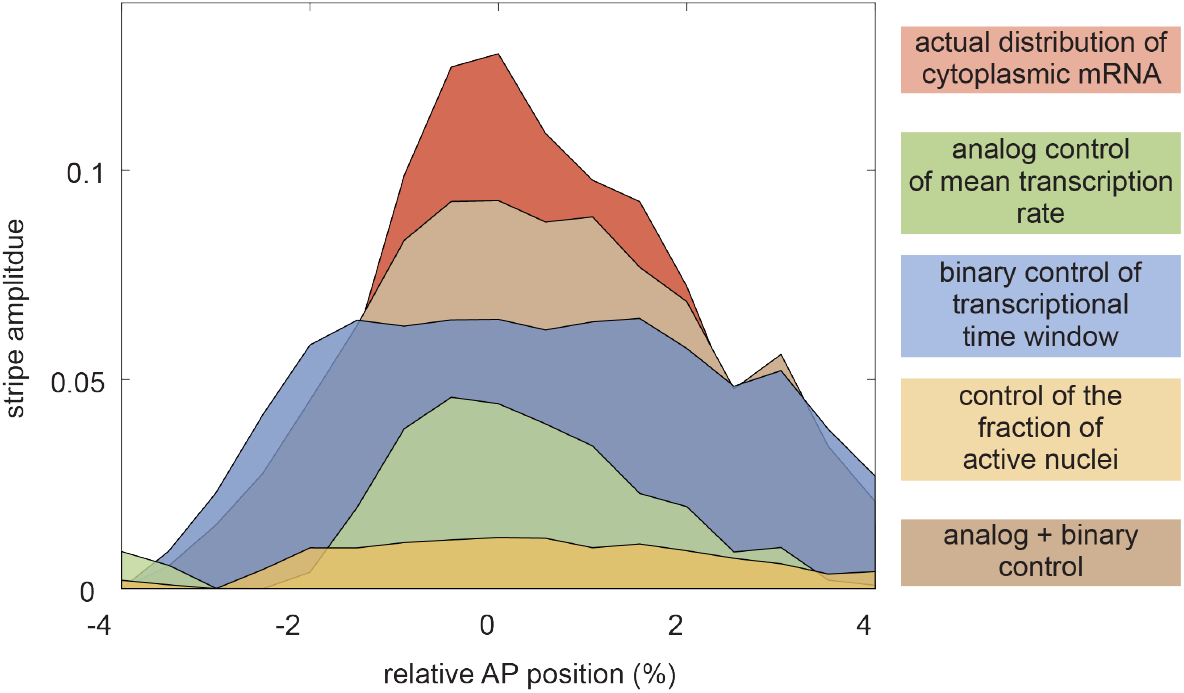
Regulatory strategy contributions to *eve* stripe 2 formation in endogenous context. As with the reporter construct, the formation of *eve* stripe 2 in the context of the full *eve* regulatory locus is dominated by the interplay between mean rate modulation (green) and control of the time window of transcriptional activity (blue).

## Appendix 3

### The memory-adjusted hidden Markov model Model introduction

To model the dynamics of an observed fluorescence series, ***y*** = {*y*_1_, *y*_2_, …, *y_T_*}, where *T* is the number of data points in a trace, we assume that, at each time step, the sister promoters can be in one of *K* effective states. In the analysis of *eve* stripe 2 data, we use a simple model with the number of effective states equal to three (*K* = 3). The method, however, allows for more complex transcription architectures with higher numbers of states. Transitions between the effective promoter states are assumed to be Markovian, meaning that the hidden promoter state *z_t_* at time step *t* is conditionally dependent only on the state in the previous time step. This dependency is modeled through a *K×K* transition probability matrix ***A*** = *p*(*z_t_*|*z*_*t*−1_), where *A_kl_* is the probability of transitioning from the *l*^th^ state into the *k*^th^ state in the time interval Δ*τ*, where Δ*τ* is the data sampling resolution. We assign a characteristic RNA polymerase initiation rate, *r*(*k*), with units of RNA polymerase per minute, to each effective promoter state, *z*(*k*), 1 ≤ *k* ≤ *K*. Thus, the number of polymerases initiated between time steps *t* − 1 and *t* will be *r*(*z_t_*)Δ*τ*, Because the fluorescence intensity contributed by each polymerase depends on the number of transcribed MS2 stem loops, the contribution will vary with the position of the polymerase on the gene. In our transcription model we assume that polymerase elongation takes place at a constant rate. Therefore, if *τ*_MS2_ is the time it takes to transcribe the MS2 loops, the fluorescence contribution of an RNA polymerase molecule will initially grow linearly (*τ* ≤ *τ*_MS2_) and will then stay constant for the remainder of transcription (*τ*_MS2_ ≤ *τ* ≤ *τ*_elong_). Given this time dependence, we define a maximum fluorescence emission per time step for each state as *υ*(*k*) = *F*_RNAP_*r*(*k*)Δ*τ*, 1 ≤ *k* ≤ *K*, where *F*_RNAP_ is the fluorescence calibration factor determined using smFISH experiments (see Materials and Methods).

**Appendix 3 Figure 1.**
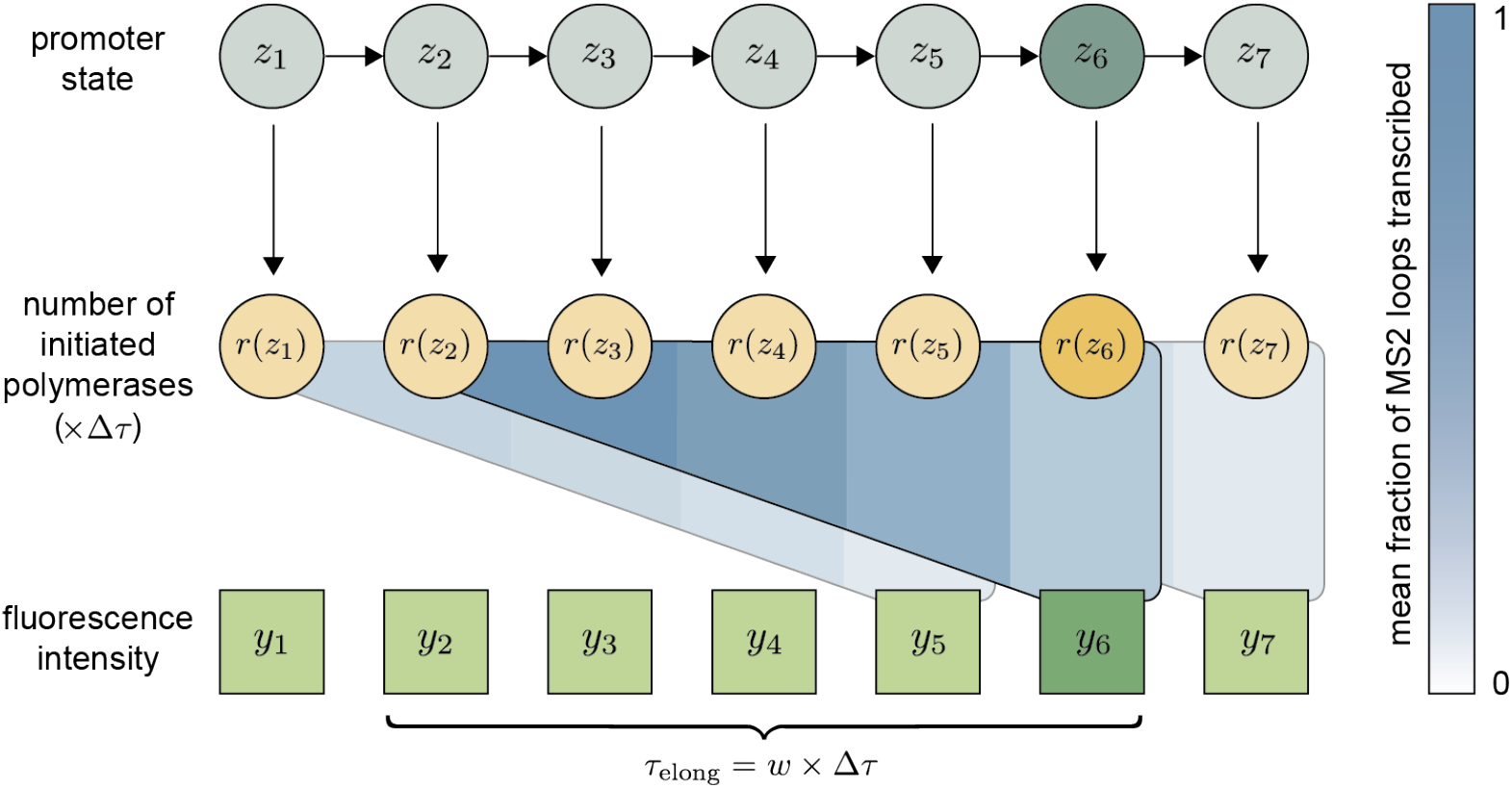
Schematic overview of the mHMM architecture. The sister promoters are modeled as undergoing a series of Markovian transitions between effective transcriptional states (*z*_t_). Each promoter state uniquely determines the number of polymerases initiated in a single time step (*r*(*z_t_*)Δ*τ*). Fluorescence emissions from polymerases initiated in the most recent *w* steps combine to produce the observed fluorescence intensity (*y_t_*). The color bar indicates the mean fraction of MS2 loops that have been transcribed and contribute fluorescence at the moment of observation. The color corresponding to the more recently initiated polymerases is therefore lighter (fewer loops transcribed) than that corresponding to polymerases initiated at earlier times(more loops transcribed).

The instantaneous fluorescence intensity is the cumulative contribution from polymerases initiated in the previous *w* time steps, where *w* = *τ*_elong_/Δ*τ* is the system-dependent integer memory. Here Δ*τ* indicates the observational time resolution, a quantity set by experimental conditions. The time required for an RNA polymerase molecule to transcribe our reporter gene (*τ*_elong_) is *a priori* unknown. We developed an autocorrelation-based method to estimate *τ*_elong_ directly from our experimental data (see Appendix 9 and Coulon and Larson (2016)). The observation *y_t_* at time step *t* conditionally depends not only on the hidden promoter state *z_t_*, but also on the hidden states in the previous *w* time steps, {*z_t_, z*_*t*−1_, …, *z*_*t*−*w*+1_}. To be able to describe the observed system dynamics through a hidden Markovmodel, the observation at time step *t* needs to be conditionally independent from the states at earlier time steps. We therefore introduce the concept of a compound state, *s_t_* = {*z_t_*, *z*_*t*−1_, …, *z*_*t*−*w*+1_}, which, together with the set of model parameters, ***θ***, is sufficient to define the probability distribution of the observation *y_t_*, thereby satisfying the Markov condition. Since *z_t_* ∈ {1, …, *K*}, each compound state can take one of *K^w^* different values, *s_t_* ∈ {1, …, *K^w^*}. While the number of possible compound states is *K^w^*, only *K* different transitions are allowed between them, since the most recent *w* − 1 promoter states are deterministically passed from one compound state to the next, i.e. the last *w* − 1 elements in *s_t_*_+1_ = {*z*_*t*+1_, *z_t_*, …, *z*_*t*−*w*+2_} are present in *s_t_* as well. The schematic overview of the mHMM architecture is shown in Appendix 3–Figure 1.

We model the fluorescence emission probabilities corresponding to each hidden compound state as Gaussian distributions with a standard deviation *σ*, which we learn during inference. The joint probability distribution *p*(***y, s|θ***) of the series of hidden compound states, ***s*** = {*s*_1_ *s*_2_, …, *s_T_*}, and fluorescence values, ***y*** = {*y*_1_, *y*_2_, …, *y_T_*}, is given by

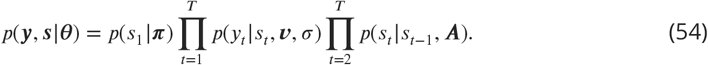

Here ***π*** is a *K*-element vector, with *π_k_* being the probability that the trace starts at the *k*^th^ effective promoter state, and ***ν*** is a *K*-element vector of fluorescence emission values per time step.

Our goal is to find an estimate of the model parameters, 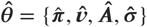, which maximizes the likelihood *p*(***y|θ***) of observing the fluorescence data, namely,

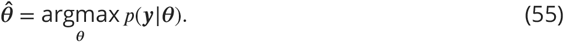

The likelihood can be obtained by marginalizing the joint probability distribution, *p*(***y, s|θ***), over the hidden compound states, that is,

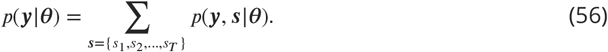

Note that the summation is performed over all possible choices of ***s*** — a vector of *T* elements, each of which can take *K^w^* possible values. The total number of terms in the sum is thus equal to *K^wT^*, which grows exponentially with the number of time points. To make the estimation of the model parameters tractable, we use an approximate inference method, the expectation-maximization (EM) algorithm.

We note that the notion of a compound state was also introduced in an earlier work (Corrigan et al., 2016)to account for the memory effect in hidden Markov modeling of *actin* transcription and then an EM methodology was applied to learn the kinetic parameters from MS2-based transcription data. Unlike their approach, however, we do not explicitly model the recruitment of individual RNA polymerase molecules, but instead, assign a continuous RNA polymerase initiation rate to each promoter state. Additionally, our model estimates the magnitude of the background noise present in the experimentally measured fluorescence signal, whereas the model presented in (Corrigan et al., 2016) takes this quantity as an input, requiring that it be estimated separately. We believe that these differences serve to make our model more flexible. Moreover, by eliminating the need for absolute calibration and noise estimation, we hoped to facilitate the use of our model in a wide variety of experimental contexts, for which one or the other quantity may not be readily obtainable. In the “Continuous vs. Poisson promoter loading” section of Appendix 4 we demonstrate that relaxing the continuous RNA polymerase loading assumption when generating synthetic data does not significantly affect the accuracy of the mHMM inference.

#### Expectation-maximization (EM) algorithm

Consistent with standard EM approaches (Bishop (2006), Chapter 13), at each iteration we maximize the lower bound of the logarithm of the likelihood using the current estimate of the model parameters, namely,

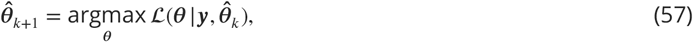

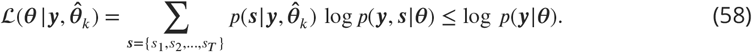

Here 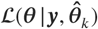 is the objective function, 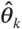 is the estimate of the model parameters in the *k*^th^ expectation step of the EM algorithm. Since we model the transitions between the effective sister promoter states as a Markov process, the logarithm of the joint probability distribution, log*p*(***y, s|θ***), can be written as

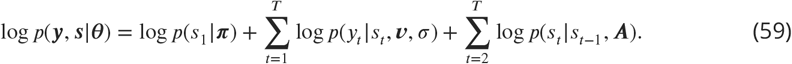

Now, we introduce several notations: 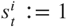 if and only if *s_t_ = i*; Δ(*s_t_, d*): = the *d*^th^ digit of the promoter state sequence *s_t_* = {*z_t_, z*_*t*−1_, …, *z_t_*−(*w*−1)}, starting from the left end; *C_zs_* = 1 if and only if Δ(*s*, 1) = *z*; *B_s′, s_* = 1 if and only if the transition *s → s′* between the compound states *s* and *s′* is allowed, which happens when the latest (*w* − 1) promoter states in the compound state *s* match the earliest (*w* − 1) promoter states of the compound state *s*′. With these notations in hand, the terms in Equation 59 can be rewritten as

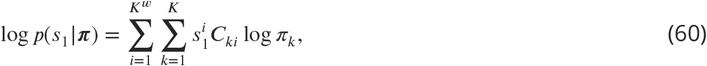

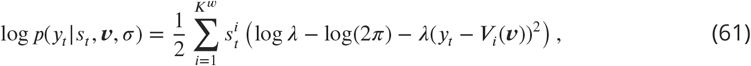

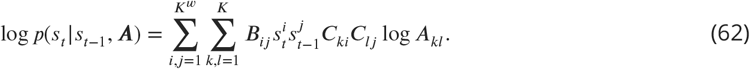

Here *λ* = 1/*σ*^2^ is the Gaussian precision parameter, and *V_i_*(***ν***) is the aggregate fluorescence produced in the *w* consecutive promoter states of the *i*^th^ compound state.

Because of the finite time *τ*_MS2_ it takes a single polymerase to transcribe the MS2 sequence, the fluorescence contribution of polymerases is weighted at different positions in the window of *w* time steps. If we define *n*_MS2_ = *τ*_MS2_/Δ*τ* as the number of time steps (not necessarily an integer) necessary for transcribing the MS2 sequence, the mean fraction of the full MS2 sequence transcribed by a polymerase at the *d*^th^ time step of the elongation window will be given by

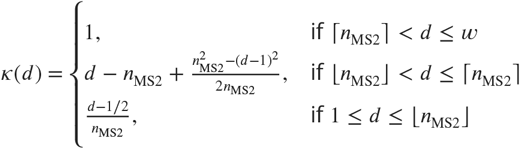

where ⌈*n*_MS2_⌉ and ⌊*n*_MS2_⌋ are the ceiling and the floor of *n*_MS2_, respectively. The dependence of the weighting function *κ*(*d*) on the position for a specific choice of parameters is illustrated in Appendix 3–Figure 2.

**Appendix 3 Figure 2.**
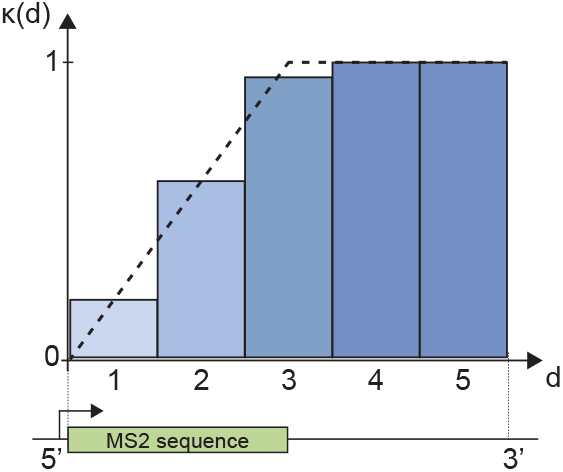
The weighting function *κ*(*d*) evaluated at different positions along the genome. The dashed line represents the fraction of the MS2 loops transcribed at a given position. Parameters used for plotting: *τ*_elong_ = 100 sec, *τ*_MS2_ = 50 sec, Δ*τ* = 20 sec, *w = τ*_elong_/Δ*τ* = 5, *n*_MS2_ = *τ*_MS2_/Δ*τ* = 2.5.

Accounting for the weighted fluorescence contribution of polymerases, the aggregate fluorescence *V_i_*(***ν***) becomes

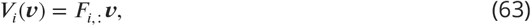

where the *i*^th^ row of the *K^w^ × K* matrix ***F*** is the number of times each promoter state is present in the *i*^th^ compound state, weighted by the position-dependent function *κ*(*d*). For example, if we consider a promoter with *K* = 3 states and memory *w* = 5, then the row of ***F*** corresponding to the compound state *s* = {1, 1, 3, 2, 3} will be [*κ*(1) + *κ*(2), *κ*(4), *κ*(3) + *κ*(5)].

Having all the pieces of the logarithm of the joint probability distribution, log *p*(***y, s|θ***), we obtain a final expression for the objective function, namely,

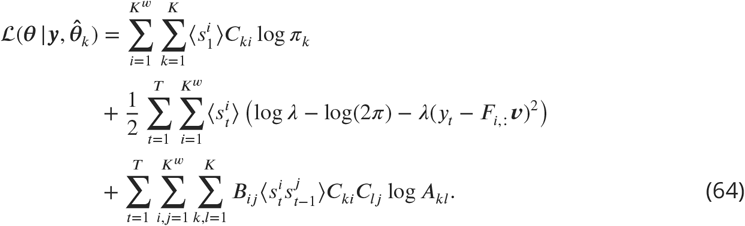

Here 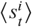 and 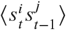 are the expectation coefficients at the *k*^th^ step of the EM algorithm defined as

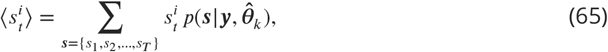

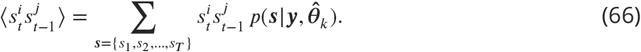

Using the current estimate of the model parameters, 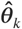, the expectation coefficients 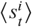 and 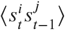 are calculated using the forward-backward algorithm. From the definitions in Equation 65 and Equation 66, we obtain

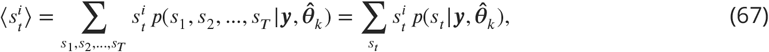

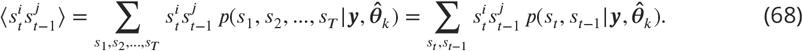

Following the conventional implementation of the forward-backward algorithm (cf. Bishop (2006), Chapter 13), we use the Markov property of the promoter state dynamics, together with the sum and products rules of probability, to write

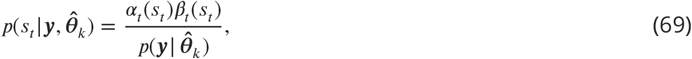

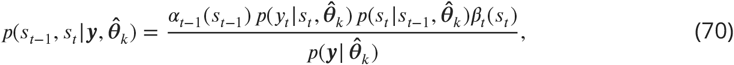

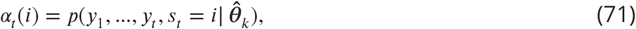

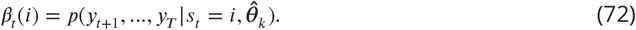

Here *α_t_*(*i*) is the joint probability of observing the fluorescence emission values in the first *t* steps and being at the *i*^th^ compound state at step *t;* while *β_t_*(*i*) is the conditional probability of observing fluorescence values from the time point (*t* + 1) till the end of the series, given that the compound state at time *t* is *i*. Note that *α* and *β* can be treated as *K^w^* × *T* matrices, where each column is a vector of length *K^w^*, accounting for the *K^w^* possible values of *i* in Equation 71 and Equation 72. We evaluate the elements of *α* and *β* matrices recursively as

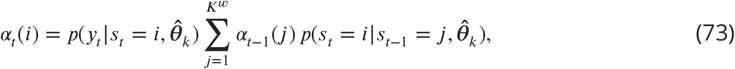

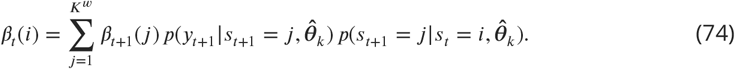

The boundary values for *α*_1_(*i*) and *β_T_*(*i*) at the first and last columns of *α* and *β* matrices, respectively, are given by

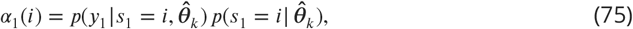

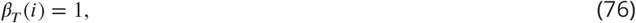

where the first follows from the definition of *α_t_*(*i*), and the second is obtained from Equation 69 by setting *t = T*. Having evaluated the *α* and *β* matrices, the likelihood 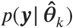 that appears in the denominator of Equation 69 and Equation 70 can be found by setting *t = T* in Equation 69 and summing over *s_T_*, namely,

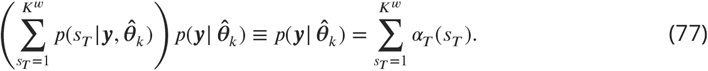

With the probabilities 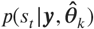 and 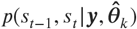 known, the expectation coefficients follow directly from Equation 67 and Equation 68.

The optimal model parameters in the (*k* + 1)^th^ step of the EM algorithm are obtained by maximizing the objective function 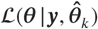 in Equation 64 with respect to {***π, ν**, λ, **A***}, subject to the probability constraints 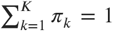 and 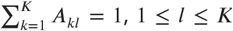. The update equations for the model parameters are found as

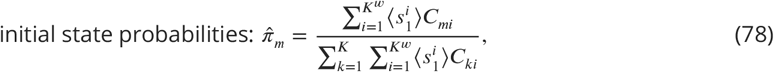

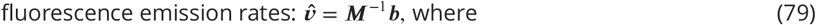

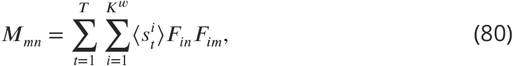

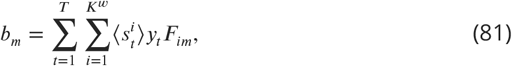

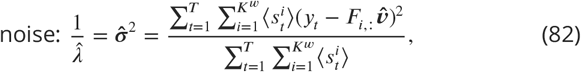

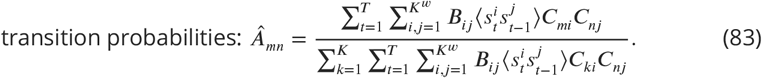

#### Pooled inference on multiple traces

Since the information available in a single MS2 fluorescence trace is not sufficient for the accurate inference of underlying model parameters, we perform a pooled EM inference assuming that the traces are statistically independent and governed by the same parameters. If ***y***_1:*N*_ are *N* different fluorescence traces with corresponding trace lengths *T*_1:*N*_, and ***s***_1:*N*_ are the hidden compound state sequences corresponding to each trace, we obtain

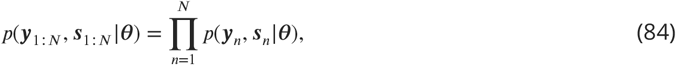

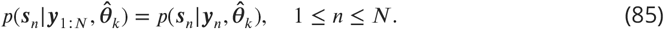

Therefore, the objective function 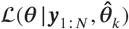 maximized at each EM iterations takes the form

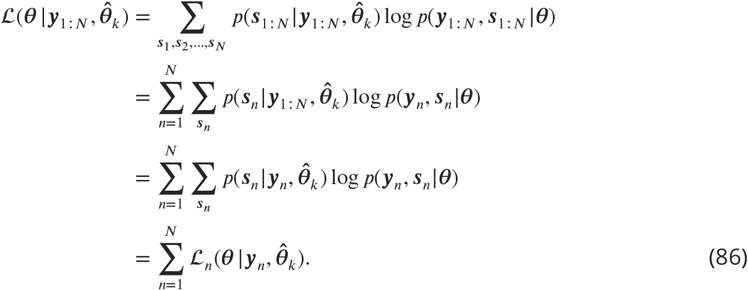

From the above equation, we recognize that the objective function for the pooled inference is the sum of objective functions written for each individual trace. Using the expression for the single-trace objective function obtained earlier (Equation 64), we find

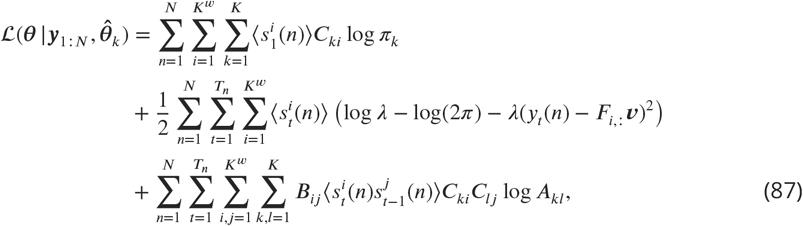

where 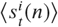 and 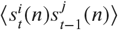 are now the expectation coefficients obtained for the *n*^th^ fluorescence trace via the forward-backward algorithm, and *y_t_*(*n*) is the fluorescence at time step *t* in the *n*^th^ trace. The update equations are then derived analogous to the single-trace case, with an additional summation performed over all traces, namely,

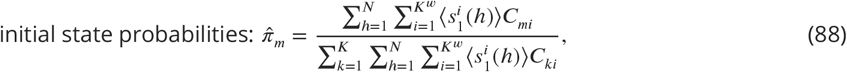

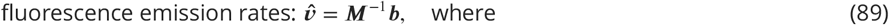

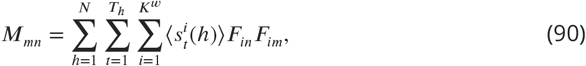

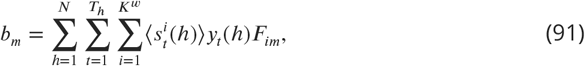

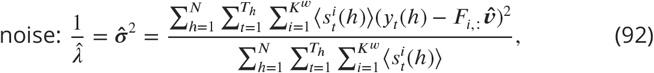

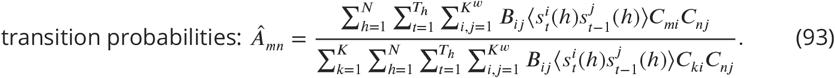

#### Execution of the mHMM method

Execution of the mHMM method starts by initializing the model parameters. ***π*** and each column of ***A***, both of which are vectors of size *K*, are initialized by randomly sampling from a Dirichlet distribution given by

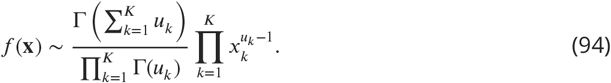

The Dirichlet distribution parameters *u_k_* are all set equal to one, which makes each initial promoter state equally likely to be occupied, and equally likely to be transitioned into.

To initialize the fluorescence emission rates, ***r***, and the Gaussian precision parameter, *λ* = 1/*σ*^2^, we first treat the fluorescence data ***y***_1:*N*_ as identical and independently distributed (i.i.d.) and use a simplified time-independent EM algorithm to find their optimal values (cf. Bishop (2006), Chapter 13). We initialize the highest emission rate by randomly choosing a value between 70%and 130% of the highest emission rate inferred by the i.i.d. approach. The lowest emission rate is initialized to 0 because of the apparent silent periods in the activity traces. The remaining (*K* − 2) emission rates are initialized by choosing random values between 0 and the highest emission rate. Finally, we initialize the Gaussian noise *σ* by randomly choosing a value between 50% and 200% of the noise inferred by the i.i.d. approach.

After initializing the model parameters, we iterate between the expectation and maximization steps of the EM algorithm until the relative changes in the Euclidean norms of the model parameters after consecutive iterations become smaller than *ε* = 10^−4^ or the number of iterations exceeds 500. Because EM approaches typically infer locally optimal parameter values, the algorithm is run on the same dataset using multiple randomly chosen initial parameters (10-20 in our implementations), and the globally optimal set of values is chosen in the end. In the Matlab implementation of the EM algorithm, the variables are all stored in logarithmic forms to avoid overflow and underflow issues, which could occur when recursively evaluating the elements of the *α* and *β* matrices. Also, special care is taken when accounting for time points less than the elongation time, i.e. *t < w*, in which case the compound state is a collection of not *w*, but *t* promoter states, i.e. *s_t_* = {*z_t_, z*_*t*−1_, …, *z*_1_}.

Because of the exponential scaling of the model complexity with the integer memory window (*w* = 7 for the *eve* construct with Δ*τ* = 20 sec data sampling resolution), significant computational resources were used when conducting inference on simulated and experimental data. It took approximately 2 hours to conduct 25 mHMM inferences with different initialization conditions on a machine with 24 CPU cores. Users of the mHMM method are advised to have this metric as a reference when estimating the computational cost of their inference.

### Windowed mHMM

To investigate temporal trends in bursting parameters, we extended the mHMM method to allow for a sliding window inference approach. From a technical perspective, this required a revision of the inference formalism to be compatible with fragments of fluorescent traces in which the beginning of the trace (initial rise in *y_t_* from *t* = 1) was not included.

To that end, we modified the first term in Equation 59 to allow for all possible promoter state sequences that could lead to the observation of the first fluorescence measurement in the chosen time window ([*T*_1_, *T*_2_]), namely,

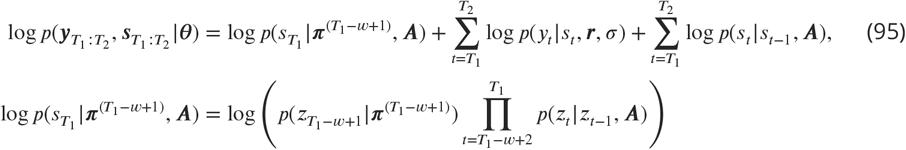

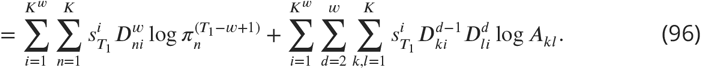

Here ***π***^(*T*_1_−*w*+1)^ is the probability distribution of the earliest promoter state that still has an impact on the observation of the first measurement in the sliding window, and 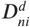 is an indicator variable which takes the value 1 only if the promoter state in the *d*^th^ position of the *i*^th^ compound state is *n*.

The modified expression for the joint probability distribution does not change the functional form of the equations used for calculating the expectation coefficients. Maximization equations for the emission rates and the noise also remain intact. Only the maximization equation for the transition probabilities is revised from Equation 83 into

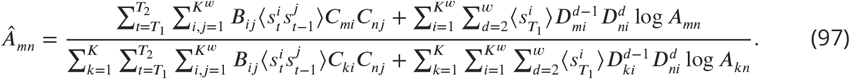

We make a steady-state assumption within the sliding window and choose ***π***^(*T*_1_−*w*+1)^ to be the stationary distribution of the current transition probability matrix, i.e. ***Aπ***^(*T*_1_−*w*+1)^ = ***π***^(*T*_1_−*w*+1)^. We therefore use the current estimate of ***A*** to evaluate ***π***^(*T*_1_−*w*+1)^ at each EM iteration, instead of performing a maximization step.

## Appendix 4

### Statistical validation of mHMM

We validated mHMM for the three-state (*K* = 3) architecture schematically illustrated in Appendix 4–Figure 1A by generating synthetic trajectories of effective promoter states using the Gillespie algorithm (Gillespie, 1976) and adding Gaussian noise to the resulting activity traces. Parameters in Appendix 4–Table 1 were used for data generation. Pooled inferences were conducted on 20 independent datasets, each containing 9,000 data points, representative of the number of experimental data points in a central stripe region. The top panel of Appendix 4–Figure 1B shows the kinetic architecture used to simulate the promoter trajectory in Appendix 4–Figure 1C (yellow) as it switches through the multiple possible states. This promoter trajectory leads to the simulated trace of the number of RNA polymerase molecules actively transcribing the gene in Appendix 4–Figure 1D (red). Using mHMM, we found the best fitted path for our observable (Appendix4–Figure 1D, black) and the corresponding most likely promoter state trajectory (Appendix 4–Figure 1C, blue).

**Appendix 4 Figure 1.**
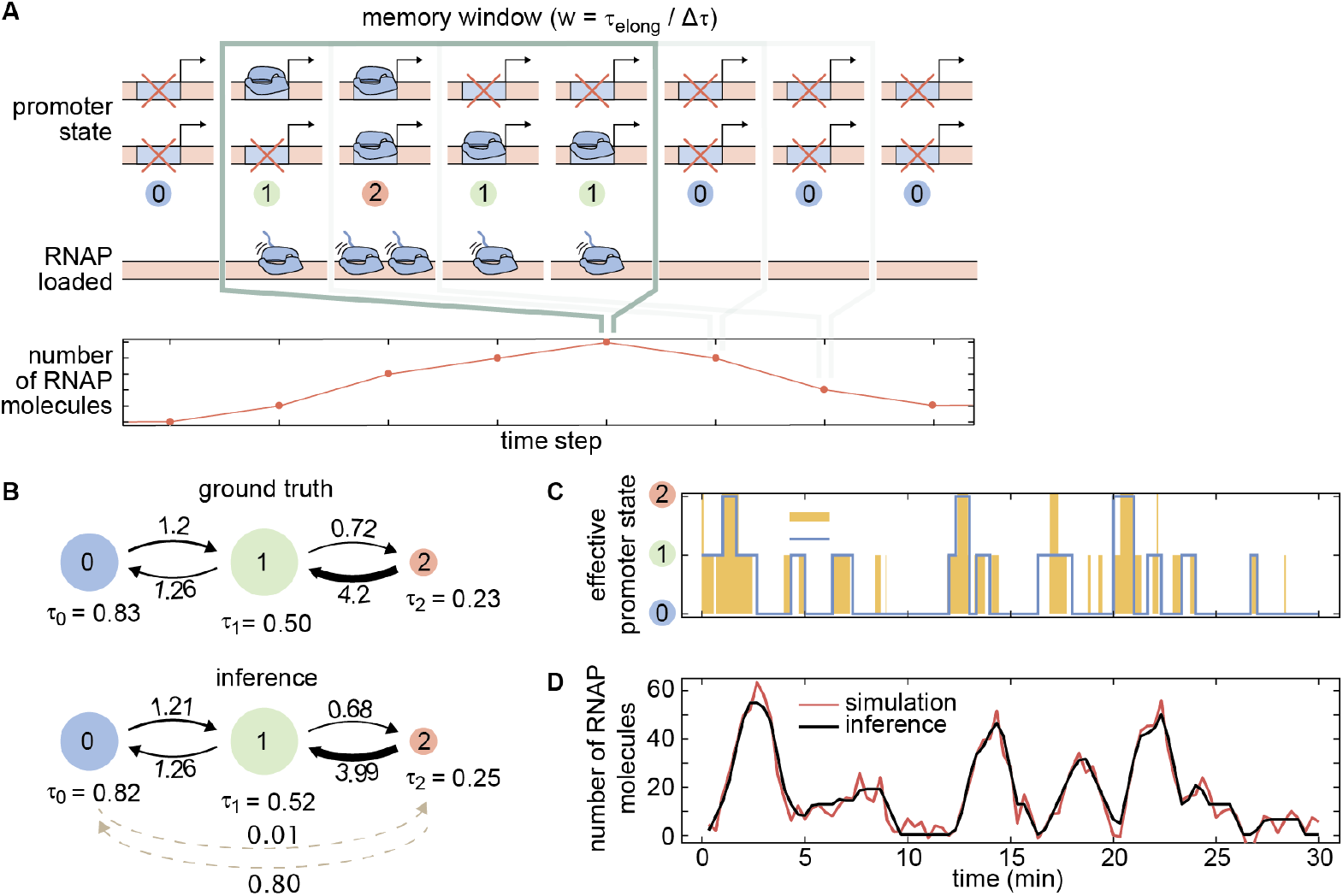
Statistical validation of mHMM. **(A)** Three-state mHMM architecture where ON and OFF promoter states on each sister chromatid result in an effective three-state model. The trajectory of effective promoter states over the memory time window given by the elongation time dictates the number of RNA polymerase molecules loaded onto the gene. **(B)** Flow diagrams of promoter states and transition rates for the true parameters used to simulate trajectories (top) and corresponding average inference results obtained from 20 independent datasets (bottom). The area of each state circle is proportional to the relative state occupancy, and the thickness of the arrows is proportional to the transition rates. Dashed lines correspond to inferred transitions with very slow rates that were absent in the simulation. Rates are in min^−1^ and dwell times are in min. Error bars for the mean inferred parameters are shown in Appendix 4–Figure 2. **(C)** Sample simulated promoter activity trace (yellow) generated using the parameters in (B), overlaid with the best fitted trace (blue) obtained using the Viterbi algorithm (Viterbi, 1967). **(D)** Simulated and best fitted observable number of RNA polymerase molecules corresponding to the promoter trajectory shown in (C).

**Appendix 4 Table 1.** Parameter values used for generating synthetic datasets in the statistical validation of the model. In order to perform this validation, we chose parameters that approximated those obtained through the mHMM inference on experimental data shown in Figure 5.

| Parameter | Value |
| --- | --- |
| Promoter switching rates ($k_{01}, k_{10}, k_{12}, k_{21}$) | (1.2, 1.26, 0.72, 4.2) $\text{min}^{-1}$ |
| RNAP initiation rates ($r_0, r_1, r_2$) | (0, 18.5, 46) RNAP/min |
| Measurement noise ($\sigma$) | 4.5 RNAP |
| RNAP elongation time ($\tau_{\text{elong}}$) | 140 sec |
| Data sampling resolution ($\Delta\tau$) | 20 sec |
| Memory window ($w = \tau_{\text{elong}}/\Delta\tau$) | 7 |
| MS2 loop transcription time ($\tau_{\text{MS2}}$) | 30 sec |
| Duration of each trace | 30 min |
| Number of time points per dataset | 9,000 |
| Number of traces per dataset | 100 |
| Number of independent datasets | 20 |

As shown in Appendix 4–Figure 1B and Appendix 4–Figure 2, comparison of the simulated and inferred parameters indicates that we reliably recovered the parameters used to generate our simulated data with high precision. We accurately inferred transition rates, dwell times, fraction of time spent in each state, and the rates of RNA polymerase loading over 20 independent datasets of simulated traces.

**Appendix 4 Figure 2.**
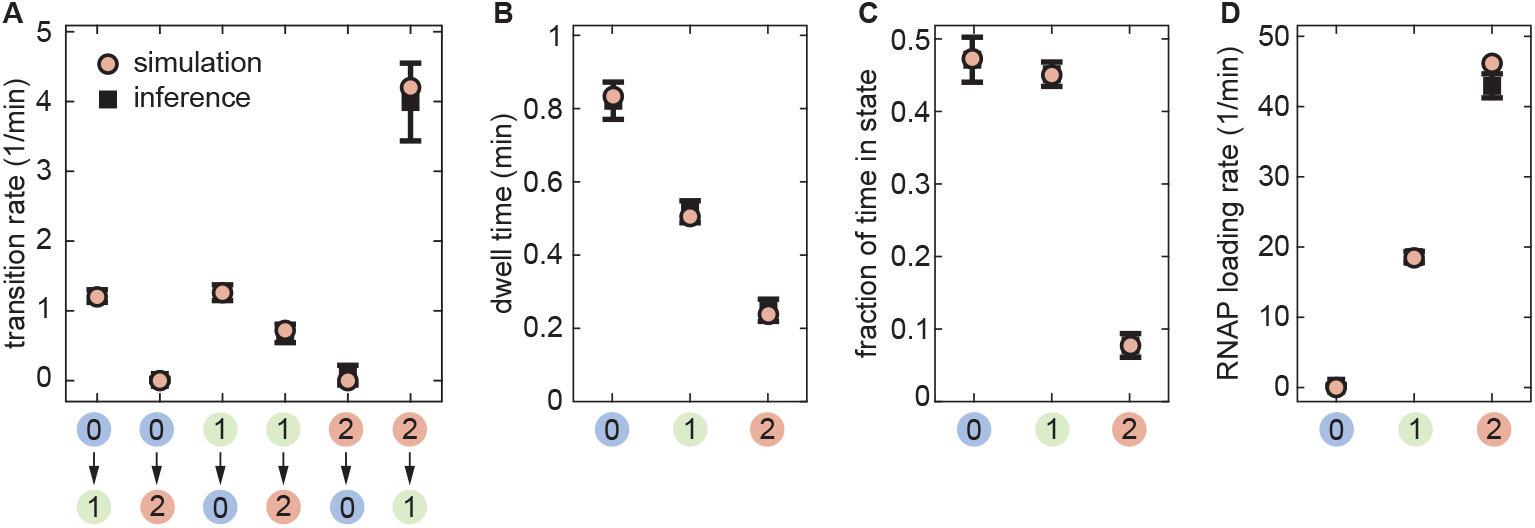
Inference statistics for the mHMM validation. The true and inferred values of **(A)** transition rates, **(B)** dwell times in states, **(C)** state occupancies, and **(D)** RNA polymerase loading rates are compared. Statistics on the inferred values are obtained from 20 independently generated datasets. (Error bars indicate one standard deviation calculated across these 20 independent replicates).

#### Validation details

We used the relation between the transition rate matrix, **R**, and the inferred transition probability matrix, **A**, defined in Appendix 3 to obtain estimates of the transition rates, namely,

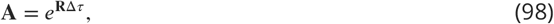

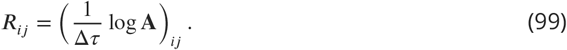

Here, the exponential and logarithm operations act on matrices **R**Δ*τ* and **A**, respectively. Occasionally, taking the matrix logarithm of the transition probability matrix **A** yielded small negative values for transition rates between states (0)and (2), which were originallyzero duringdata generation. In those cases, we assigned them a 0 value to keep them physically admissible.

#### Continuous vs. Poisson promoter loading

To demonstrate the validity of our choice to use continuous RNA polymerase initiation rates in the transcription model (Appendix 3), we repurposed our simulation to, instead of considering a constant rate of RNA polymerase loading, explicitly account for individual RNA polymerase loading events when generating the traces. We assumed that individual polymerase molecules traverse at a constant elongation rate (*υ*_elong_ = 46 bp/sec, Appendix 9) and that their arrival to the promoter region has a Poisson waiting time distribution, provided that the promoter is cleared from the previous polymerase molecule which has a finite footprint size of *l*_RNAP_ = 50 bp (Rice et al., 1993). This led to a two-step model for the process of RNA polymerase initiation, with Poisson-distributed wait times for the recruitment of RNA polymerase to the promoter followed by a finite wait period as the RNA polymerase cleared the promoter—a process taken to be approximately deterministic. With this information in hand, we expressed the mean loading time of RNA polymerase at a single promoter 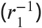 as the sum of the mean time of polymerase arrival at an empty promoter, 〈*τ*_arrival_〉, and the time required to clear it after arrival, 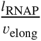, that is,

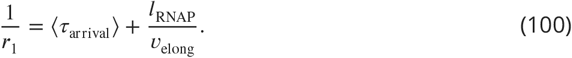

Having the values of *r*_1_, *l*_RNAP_, and *υ*_elong_, we found 〈*τ*_arrival_〉 and used it in simulating the arrival events of individual polymerases.

We performed inference on these simulated traces using mHMM with the objective of determining whether a Poisson loading rate had an effect on the obtained parameters. As shown in Appendix4–Figure 3, when the data is generated using Poisson RNA polymerase loading, mHMM slightly overestimates the high transition rate, but otherwise manages to accurately recover the model parameters. This therefore justifies our modeling approach of assigning continuous RNA polymerase initiation rates to each promoter state, instead of explicitly modeling the recruitment of individual polymerases.

**Appendix 4 Figure 3.**
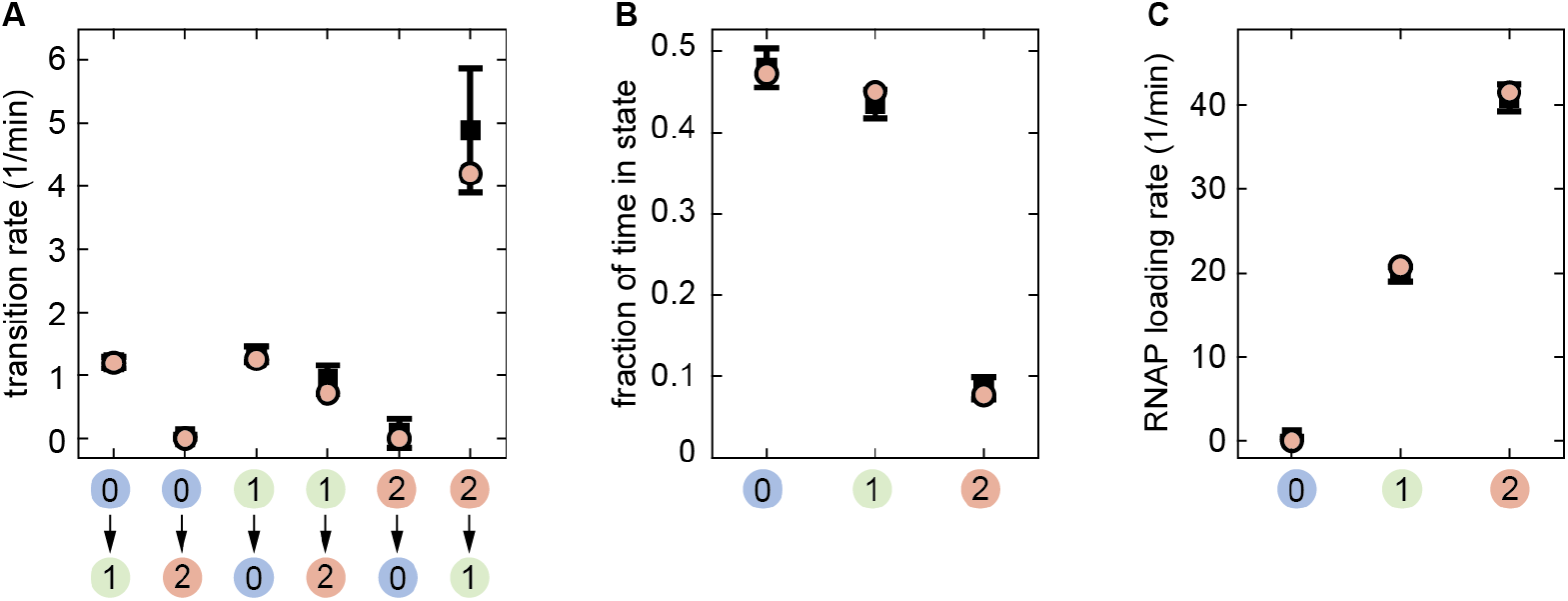
Validation of mHMM on Poisson RNA polymerase loading data. **(A)** Transition rates, **(B)** state occupancies and **(C)** RNA polymerase loading rates inferred from 15 independently generated datasets assuming Poisson loading of RNA polymerase. (Error bars represent one standard deviation calculated across these 15 independent replicates.)

#### Sensitivity of mHMM to data sampling resolution

In our mHMM framework, we modeled the stochastic transitions between effective promoter states using a discrete time Markov chain model which assumes that the state of the promoter remains constant during the experimental time step (Δ*τ*), and that transitions to the next promoter state can occur only at the end of each step. This means that, if the fastest promoter switching rate is greater than the data sampling rate (1/Δ*τ*), ourmodel might be unable to capture all those transitions. To study this possible limitation of mHMM, we conducted inference on synthetic activity traces generated with varying sampling rates. Since the system memory (*w* = *τ*_elong_/Δ*τ*) needs to be an integer, we varied *w* in the [3, 7] range, correspondingly changing the sampling resolution from low (*τ*_elong_/3 ≈ 46s) to high (*τ*_elong_/7 = 20s). We used the values in Appendix 4–Table 1 for the remaining model parameters.

Appendix 4–Figure 4 summarizes the findings of this study. As expected, the accuracy of inference improves with increasing data sampling rate, and inference results get very close to the ground truth values when the highest sampling rate (1/20 sec = 0.05s^−1^) becomes comparable to the fastest transition rate (0.07s^−1^). Except for the fastest transition rate, all other rates are inferred accurately for the whole spectrum of sampling resolutions (Appendix 4–Figure 4A). The accuracy of inferred state occupancies is also remarkably high, making it robust to variations in the data sampling rate (Appendix 4–Figure 4B). The high RNApolymerase loading rate tends to be underestimated for slower sampling resolutions, which is reasonable since the chances of promoter leaving state (2) during a single time step become greater, effectively reducing the net rate of loaded RNAP molecules per time step (Appendix4–Figure 4C). Generally, we find the inference of model parameters to be reasonably accurate for the entire spectrum of experimentally realizable data sampling rates, and highly accurate when the timescale of the fastest transition and data sampling are comparable.

**Appendix 4 Figure 4.**
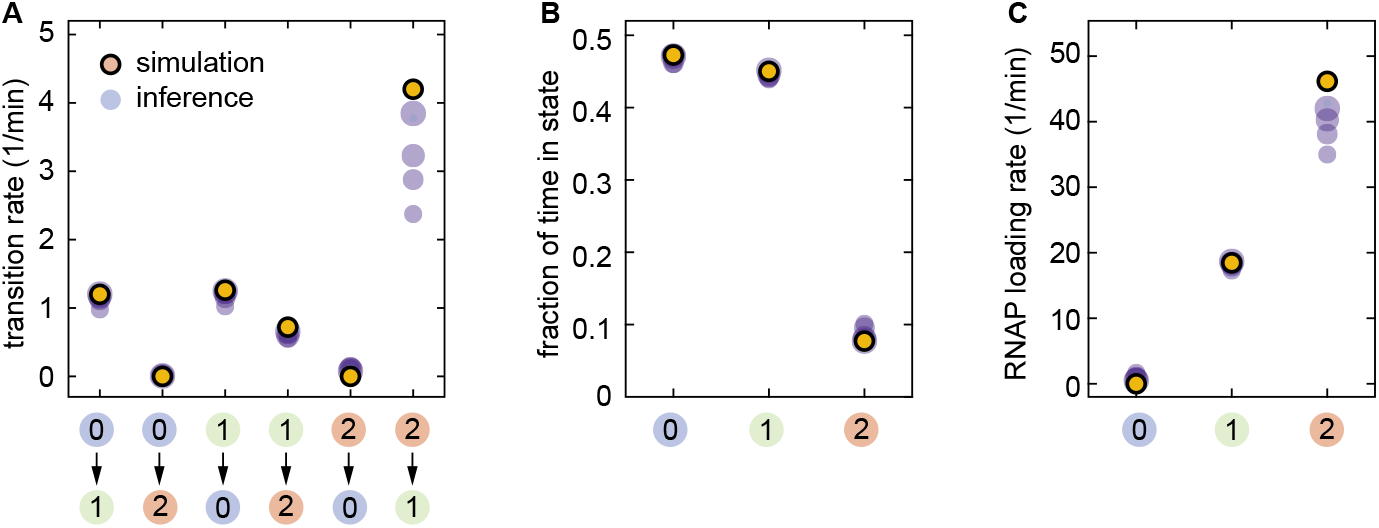
Sensitivity of mHMM to data sampling resolution. **(A)**Transition rates, **(B)** state occupancies and **(C)** RNA polymerase loading rates inferred from datasets generated with varying time resolutions. Transparent circles represent averages over 20 independently generated samples. The increasing size of the blue circles corresponds to higher data sampling resolutions (largest: 20s, smallest: 46s).

#### Performance of mHMM in different kinetic regimes

Thus far, the validation of mHMM was performed on datasets that were generated using parameters similar to those inferred for the *eve* promoter. These parameters have characteristic low ON rates (*k*_01_, *k*_12_) and a high OFF rate (*k*_21_), where “low” and “high” are relative to the data sampling frequency, which for our experimental setup is 3/min. To assess the utility of our inference method for a generic choice of model parameters, we performed additional inference studies in three different parameter regimes: low ON rates and low OFF rates (Appendix4–Figure 5A-C), high ON rates and low OFF rates (Appendix4–Figure 5D-F), and high ON rates and high OFF rates (Appendix4–Figure 5G-I).

As expected, the inference is the most accurate when the data sampling frequency is greater than the transition rates (Appendix 4–Figure 5A-C), in which case multiple transitions within a single time frame occur only rarely, making our discrete Markovian representation of the state dynamics a valid approximation. The largest deviations of the inferred model parameters from their ground truth values occur when the ON rates are high and the OFF rates are low (Appendix 4–Figure 5D-F). Since the promoter rarely remains in the lower initiation states (0 or 1) for the entire duration of a frame and tends to rapidly transfer into a higher initiation state (1 or 2, respectively), the rates of RNA polymerase loading for states 0 and 1 are significantly overestimated Appendix 4–Figure 5F). Despite the inaccuracies in estimating the RNA polymerase loading rates, all transition rates, with the exception of *k*_10_, are inferred with a high accuracy (Appendix 4–Figure 5D). Remarkably, the deviations caused by the high ON rates get substantially suppressed when the OFF rates are also made comparably high (Appendix 3–Figure 5G-I). This can be thought ofasa consequence of an effective counterbalancing between unwanted ON and OFF transitions within a single time frame.

Overall, these additional studies, together with the statistical validation studies discussed earlier (Appendix 4–Figure 2), elucidate the domain of applicability of mHMM: the method performs accurate inference when the ON/OFF transition rates are respectively slow/slow, slow/high, or high/high; and is not successful in accurately inferring some of the model parameters when the ON rates are high, but the OFF rates are low. We hope that these characteristics of the method will be useful in informing the design of promoter architectures and new experiments.

**Appendix 4 Figure 5.**
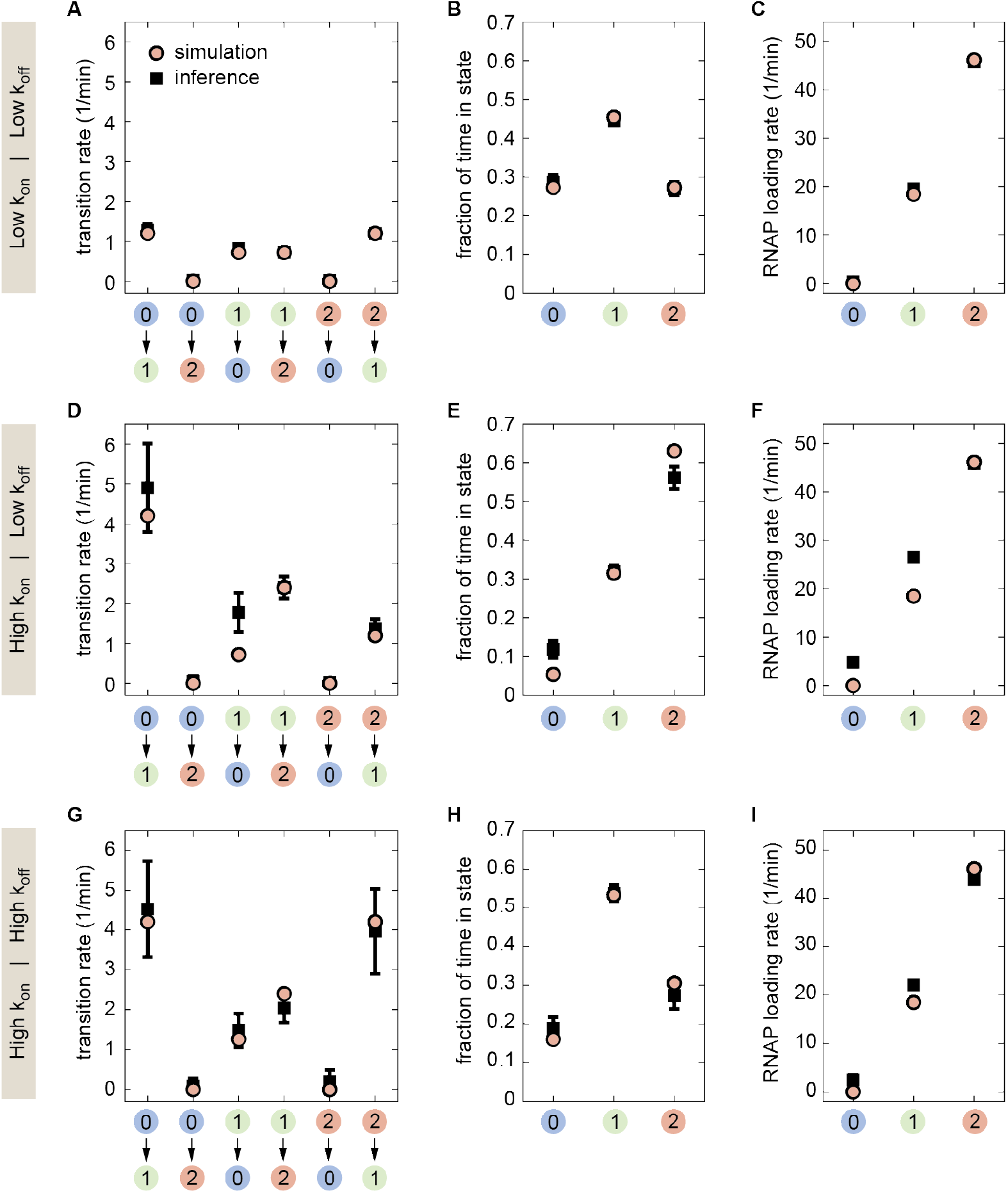
Study of mHMM performance for different choices of the ON/OFF transition rates. Comparison of inference performance for different ON/OFF rates using a data sampling frequency of 3/min. **(A-C)** low/low, (**D-F**) high/low, (**G-I**) high/high. The statistics of inferred model parameter values is obtained from 20 independent datasets. (Error bars indicate one standard deviation calculated across these 20 independent replicates.)

#### Windowed mHMM

To check that our windowed mHMM was capable of fitting time-varying data, we conducted statistical validation using simulated traces exhibiting various time-dependent trends in the bursting parameters. We studied three scenarios that mimicked ways in which bursting parameters could, in principle, be modulated to drive the onset of transcriptional quiescence: a decrease in *k*_on_ over time, an increase in *k*_off_ and a decrease in *r*. We also studied the case of increasing *k*_on_, as this was the strongest temporal trend observed in our experimental data. Appendix 4–Figure 6 summarizes the results for these validation tests.

For each test, 100 simulated traces, 40 minutes in length, were generated (Δ*τ* =20 s) that exhibited the desired parameter trends. Consistent with our approach to the experimental data, a sliding window of 15 minutes was used for inference, meaning that for each inference time, *τ*_inf_, all data points within 7.5 minutes of *τ*_inf_ were included in the inference. This led to inference groups consisting of 4500 data points, with the exception of the first and last time points, each of which had 3700 data points (first and last *w* + 1 points were excluded from inference). Transition and initiation rates shown in Appendix 4–Figure 6 are associated with state (1) of the three-state model (*k*_on_ = *k*_01_/2, *k*_off_ = *k*_10_ and *r* = *r*_1_ in Appendix 5–Figure 2A), as these were found to provide the most faithful indication of underlying system trends.

**Appendix 4 Figure 6.**
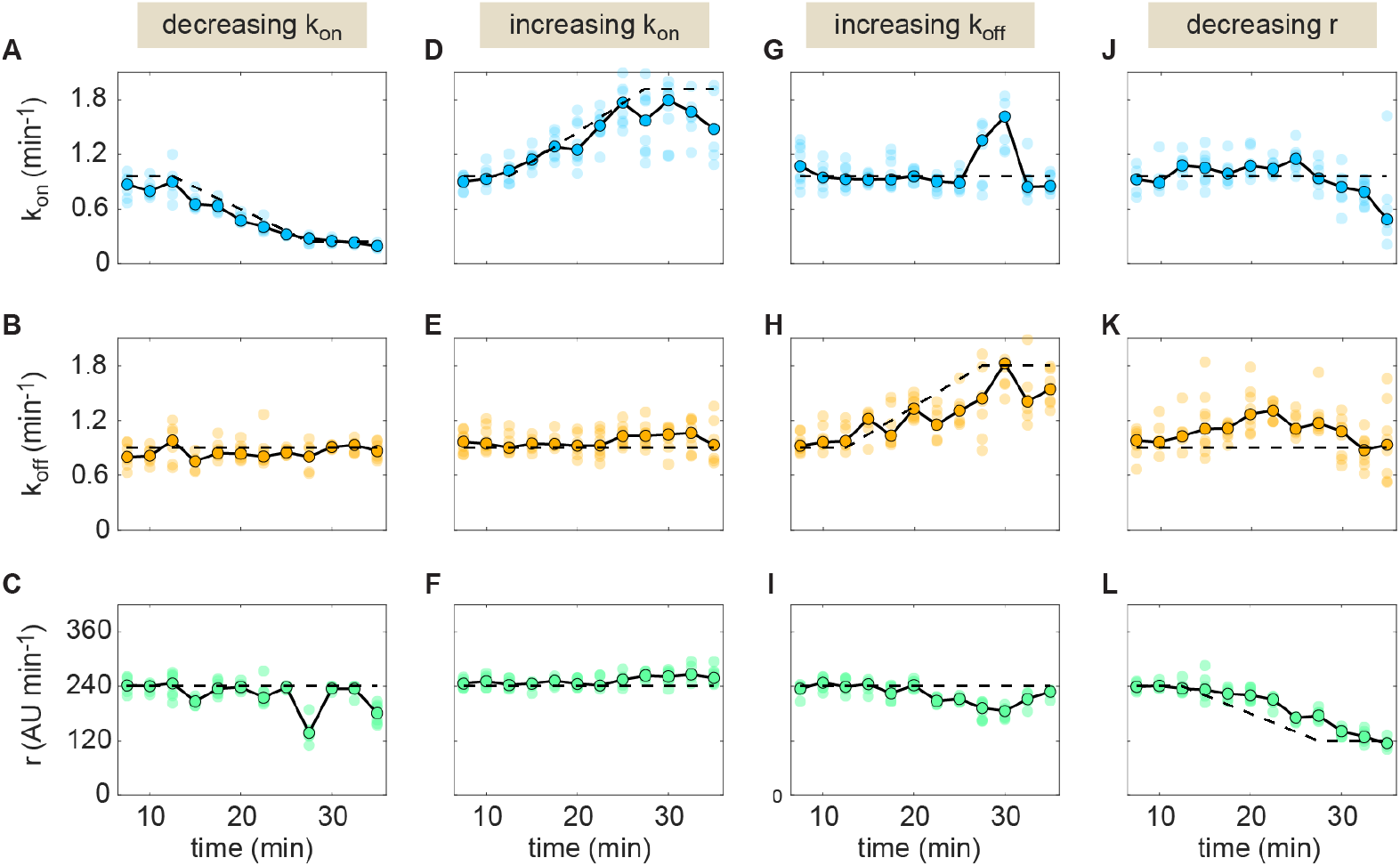
Validation of windowed mHMM inference. The method’s accuracy was tested for four distinct sets of parameter time trends. Results for each scenario are organized by column. In each plot, the black dashed line indicates the true parameter value as a function of time. Connected points (outlined in black) indicate the median inferred parameter value at each time point across 10 distinct replicates. Translucent points indicate inference values from individual replicates. Thus, the dispersion of these replicates at a given time point indicates the precision of the inference.

For each scenario, we assessed whether and to what degree the windowed mHMM method could accurately recover the temporal profiles. In general, the method was found to perform quite well within the parameter regimes that were tested. For both the increasing and decreasing *k*_on_ scenarios (Appendix4–Figure 6A-C,D-E), windowed mHMM inference accurately captured the modulation in *k*_on_ with no significant variation evident in the *r* and *k*_off_ trends. In the case of increasing *k*_off_ (Appendix4–Figure 6G-I), we observed deviations in *k*_on_ and *r* from their true values at the inflection point of the *k*_off_ curve (around 30 min). However, the deviation in *r* is relatively mild and the “blip” in *k*_on_, while of larger magnitude, is comprised of only two time points and so would likely not be mistaken for a legitimate indication of underlying system behavior. In the case of a decrease in the initiation rate (Appendix4–Figure 6J-L) we observe a ~ 5 min delay in the model response. We attribute this delay to the finite dwell time of RNA polymerase molecules on the gene (in this case *τ*_elong_ =140 sec, although further studies will be needed to determine why the observed lag appears larger than the elongation time). In addition, we note a degradation in the precision of the inference of *k*_on_ and *k*_off_ at low *r* (RHS of Appendix 4–Figure 6J, K).

Overall, we conclude that the windowed mHMM method is capable of accurately inferring time-resolved parameter values. An important caveat to these results is that the size of the sliding window (15 min in this case) places an inherent limit on the time scales of the parameter trends the model is capable of inferring. Changes that occur on shorter time scales will be registered, but the temporal averaging introduced by the sliding window will lead to underestimates of the rate of the parameter changes in the underlying system.

## Appendix 5

### Sister chromatids

#### Detection of sister chromatid appearance

Previous studies have indicated that the *D. melanogaster* genome is quickly replicated at the beginning of each nuclear cycle in early development (Rabinowitz, 1941; Shermoen et al., 2010), suggesting that each diffraction-limited spot in our imaging data likely contains two distinct *eve* loci. We sought use our live imaging data to verify whether genome replication occurred early enough in the nuclear cycle such that the presence of the replicated promoters would have to be taken into account. While the two *eve* loci are located within a diffraction-limited spot for the majority of frames in our data, there are a subset of frames in which two distinct puncta can be clearly observed due to fluctuations in the separation between chromatids (see Figure 4D). We reasoned that, by tracking the frequency of frames with resolved puncta over time, we could ascertain how the timing of genome replication compares to the onset of transcription. If replication precedes the onset of transcription, then the fraction of resolved frames should be relatively stable over for the duration of *eve* expression in nuclear cycle 14. If, on the other hand, replication happens after the onset of transcription, we should see a significant increase in the frequency of resolved sister chromatids over time as development progresses.

To pursue this question, we randomly selected snapshots of transcriptional loci in 100 different nuclei for each of the 11 embryos used in this study. We then determined the fraction of these sampled snapshots in which two distinct puncta were clearly visible by eye and observed how these instances of resolved chromatids were distributed in time. As indicated in Appendix 5–Figure 1A, we see evidence for resolved puncta by around 7 minutes into nuclear cycle 14. This is well within the average range for turn-on times observed throughout the stripe (see Figure 3–Figure Supplement 2B). Our results indicate that, at the very least, the genomic region containing our *eve* stripe 2 reporter is replicated within *some* nuclei by 6-8 minutes into nuclear cycle 14. Appendix 5–Figure 1B tracks the share of total observations for which we detected resolved puncta as a function of time. A systematic delay in DNA replication would be expected to result in a progressive increase in this metric over time. However, such a trend is not evident. While we see *no* resolved sister loci between 4 and 8 minutes (first point in the plot in Appendix 5–Figure 1B), this absence could be attributed to other factors at play early on in nuclear cycle 14. For example, part of this apparent lag could be attributable to the fact that loci are, on average, dimmer early on in the nuclear cycle, which could mask the presence of two *eve* loci by reducing the probability of both producing observable amounts of fluorescence at the same time. It is also possible that the precise timing of locus turn-on varies for each sister locus, as it does for loci in different nuclei. Regardless, even if the initial rise between 6 and 10 minutes in Appendix 5–Figure 1B is reflective of the replication of the locus during this period of time, the relative stability of the frequency of resolved loci from 10 minutes onward indicates that this process is restricted to the first few minutes of transcription. Additional experiments are needed to further elucidate the interplay between DNA replication and the onset of transcription. Regardless, the examination of our live imaging data supports the conclusion that the majority of our data consist of diffraction limited spots containing two distinct *eve* loci.

**Appendix 5 Figure 1.**
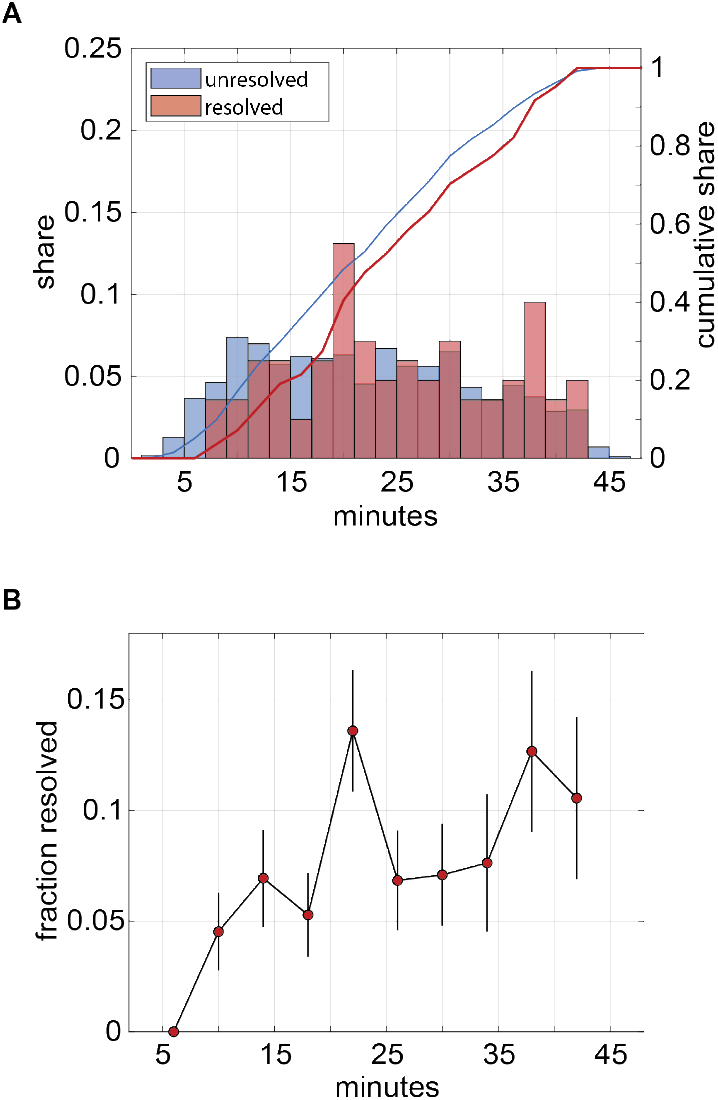
Live imaging data indicate timing of sister chromatid appearance. **(A)** Distribution of observation times for frames in which chromatids were resolveable (red) and diffraction-limited (blue). Bars indicate emprical probability distribution function. Lines indicate cumulative density function. Data indicate the presence of chromatids by no later than 7-8 minutes into nuclear cycle 14. **(B)** Fraction of frames featuring resolved chromatids as a function of time. Trend suggests replication of relevant portion of genome across all observed nuclei is completed by approximately 10 minutes into nuclear cycle 14. Inititial lag is likely attributable—at least in part—to stochastic turn-on times between sister *eve* loci and lower fluorescence levels early on in the nuclear cycle.

#### Probing for interactions between sister chromatids

If each fluorescent punctum contains two promoters (Figure 4D), then it is necessary to revisit the widely used two-state model of transcriptional bursting. In this revised scenario, each promoter on one of the sister chromatids undergoes fast ON/OFF switching. Therefore, each spot (encompassing two identical loci) can be in one of three states: (0) both promoters OFF, (1) one promoter ON and the other OFF, and (2) both promoters ON (Appendix 5–Figure 2B). States (1) and (2) are expected to exhibit different rates of RNA polymerase loading, *r*_1_ and *r*_2_, respectively. See Appendix 1 and Appendix 3 and for details regarding the implementation of this three-state model.

The presence of two transcriptional loci within each fluorescent punctum suggests three constraints on the relationship between bursting parameters in the model shown in Appendix 5–Figure 2A. First, if these two promoters transcribe independently, then state (2) will have double the loading rate of state (1) such that *r*_2_ = 2*r*_1_. Second, the probability of both promoters transitioning simultaneously should be negligible; we expect no transitions between states (0) and (2) such that *k*_02_ = *k*_20_ = 0. Finally, if the promoters switch between their states in an independent manner, then there will be an extra constraint on their transitions rates. For example, there are two paths to transition from (0) to (1) as either promoter can turn on in this case. However, there is only one possible trajectory from (1) to (2) because only one promoter has to turn on. This condition sets the constraint *k*_01_ = 2*k*_12_. Similarly, *k* = *k*_21_/2.

**Appendix 5 Figure 2.**
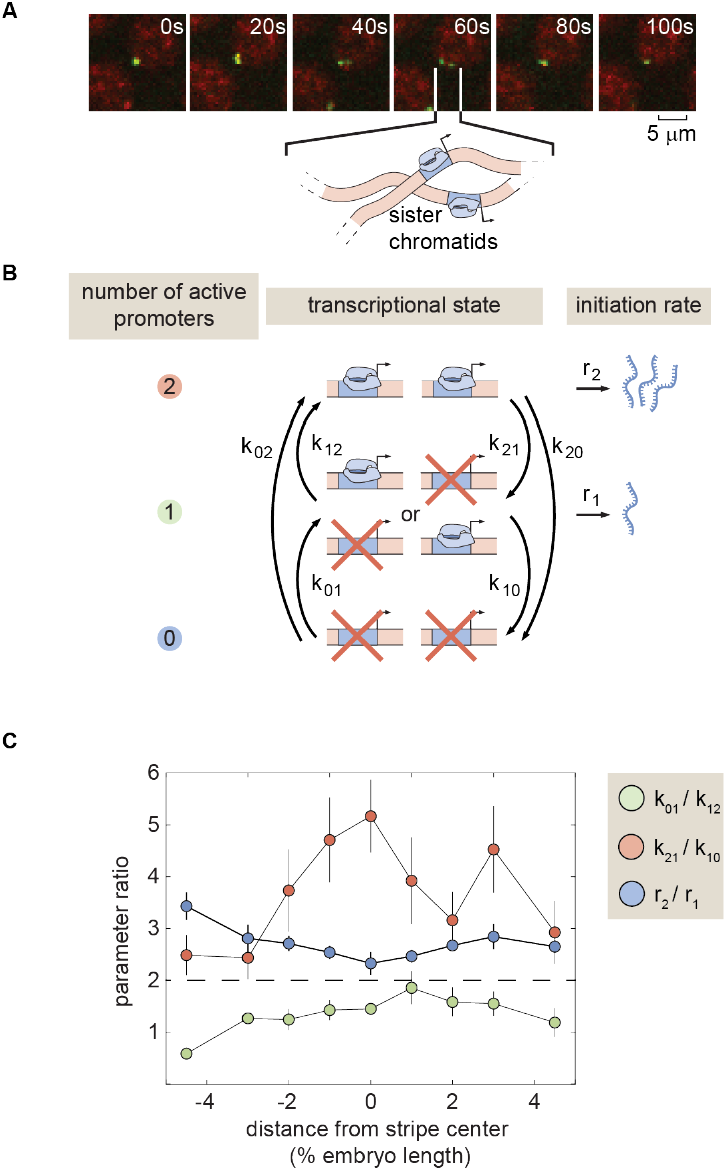
Probing combined transcription of sister chromatids. **(A)** Revised three-state model of promoter switching within a fluorescent punctum that accounts for the combined action of both sister chromatids. **(B)** Summary of bursting parameter ratios. All three bursting parameter ratios deviate from their expected values under the independence assumption given by the horizontal dashed line. (Error bars indicate magnitude of difference between first and third quartiles of mHMM inference results for bootstrap samples of experimental data over multiple embryos. See Materials and Methods for details)

While the independence of sister chromatids is supported by recent single-molecule FISH experiments (Little et al., 2011; Zoller et al.,2018), classic electron microscopy work suggests a scenario in which sister chromatids are tightly correlated in their transcriptional activity (McKnight and Miller, 1977, 1979). Given this uncertainty regarding chromatid independence, we elected to employ a general three-state model that makes no assumptions about the nature and strength of sister chromatid interactions. In addition to permitting greater flexibility, this agnostic approach also meant that the structure of the kinetic model returned by mHMM inference provided clues regarding the nature of the coupling between sister loci. Specifically, we examined the ratios between the high and low on rates (*k*_01_ and *k*_12_), off rates (*k*_21_ and *k*_10_), and initiation rates (*r*_2_ and *r*_1_). A deviation from these expectations would indicate either that the two sister loci do not initiate RNA polymerase independently (first constraint), or that they do not transition between activity states independently (second and third constraint).

Overall, our results suggest that the two loci are coupled to a nontrivial degree. We observe that the rate of initiation for the high state, *r*_2_(*x*), (corresponding to two active promoters) is consistently greater than twice the middle state, *r*_1_(*x*) (Appendix5–Figure 2B, bliue). This trend suggests some sort of synergy in the RNA polymerase initiation dynamics of the sister promoters. Even more strikingly, we observe that the rate of switching from (2) to (1), *k*_21_, is *much* higher than twice the rate of switching from (1) to (0), *k*_10_, (Appendix 5–Figure 2C, red). This indicates that each promoter is more likely to switch off when its sister locus is also active. This anti-correlation is consistent with some form of competition between the loci, a scenario that could arise, for instance, if local concentrations of activating TFs are limiting. In addition, we observe substantial variation in the relationship between the high and low on rates (*k*_01_ and *k*_12_, respectively), ranging from one of near equality in the anterior flank to nearly the 2-to-1 ratio that would be expected of independent loci in the stripe center and posterior (Appendix 5–Figure 2C, green). Finally, as shown in Appendix6–Figure 1, we observe no transitions between the (0) and (2) states, lending support to the hypothesis that, despite their correlation, our spots do contain two promoters.

Further experiments in which the sister chromatids are labeled in an orthogonal manner are needed to confirm and elaborate upon these results. One important consideration to address is the fact that the spatial proximity of the two loci appears to fluctuate significantly over time. Thus, if (as seems plausible) the strength of the coupling between loci depends in some way upon the radial separation of the loci, then the results reported here are effectively an average of time-varying system behavior. Valuable information may be obscured as a result of this averaging.

## Appendix 6

### mHMM inference sensitivities

#### Full three-state inference results

For the sake of simplicity, we presented our inference results in the main text using an effective two-state model in which two distinct active transcriptional states were combined into a single effective ON state (see Figure 4E and F). Here, for completeness, we include time-averaged and time-resolved inference results for the full three-state model where, as shown in Appendix 5–Figure 2, (0) corresponds to the state where both promoters are in the OFF state, (1) indicates the state where either promoter is in the ON state, and (2) represents the states where both promoters are in the ON state.

As indicated in the main text, the full three-state results (Appendix 6–Figure 1) exhibited the same trends as were evident in the effective two-state plots (Figure 5). In agreement with the effective two-state model, the rate of transcript initiation is not modulated to a significant degree across the stripe (Appendix 6–Figure 1D). Moreover, we once again see that activation rates, and specifically the rate of switching from OFF to the middle ON rate (states 0 and 1 in Appendix 6–Figure 1E) are strongly elevated in the stripe center.

**Appendix 6 Figure 1.**
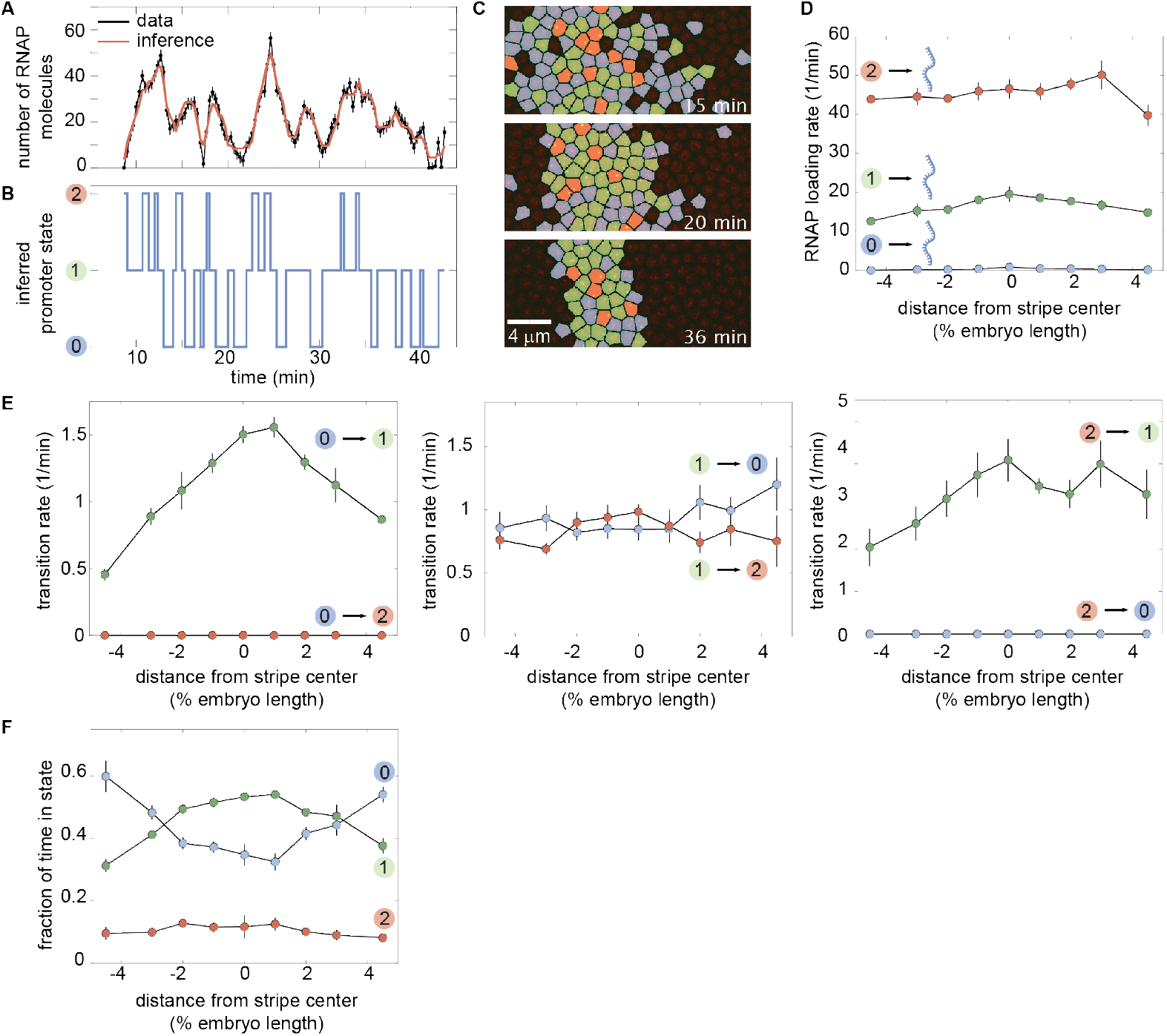
Full three-state results for time-averaged mHMM inference. **(A)** Representative experimental trace along with its best fit and **(B)** its most likely corresponding promoter state trajectory. **(C)** Instantaneous visualization of promoter state in individual cells throughout development through the false coloring of nuclei by promoter state (colors as in B). **(D)** The rate of initiation for each transcriptional state is not significantly modulated along the embryo. **(E)** Our mHMM revealed that the transition rate between the OFF (0) and middle ON state (1) is up-regulated in the stripe center. In contrast, the rates of switching out of the middle and high ON states show little to no significant AP-dependent modulation. **(F)** The modulation of the rate of switching from 0 to 1 acts to increase the fraction of time the promoter spends in the active states in the stripe center. (A, error bars obtained from estimation of background fluorescent fluctuations, as described in Materials and Methods and Garcia et al. (2013); D, E, and F, error bars indicate the magnitude of the difference between the first and third quartiles of mHMM inference results for bootstrap samples of experimental data taken across 11 embryos. See Materials and Methods for details.)

Like the time-averaged results, time-resolved inference trends for the full three-state model agree closely with effective two-state results shown in main text (compare Appendix 6–Figure 2 to Figure 6D-F). Due to a lack of statistics for state (2), we show only transition rates into and out of the first active state (middle state in Figure 4E).

**Appendix 6 Figure 2.**
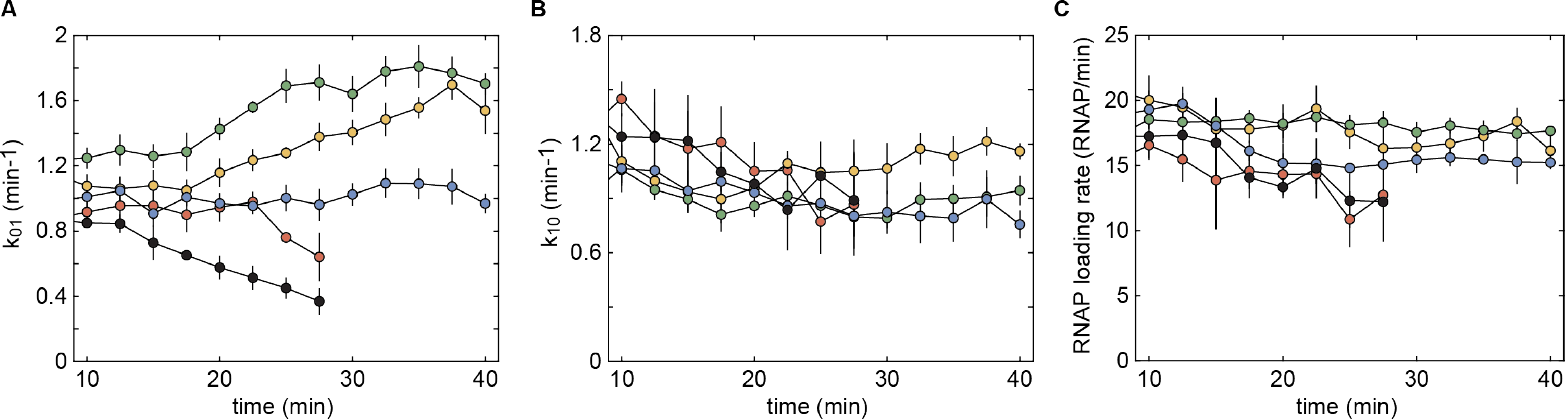
Full three-state results for time-dependent mHMM inference. **(A)** Transition rate from transcriptionally inactive state (0) to the first active state (1). Same trends evident as for effective 2 state model. **(B)** Transition rate from first on state (1) to OFF state (0). **(C)** Rate of transcript initiation in first on state (1) as a function of time. (Error bars indicate the magnitude of the difference between the first and third quartiles of mHMM inference results for bootstrap samples of experimental data taken across 11 embryos. See Materials and Methods for details.)

#### Two-state inference results

Although the presence of sister chromatids indicated that the three-state model was most appropriate for the *eve* stripe 2 system, we wanted to check that our conclusions were robust to this assumption. To do this, we conducted time-averaged and windowed inference assuming a simpler, two-state model (see, *e.g*. Figure 4B). Note that this approach is distinct from the *effective* two-state results presented in the main text. There, as outlined in Figure 4D-F, a three-state model was specified for inference and the results for the two active (ON) states were aggregated after the fact to simplify the presentation of the results. Conversely, here, we explicitly conducted inference using a two-state model.

Most of our findings remained unchanged in the context of the two-state model. Consistent with the three-state case, the two-state time-averaged mHMM inference indicated that the fraction of time spent in an active state, rather than the rate of RNA polymerase initiation, drives the difference in mRNA production rates across the stripe (Appendix 6–Figure 3A-C). Moreover, as with the three-state case, two-state results indicated that the bulk of this variation stem from modulation in *k*_on_ (Appendix 6–Figure 3C, green). Interestingly, whereas we did see a degree of spatial dependence in *k*_off_ for 3-states, we observed no such trend for 2-states (Appendix 6–Figure 3C, red). In general, this is not surprising, as our use of a simpler model likely means that multiple switching rates are being projected onto the *k*_off_ parameter. Specifically, if the *eve* stripe 2 system is indeed a true three-state system, then we would expect the two-state *k*_off_ estimate to reflect the joint action of the *k*_10_, *k*_21_, and *k*_12_ rates from the three-state model. As a result, the spatial dependence of each one of these rates would get averaged out when combined onto *k*_off_.

**Appendix 6 Figure 3.**
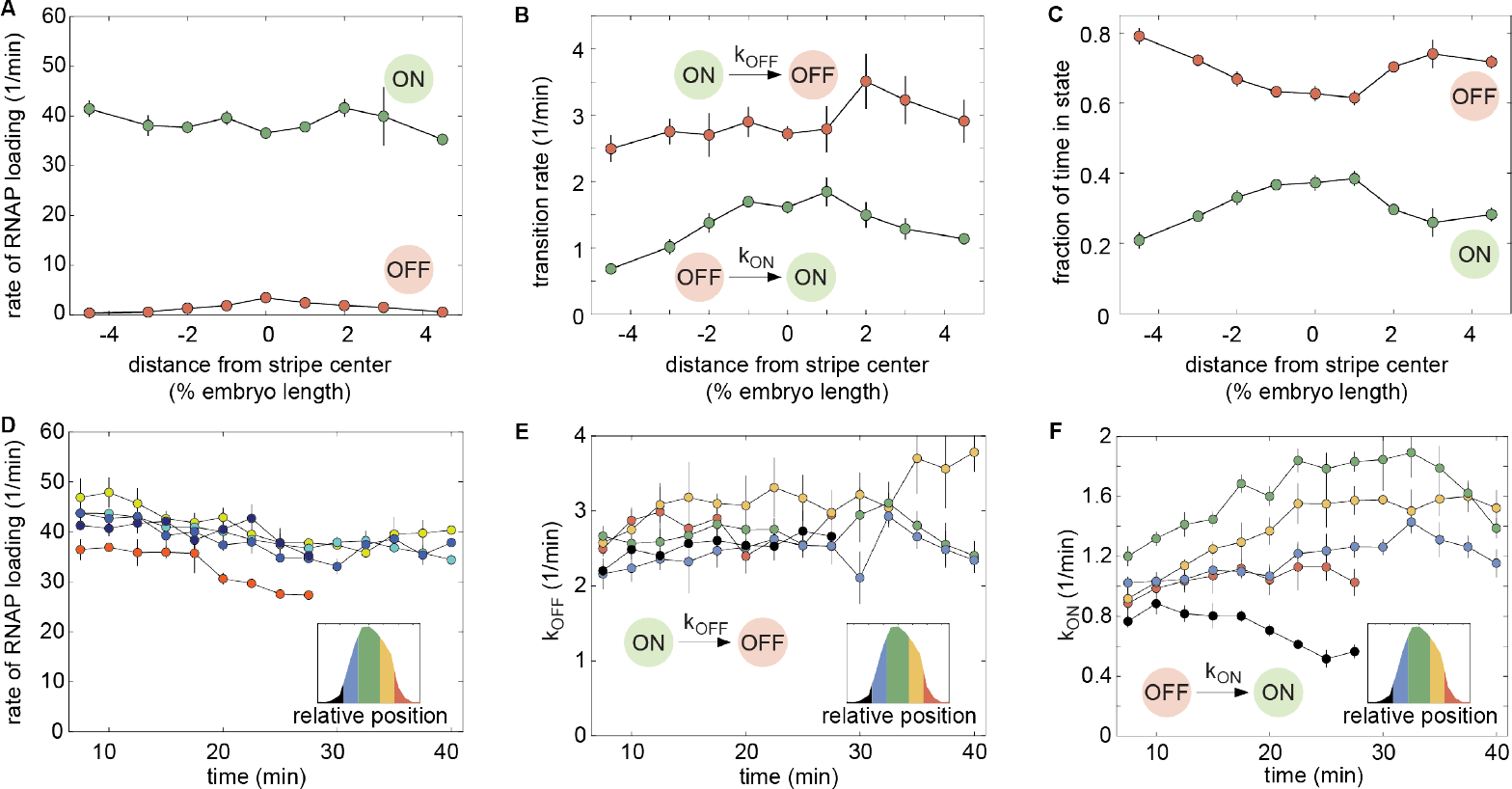
Two-state mHMM inference. **(A-C)** Time-averaged 2-state inference results. (A) Consistent with three-state inference results, we observed no significant modulation in the rate of initiation along the axis of the embryo. Moreover, we found that *k*_on_ (green plot in (B)) was modulated along the anterior-posterior axis to vary the amount of time the promoter spent in the ON state (green curve (C)). In a departure from the three-state case, we observed no significant spatial trend in *k*_off_, though we noted a spike in *k*_off_ at 3% of the stripe center. **(D-F)** Time-resolved (windowed) two-state mHMM results. (D) Consistent with the 3-state inference, we saw little to no modulation in the rate of RNA polymerase loading *r* over time, although we noted a mild downward trend across all AP bins that was most pronounced in the posterior flank (red curve). (E) Two-state inference indicated no significant temporal trends in *k*_off_. (F) *k*_on_ time trends largely agreed with the three-state case, although we noted that the decrease in *k*_on_ in the posterior flank that was apparent in the three-state results was not observable in this two-state context (Figure 6E, red). (Error bars indicate the magnitude of the difference between the first and third quartiles of mHMM inference results for bootstrapped samples of experimental data. See Materials and Methods for details.)

As with the time-averaged case, we found that results for two-state windowed mHMM were generally consistent with three-state trends. A notable exception to this rule was the absence of any significant decrease in *k*_on_ in the posterior stripe flank (Appendix 6 Figure 3F, red). This is not entirely surprising, as the trend returned by the three-state inference was relatively mild (Figure 6E, red), encompassing only the final two time points for which there was sufficient data to conduct inference. It is possible that the added complexity of the three-state model allowed it to register a subtle shift in the activation rate that was convolved with countervailing features in the two-state case. Future work will seek to elucidate the source of this discrepancy and further test the validity of the trend uncovered in the three-state case.

#### Comparing true and effective two-state inference results

Here, for completeness, we provide direct comparisons between the time-averaged inference for the effective two-state results presented in the main text and the true two-state results presented in the previous section.

**Appendix 6 Figure 4.**
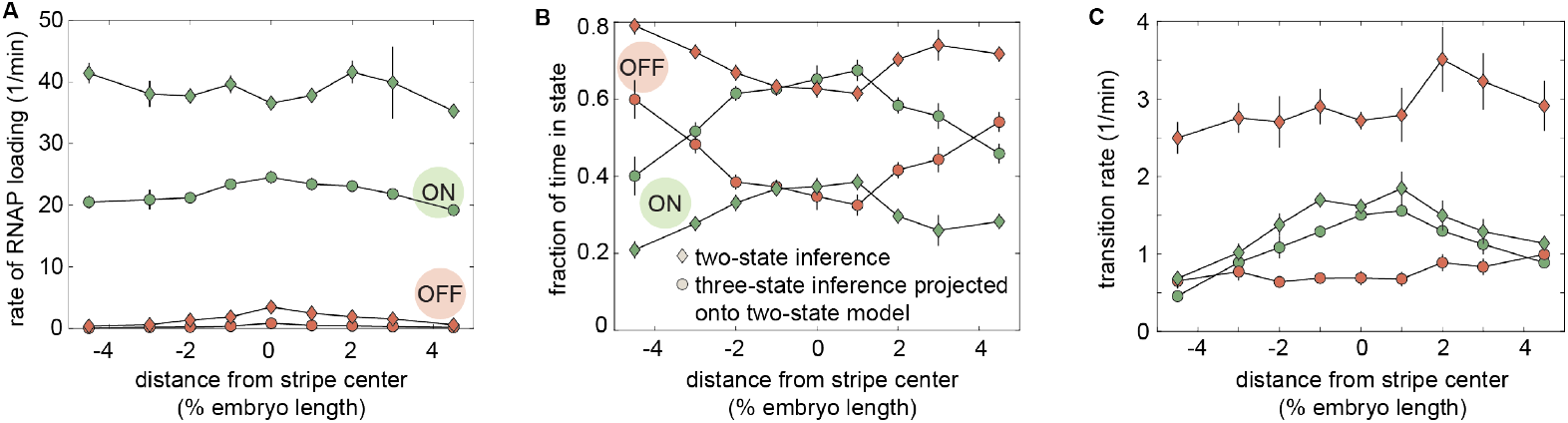
Comparing two- and three-state mHMM inference results. Three-state inference results can be presented in terms of a two-state model in which states (1) and (2) are aggregated into a single ON state (see Figure 4E and F). Here, color schemes are consistent with those employed in Appendix 6–Figure 3A-C. Squares indicate true two-state results (presented in the previous section) and circles indicate effective two-state trends derived from the three-state results presented in Figure 5. **(A)** Anterior-posterior-dependent trends in the rate of RNA polymerase initiation are nearly identical between the true and effective initiation rates, however the initiation rate returned by two-state mHMM inference (green squares) is roughly twice as large as that implied by the three state results (green circles). **(B)** As with the initiation rates, we observe similar trends between the true and effective cases, but substantial differences in magnitude. The effective two-state model recovers an ON state occupancy that is roughly double that returned by two state mHMM inference. **(C)** While the ON rate trends and magnitudes are nearly identical, the OFF rate returned by two-state mHMM inference is roughly triple that implied by three-state inference. Thus it is clear that this difference in OFF rate underlies the observed departures in both state occupancies (B) and state initiation rates (A). (Error bars indicate magnitude of the difference between the first and third quartiles of mHMM inference results for bootstrap samples of experimental data. See Materials and Methods for details.)

As Appendix 6–Figure 4 makes clear, while anterior-posterior-dependent parameter trends are by and large consistent between the true and effective two state models, we do observe substantial differences in the absolute magnitudes of parameter values. These differences originate (directly or indirectly) from the three-fold difference in the value of *k*_off_ between the true and effective models (Appendix 6–Figure 4C, red squares and circles, respectively). The *k*_off_ value for the effective two-state model is defined as

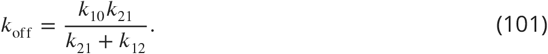

See Appendix 1 for expressions for all three effective two-state bursting parameters (*k*_on_, *k*_off_, and *r*) in terms of these three-state transition rates. This value represents the inverse of the mean amount of time the system, upon switching out of state (0), spends in one of the active states before returning to (0), and we can see that it is necessarily less than or equal to *k*_10_.

Thus, the two- and three-state results imply that the systems switch out of the active state(s) on substantially different timescales. On the other hand, the ON rates are strikingly similar across the two models. As a result, the effective two-state model implies that the system is in one of the active states for between 40 and 70% of time, whereas two-state mHMM inference implies significantly lower shares falling between 20 to and 40%. Since both models must reproduce the same mean production rate—this is an inherent feature of the experimental traces—we see that the two-state mHMM inference returns an estimated initiation rate that is consistently twice as large as the initiation rate implied by the effective two-state model.

Thus, while most of the conclusions featured in this paper are robust to our choice of model architecture, this decision does, nonetheless hold important implications for how we understand the underlying system. Further work is needed elucidate the root cause of this discrepancy and move towards a more concrete understanding of the correspondence between the structure of the model and that of the physical system.

## Appendix 7

### Inherent limits of bursting parameter inference

By definition, the onset of transcriptional quiescence coincides with the cessation of observable bursting activity. In the main text, we argue that this cessation appears to be driven by processes that are mechanistically distinct from those driving transcriptional bursting. It remains possible, however, that quiescence is instead driven by changes in the bursting machinery itself as illustrated in scenario (ii) in Figure 6A. If this is the case, it is important to note that fundamental limits exist to the time-scale of shifts in the bursting parameters that could be detected in any sort of time-dependent burst parameter inference (see, *e.g*., Figure 6): changes of order-with or faster-than the time scale on which transcriptional bursts occur (1-3 min for *eve* stripe 2) cannot be detected. Notably, this is not a limit of the mHMM method, but rather reflects an inherent limitation set by system itself—in order to infer bursting parameters, we must observe bursts and, in order to infer a change in parameters, we must have access to bursting activity that reflects this change. Thus, the characteristic frequency of bursts sets a resolution limit for any kind of bursting parameter inference.

To illustrate this limitation, we simulated three scenarios for a two-state transcriptional system in which *k*_on_ decreases to 0 *s*^−1^ over periods 15, 5, and 1 min in length. We then sought to recover the trend in *k*_on_. To emphasize that the limitations are not specific to mHMM, we used the *true* promoter trajectories generated by our simulation algorithm to estimate *k*_on_. These estimates thus represent the absolute best-case scenario for parameter inference, in which we recover the underlying behavior of the system exactly. The results indicate that, as expected, a transition in *k*_on_ that happens in the span of 1 minute is not detectable from a burst inference perspective (Appendix 7–Figure 1A-C). This indicates that, at this timescale, a shift in burst parameters (scenario ii in Figure 6A) would be indistinguishable from an abrupt, change in which the promoter entered a silent state outside of those considered by the bursting model (scenario (i) in Figure 6A). Interestingly, results for 5 and 15 minute *k*_on_ transitions [Appendix 7–Figure 1D-I) also indicate that even transitions that occur over longer periods of time cannot be fully recovered due to the fact that bursting behavior is observed over a limited window of time (40 minutes in our case). Thus, once the burst frequency decreases to a sufficiently low level, there simply are not enough bursts observed within the window of observation to estimate the burst frequency from the data.

**Appendix 7 Figure 1.**
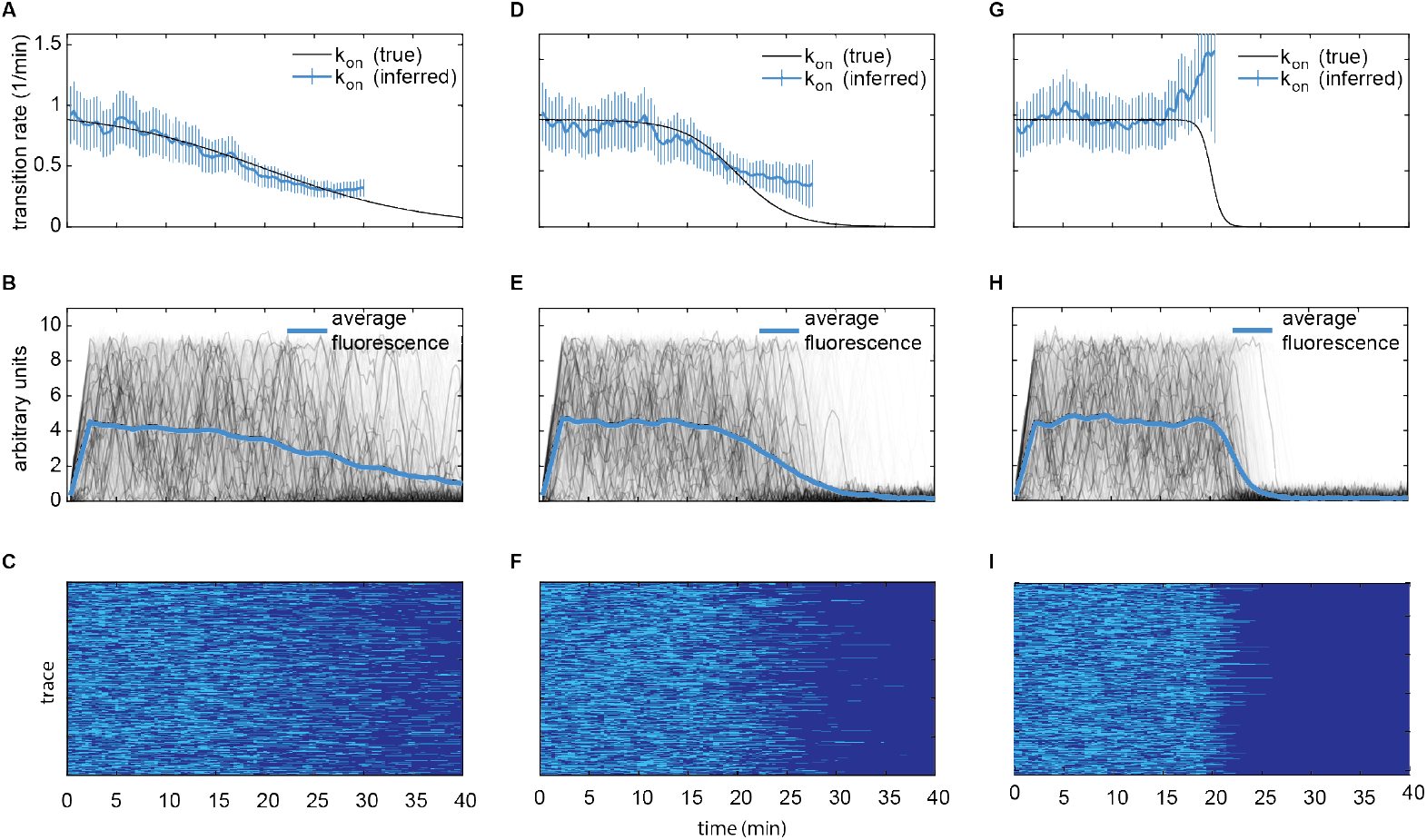
Inherent limits of bursting parameter inference. **(A-C)** Simulating a 1 min transition in *k*_on_. **(A)** Black curve indicates true *k*_on_ value as a function of time and blue curve indicates inferred value. Because the change unfolds on a time scale that is much faster than the bursting timescale, it is not possible to accurately recover the underlying *k*_on_ trend from the fluorescent traces. **(B)** The temporal trend in the average fluorescence across simulated traces (blue curve) reflects this fast decrease in *k*_on_. Note that variation in simulated traces (gray) unfolds on a significantly faster timescale than the change in the mean. **(C)** Visualization of promoter switching showing how the *k*_on_ transition occurs on the timescale of a single burst. Light blue indicates ON periods and dark blue indicates OFF periods. Since there are almost no active traces after the transition of *k*_on_ to perform an inference. it would be impossible to determine if a modulation in the bursting parameters—as opposed to a transition into some alternative, silent state—drives the onset of quiescence. **(D-F)** Simulation of a 5 min transition in *k*_on_. **(D)** We are able to recover first half of *k*_on_ trend, but due to the speed of transition, insufficient active traces remain to permit the accurate recovery of the full profile. **(E,F)** Because the transition happens slower than in the 1 min case shown in (A-C), there are some bursts that unfold during the transition and, hence, we have some reference points with which to infer the underlying trend. **(G-I)** Simulating a 15 min transition in *k*_on_. **(G)** The mHMM can reliably infer the temporal variation in *k*_on_. **(H,I)** The observation that bursts of activity are interspersed throughout the *k*_on_ transition makes it possible to recover the temporal trend. (A,D,G, error bars indicate 95% confidence interval of exponential fits used to estimate *k*_on_).

## Appendix 8

### Input-Output analysis details

In this appendix, we provide additional information about data sources, inference methodology, and inference sensitivies related to the input-output analysis presented in the main text.

#### Input transcription factor data

##### Data sources

The input-output analysis presented in the main text made use of previously published data sets for the spatiotemporal concentration profiles of the gap genes Hunchback, Krüppel, and Giant (Appendix 8–Figure 1A, C and D). These data derive from elegant experiments in which individual embryos were co-immunostained for transcription factors of interest and precisely staged by measuring progressive cellularization over the course of nuclear cycle 14 to generate a time series of protein concentration profiles spanning the course of this period of development (Dubuis et al., 2013). The Bicoid concentration data used for this analysis derives from live imaging experiments using a Bicoid-GFP fusion established by Gregor et al. (2007). These data come courtesy of Jonathan Liu and Elizabeth Eck (Appendix 8–Figure 1B).

**Appendix 8 Figure 1.**
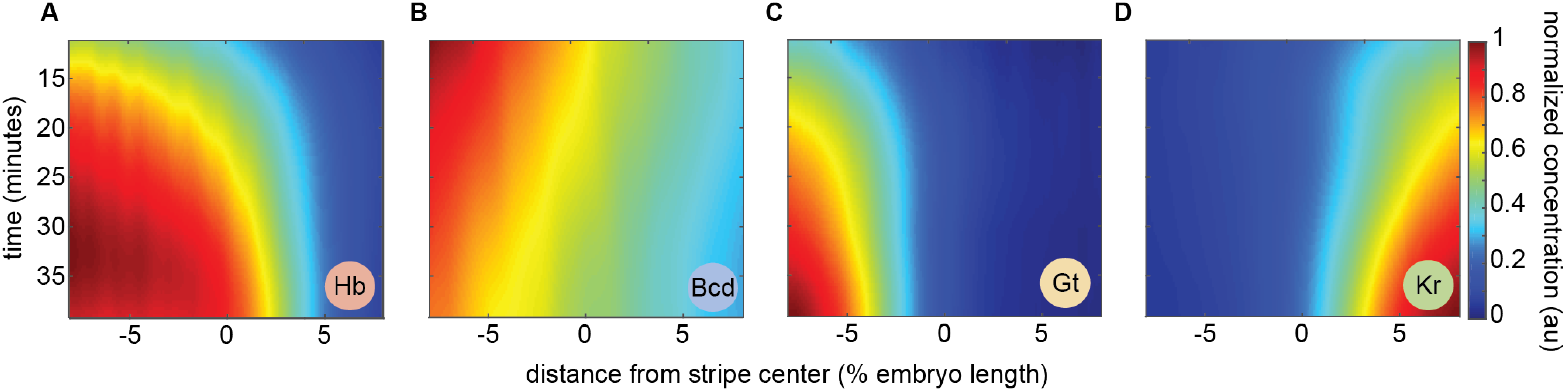
Spatiotemporal transcription factor concentration maps. Heatmaps indicate normalized concentration profiles for the *eve* stripe 2 regulators **(A)** Hunchback, **(B)** Bicoid, **(C)** Giant, and **(D)** Krüppel as a function of space and time. In each case, levels were normalized relative to the maximum concentration observed within the spatiotemporal window of interest.

##### Data processing

To prepare the Krüppel, Giant, and Hunchback profiles for use in our logistic regression analysis, we adopted an approach similar to that described in Dubuis et al. (2013). Dorso-ventral orientation of embryos was found to have negligible effect on calculated intensity profiles and was ignored (i.e. all embryos were included, regardless of orientation). For each time point in nuclear cycle 14, a weighted temporal average was calculated using a sliding Gaussian kernel with *σ_t_* = 5 min. For each time point, the minimum observed value across all anterior-posterior positions was then calculated and subtracted in order to remove background fluorescence. Normalized profiles were then calculated using the formula

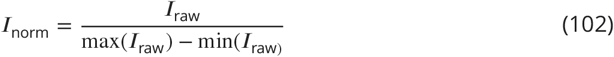

An identical procedure was followed for processing the Bicoid-GFP data, with the addition of a spatial averaging step using a sliding Gaussian window of *σ*_AP_ = .5 % embryo length. This step was necessitated by the fact that, because individual embryos were imaged for the duration of nuclear cycle 14, multiple experiments contributed concentration data along the anterior-posterior axis for each time point. Thus averages in both space and time were needed in order to effectively aggregate these data into a single average spatiotemporal profile.

Finally, we discovered that the anterior-posterior axes in our live imaging data (both for *eve* stripe 2 and Bicoid-GFP) were inconsistent with the axes employed by the fixed data reported by the authors in Dubuis et al. (2013). We addressed this issue by using *eve* stripe 2 as a fiduciary mark to register the positions of the fixed and live data sets. Specifically, we aligned the mRNA peak predicted by our model at 40 minutes into nuclear cycle 14 with the peak in second stripe of the *eve* protein profile at 40 minutes, as reported in Petkova et al. (2019).

#### Logistic regression framework

The binomial logistic regression is a widely used statistical method for assessing the relationship between a set of predictor variables and a response variable of interest that is constrained to take on one of only two possible outcomes. In the context of our analysis, the predictor variables were the normalized transcription factor concentration profiles and the response variables were (i) the overall transcriptional state given by the transcriptional time window (active or silent?) and (ii) the bursting state amongst trancriptionally active loci (ON or OFF?). Inference was conducted at the level of individual gene loci. fmincon, a standard matlab function for constrained optimization, was used to fit all models discussed both in the main text and in this appendix.

To prevent overfitting at the stripe centers, the selection of data sets for input-output inference were weighted to ensure equal representation of data points from across all regions of space and time included in the analysis. The data were divided into cells of size 1% of the embryo length in width and 1 minute in duration for the purpose of calculating and assigning these weights. The number of data points in adjacent regions were factored into each region’s weight score using a 2D Gaussian averaging kernel. Regions with fewer than 25 total data points were not included in the inference.

##### Inference details: transcriptional time window

For the time window input-output analysis, we considered only loci that were transcriptionally active for one or more time steps in nuclear cycle 14. Loci were classified as transcriptionally active for all time points between the first and last time points for which they exhibited detectable levels of transcriptional activity and silent for all time points following their final shut-off for which their nuclei were still present in the experimental field of view. Time points preceding the onset of activity were discarded. Appendix 8–Figure 2A illustrates how this quantity varies over space and time in our experimental data. We considered a class of logistic regression models in which each transcription factor was permitted to appear at most once, thus requiring that each factor act on *eve2* in a uniform manner through space and time; i.e., the same protein could not activate expression on one stripe flank and repress on the other.

##### Inference details: transcriptional bursting

The bursting input-output analysis focused exclusively on transcriptionally engaged loci. The Viterbi algorithm was used to infer the instantaneous activity state (ON vs. OFF) for all loci. This activity state was taken as the response variable in our regression analysis. In all other respects, the inference procedure was identical to that conducted for the time window.

#### Results of unconstrained inference

For the input-output inference results presented in the main text (Figure 7), we used prior knowledge about the regulatory function of each input transcription factor to constrain its range of permissible values in our inference. Specifically, we constrained the activators Bicoid and Hunchback to play activating roles in our model and, likewise, required that the repressors Krüppel and Giant played repressing roles. In several cases, this constrained inference led to models in which one or more transcription factors played no significant regulatory role (Bicoid and Hunchback for the time window and Bicoid for transcriptional bursting). In this section, we tested the sensitivity of the conclusions presented in the main text to our use of functional constraints by conducting unconstrained input-output inference runs.

##### Transcriptional time window

The results of our unconstrained input-output inference for the transcriptional time window are identical to those presented in the main text. Despite the fact that no limitations were imposed on the regulatory function of each factor, we nonetheless recovered a model in which the two repressors, Giant and Krüppel, are necessary and sufficient to explain the onset of transcriptional quiescence in the stripe flanks. In agreement with the constrained case, we found that the addition of Hunchback and Bicoid to this two-repressor model had no qualitative effect on the output profile predicted by the model (Appendix 8–Figure 2B). A quantitative comparison of model fit scores confirmed that the addition of Hunchback and Bicoid did nothing to improve model fit (Appendix 8–Figure 2C). Thus, we conclude that our finding that the transcriptional time window can be explained entirely by the joint repressive action of Krüppel and Giant is insensitive to our choice to impose functional constraints.

**Appendix 8 Figure 2.**
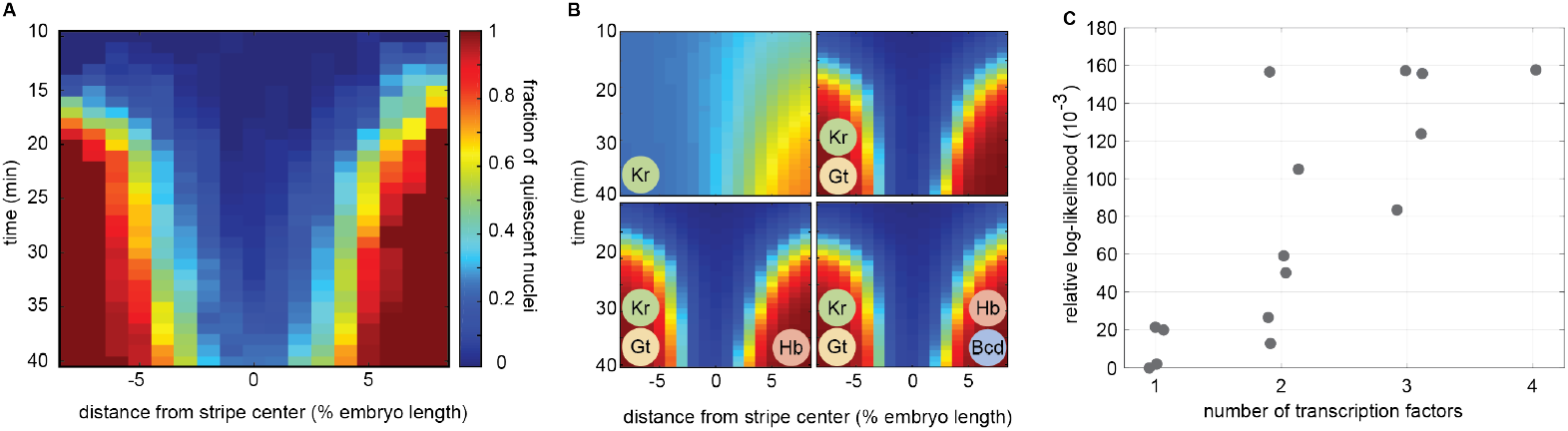
Unconstrained inference results for the transcriptional time window. **(A)** Observed fraction of quiescent nuclei as a function of space and time. Identical data to that presented in Figure 7A. **(B)** Relaxing constraints on the functional nature of each transcription factor had no appreciable effect on the inference results. Profiles shown here are indistinguishable from those shown in Figure 7D. Once again, we find that the joint action of the repressors Giant and Krüppel is sufficient to explain the progressive onset of transcriptional quiescence in the stripe flanks. **(C)** A quantitative comparison of model fits reinforces the qualitative conclusions drawn from (B). Models including 3 and 4 transcription factors cannot improve on the fit achieved by the simpler double repressor model. Here blue dots indicate models for which only Giant and Krüppel make significant contributions to the model fit. This indicates that, while the 3 and 4 transcription factor models include additional parameters, these do not contribute appreciably to overall model fit, emphasizing the fact that these models behave, effectively, as double repressor models.

##### Transcriptional bursting

In the context of the transcriptional bursting input-output analysis, the removal of functional constraints led to a significantly more complex landscape of inferred regulatory models. While the functional roles of Krüppel, Giant, and Hunchback were consistent with the constrained case (repressing, repressing, and activating, respectively), Bicoid was consistently inferred to play a repressing role. Despite this complication, the three-factor Krüppel-Giant-Hunchback model favored by the constrained inference remained the best-fitting three-factor model (Appendix 8–Figure 3C, red circle). While the addition of Bicoid as a repressor to create a model dependent on all four input transcription factors led to a small improvement in model fit (Appendix 8–Figure 3C), comparison of this four-factor model’s predicted activity profile with that of the Krüppel-Giant-Hunchback model revealed no material improvement in the model’s agreement with the experimental data (Appendix 8–Figure 3B, bottom left vs. bottom right). Moreover, there is (to our knowledge) no experimental evidence for Bicoid playing a repressive role in the regulation of *eve* stripe 2. Indeed, there is strong evidence that Bicoid is necessary for *eve* stripe 2 activity (Small et al., 1992). We thus conclude the Krüppel-Giant-Hunchback model remains the most plausible option in the unconstrained case.

**Appendix 8 Figure 3.**
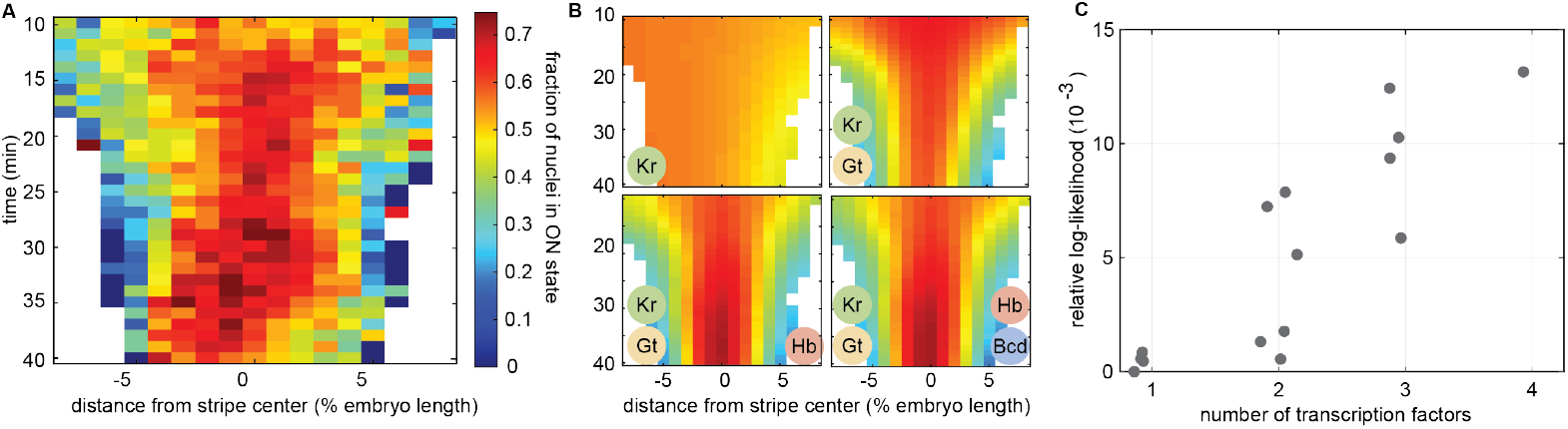
Unconstrained inference results for transcriptional bursting. **(A)** Observed fraction of transcriptionally active nuclei in the ON (bursting) state. Identical data to that presented in Figure 7B. **(B)** As with time window, relaxing the constraints on the functional nature of each transcription factor did little to alter the inference results presented in the main text (compare to Figure 7E). As with the constrained results, the joint action of Giant, Krüppel, and Hunchback appears sufficient to explain the spatiotemporal activity pattern revealed by mHMM inference. **(C)** A quantitative comparison of model fits.

## Appendix 9

### Determining the RNA polymerase dwell time using autocorrelation

In order to conduct mHMM inference, it is necessary to specify the number of time steps *w* required for an RNA polymerase molecule to traverse the reporter gene,

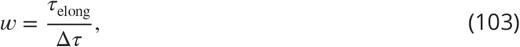

where Δ*τ* is set by the temporal resolution of our data acquisition and *τ*_elong_ is the elongation time which is unknown *a priori*. Past studies have estimated elongation rates for other systems involved in early patterning in the *Drosophila* embryo, but there is substantial disparity between the reported values. A live imaging study of transcriptional activity driven by the *hunchback* P2 enhancer reported an elongation rate of 1.4 – 1.7 kb min^−1^ (Garcia et al., 2013). However, a recent study of the same regulatory element reported elongation rates of 2.4 – 3.0 kb min^−1^—nearly twice as fast (Fukaya et al., 2017). These results suggested that RNA polymerase elongation rates measured for other systems might not apply to our *eve* stripe 2 reporter. Thus, in order to ensure the validity of our inference, we developed an approach that uses the mean autocorrelation function of experimental fluorescence traces to estimate the elongation time directly from our data.

The autocorrelation function *R_F_*(*τ*) quantifies the degree to which a signal, *F*(*t*), is correlated with a lagged version of itself, *F*(*t* – *τ*), and is given as a function of the time delay, *τ*, between the two signal copies being compared such that

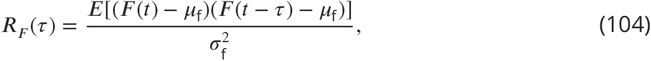

where *μ*_f_ is the average observed fluorescence, *σ*_f_ is the standard deviation of the fluorescence and *E* denotes the expectation value operator. As illustrated in Appendix 9–Figure 1A, the fact that it takes RNA polymerase molecules some finite amount of time to traverse the gene implies that the observed fluorescence at a transcriptional locus at some time *t*, *F*(*t*), will be correlated with preceding fluorescence values *F*(*t* – *τ*) so long as *τ* < *τ*_elong_ because the two time points will share a subset of the same elongating RNA polymerase molecules. As *τ* increases, the correlation between *F*(*t*) and *F*(*t* – *τ*) due to these shared RNA polymerase molecules will decay in a linear fashion (since the average number of shared RNA polymerase molecules decreases linearly with *τ*) until it reaches zero when *τ* = *τ*_elong_ (Appendix 9–Figure 1B, blue curve).

**Appendix 9 Figure 1.**
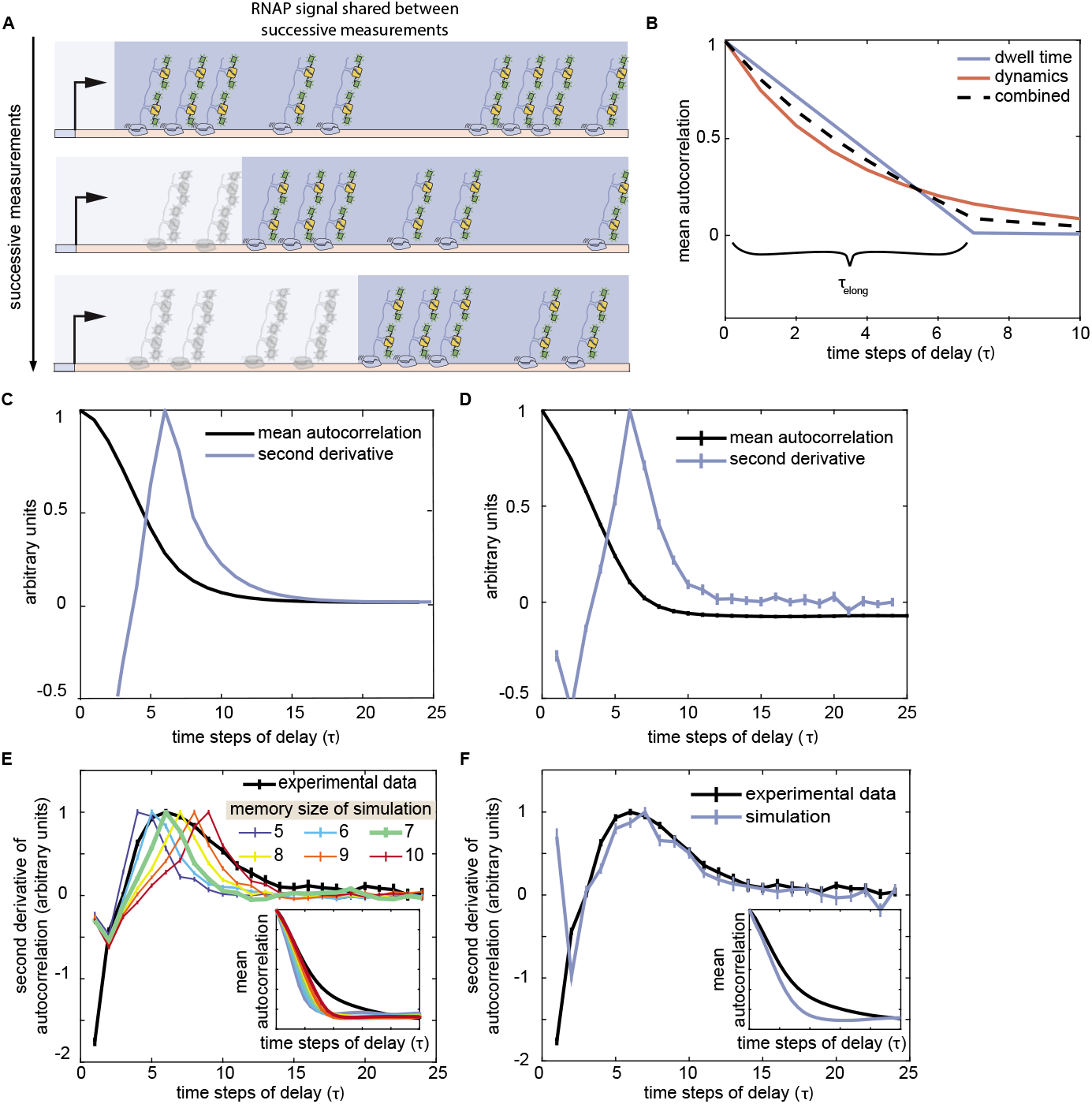
Using the autocorrelation of the fluorescence signal to estimate RNA polymerase dwell time. **(A)** It takes a finite amount of time for RNA polymerase molecules to transcribe the full length of the reporter gene. As a result, successive fluorescence measurements will contain some of the same GFP-tagged RNA polymerase molecules. Dark blue-shaded regions indicate the subset of RNA polymerase molecules that are present on the gene for successive measurements. **(B)** This overlap causes successive measurements to be correlated, and the degree of correlation due to the overlap decays linearly, reaching zero when the separation between measurements is equal to the elongation time, *τ*_elong_ (blue curve). However, the trace autocorrelation function contains other signatures that can obscure the inflection induced by RNA polymerase elongation dynamics. For instance, successive time points also exhibit correlation due to the promoter switching dynamics (red curve). **(C)** Theoretical analysis of the autocorrelation function and **(D)** stochastic simulations indicate that the second derivative of the mean autocorrelation function (dark blue curves) can be used to find the structural break in the function (black curves) that corresponds to *τ*_elong_. Here, a peak at 6 time steps of delay indicates an elongation time of 7 times steps (140 s). **(E)** Simulated traces with elongation time of 7 time steps (green curve) exhibit a peak in the second derivative that coincides with the maximum of the experimental curve. Inset plots show corresponding mean autocorrelation curves for experimental data and simulations. **(F)** Stochastic simulations in which we allow for variation in elongation times distributed around a mean of 7 time steps qualitatively recapitulates the observed curve. (C-F, second derivative profiles depicted here are normalized relative to their maximum value for ease of depiction.)

The dramatic change in the slope of the autocorrelation function that occurs at *τ* = *τ*_elong_ can be used to estimate the elongation time of the system; however, it is not the only feature present in Equation 104. Because the time series of promoter states constitutes a Markov chain, the instantaneous promoter state and, therefore, the instantaneous rate of RNA polymerase loading, exhibits a nontrivial, positive autocorrelation due to the promoter switching dynamics of the system. For instance, if it takes the promoter an average of 1 minute to switch states, then it is clear that promoter activity for *τ* < 1 min will be strongly correlated with itself. Thus, we see that the rates of promoter switching dictate the speed with which this “dynamic” autocorrelation decreases with increasing *τ*. More precisely, the dynamics autocorrelation will take the form of a decaying exponential in *τ*, with the time scale set, approximately, by the second largest eigenvalue of the Markov chain’s transition rate matrix (Appendix 9–Figure 1B, red curve)

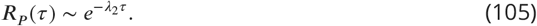

Where *λ*_2_ denotes the second larrgest eigenvalue of the transtion rate matrix. Thus, the observed autocorrelation function contains, at a minimum, signatures of both the finite RNA polymerase dwell time (*τ*_elong_) and of promoter switching dynamics. As a result, inferring elongation times from the change in slope in the mean autocorrelation is often relatively subtle in practice.

A theoretical analysis of *R_F_*(*τ*) indicated that the second derivative of the mean autocorrelation function reliably exhibits a peak that can be use to read out the value of *τ*_elong_ (Medin et al., 2019). Appendix 9–Figure 1C shows the analytic prediction for the autocorrelation and second derivative when *τ*_elong_ is equal to 7 time steps (*w* = 7). We confirmed that the same second derivative approach works in the context of stochastic simulations using realistic parameters for the *eve* stripe 2 system (Appendix 9–Figure 1D). These simulated traces included the expected contributions from both the Markov dynamics (red profile in Appendix 9–Figure 1B) and the finite RNA polymerase dwell time (blue profile in Appendix 9–Figure 1C). Having confirmed the efficacy of the autocorrelation method for simulated data, we next applied the same technique to uncover *τ*_elong_ for our experimental traces.

The black profile in Appendix 9–Figure 1E indicates the form of the autocorrelation second derivative for the set of traces used for mHMM inference. We observed that, while there is a definite inflection point, the peak for the experimental data is much broader than for simulated traces. The most likely cause of this feature is the existence of variability in *τ*_elong_ (see below). From comparisons of the position of the second derivative peak for experimental traces with simulated profiles, we concluded that an elongation time of *w* = 7 (*τ*_elong_ = 140 s) best characterized our data (Appendix 9–Figure 1E, green curve). This implies that

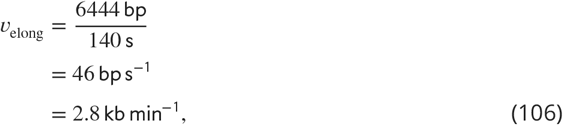

where the length used represents the distance from the start of the MS2 step loop sequence to the end of the 3’ end of the construct. Interestingly, this elongation rate falls within the 2.4 – 3.0 kb min^−1^ range reported in Fukaya et al. (2017).

Appendix 9–Figure 1F shows how a simple adjustment to our simulation approach, wherein the elongation time steps *w* for individual RNA polymerase molecules were drawn from a Gaussian distribution with mean *μ_w_* = 7 and standard deviation *σ_w_* = 2.5 time steps can qualitatively reproduce the wider profile observed in experimental data, indicating that our observations are indeed consistent with the presence of variability in RNA polymerase elongation times. Additional experimental and theoretical work will be necessary to uncover the biological source of this variability.

In light of the ambiguity introduced by the broad second derivative peak exhibited by our experimental data, we also verified that our inference was robust to the choice of *τ*_elong_, testing cases where *τ*_elong_ = 120 s and *τ*_elong_ = 160 s (see below).

### mHMM inference is insensitive to small changes in RNA polymerase dwell time

Due to the uncertainty in our estimate of *τ*_elong_, we conducted sensitivity studies to ensure that our inference results were robust to our input assumption for *w*. As shown in Appendix 9–Figure 2, we conducted time-averaged mHMM inference on our experimental data assuming different values of *w*. Based upon our autocorrelation analysis, *w* values of 6,7 and 8 seemed the most plausible candidates for the average system elongation time (see Appendix 9–Figure 1E). While small quantitative difference are apparent across these three cases, the results for different values of *w* generally showed a constant offset throughout the embryo, such that qualitative trends were largely robust to the assumed *w* value.

**Appendix 9 Figure 2.**
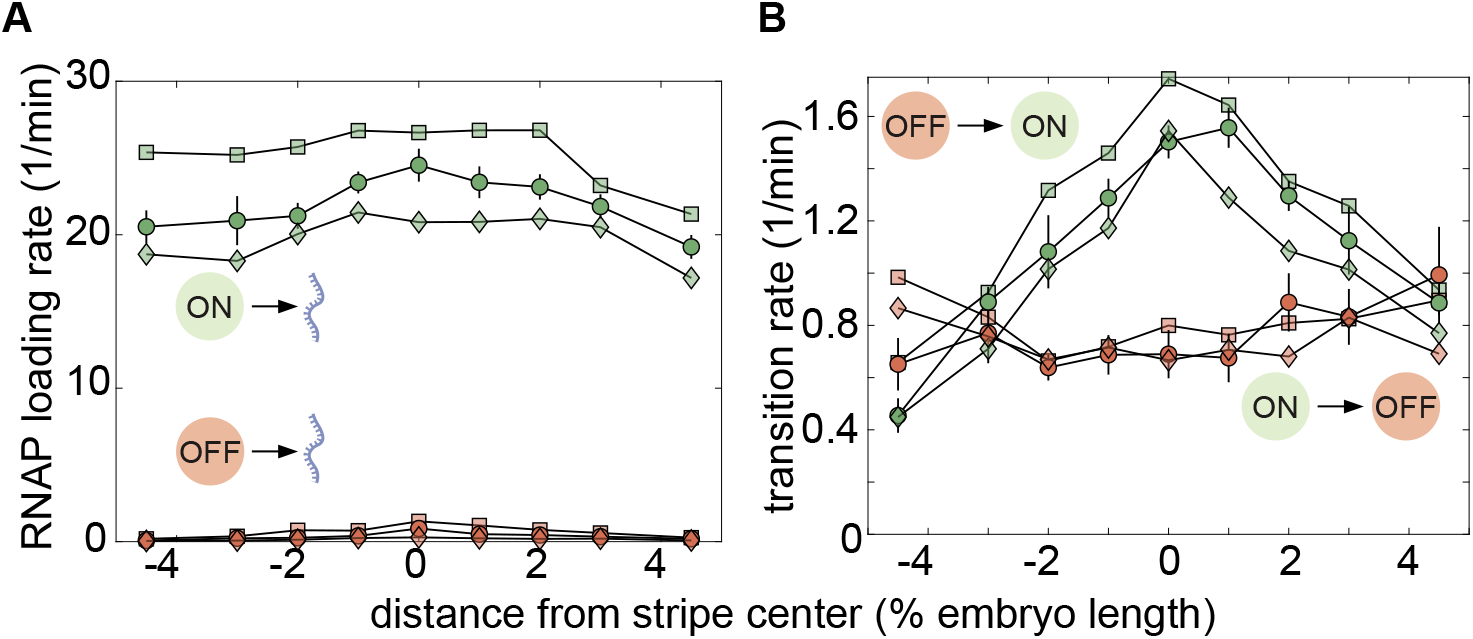
Elongation time sensitivities. Square, circle, and diamond symbols denote inference results for memory time window values *w* of 6, 7, and 8 (*τ*_elong_ of 120s, 140s, and 160s), respectively. Parameter plots for *w* = 7 case are bolded. Bootstrap errors are shown for *w* = 7 case (error magnitudes are comparable across conditions). **(A)** Although the absolute magnitude of the inferred effective initiation rate varies by approximately 10 to 25% across the three conditions, we found that the AP trends (or lack thereof) are robust to our choice of memory. **(B)** Transition rates also exhibit a high degree of robustness to the *w* used for inference. While we observed moderate variation in the inferred magnitude of *k*_on_ (green markers), AP trends are insenstive to *w* assumed for inference within the range tested. Very little variation was observed in *k*_off_ (red markers) across conditions. (Error bars indicate magnitude of difference between first and third quartiles of mHMM inference results for bootstrap samples of experimental data. See Materials and Methods for details.)

---

**Figure 2–Figure supplement 1.**
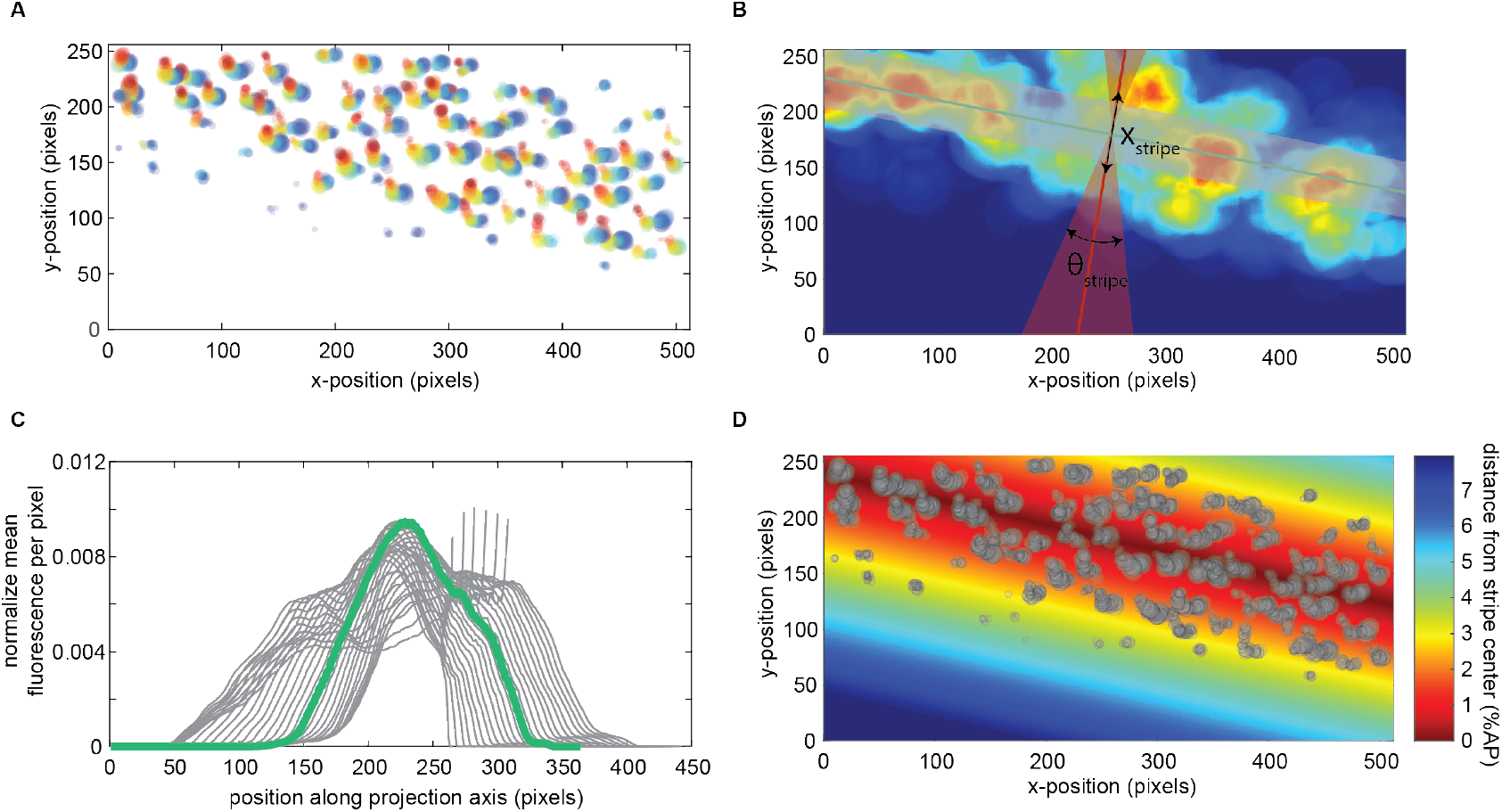
Aligning stripes from multiple embryos. In order to minimize alignment errors when combining data from across multiple *Drosophila* embryos, an automated routine was employed to define a new experimental axis for each data set based upon the spatial distribution of transcriptional activity in the mature *eve* stripe 2 pattern. **(A)** Example of the spatiotemporal distribution of observed fluorescence for an individual embryo. Each circle corresponds to the fluorescence from a single locus at a single point in time. Only observations after 30 min into nuclear cycle 14 were used. Circle size indicates fluorescence intensity. Color indicates temporal ordering: 30 min (blue) to 47 min (red). **(B)** A Gaussian filter was convolved with the raw data points in (A). This filtering ameliorated stripe fitting artifacts that arose due to the relative sparsity of the raw data. The fitting procedure considered both a range of possible stripe orientations (*θ*_stripe_) and, within each orientation, a range of possible positions of the stripe along the anterior-posterior axis (*x*_stripe_) that, together constituted a set of possibilities for the new stripe center position and orientation. Here, the shaded red region indicates the range of values for *θ*_stripe_ that were considered. The red line indicates the best stripe axis inferred by the algorithm and the green line indicates the corresponding optimal stripe center. No constraints were placed on *x*_stripe_, save for the limits of the experimental field of view. **(C)** For each proposed stripe orientation (*θ*_stripe_), a projected stripe profile was generated by taking the average pixel intensity for each position, *x_i_*, along the proposed stripe axis. To determine the optimal center location for each orientation, a sliding window with a width equal to 4% of the embryo length was used to determine the fraction of the total profile fluorescence that fell within 2% embryo length of the stripe center. For example, the gray shaded region in (B) illustrates what this range would be for the green stripe center line (B). This fraction of the total profile was used as a baseline for the comparison of potential stripe center positions. The *θ*_stripe_ and *x*_stripe_ that maximized this metric (green profile in (C)) were taken to define a new, empirically determined stripe center. **(D)** This inferred stripe position defined an experimental axis for each embryo that was used to aggregate observations from across embryos. Gray circles indicate experimental observations (size corresponds to intensity as in (A)) and shading indicates distance from inferred stripe center.

**Figure 2–Figure supplement 2.**
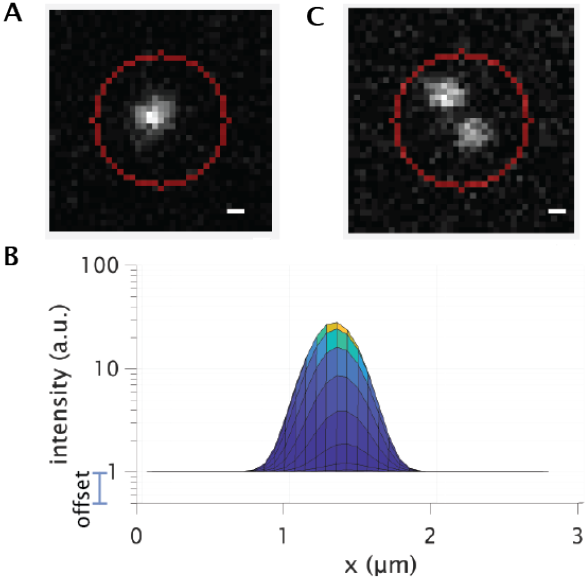
Integrating MS2 Spots. **(A)** Fluorescence of a site of nascent transcript formation is measured by integrating raw pixel intensities in a circular region around the fluroescent MS2 spot of a predefined area (indicated by the red circle) and then subtracting off the background intensity obtained as outlined in **(B)**. (B) X-Z projection of 2D Gaussian function fitted to MS2 spot shown in (A). Background intensity is estimated using the offset value fo this Gaussian fit. The per-pixel offset is then multiplied by the area of the integration region. This background value is then subtracted from the fluorescence integrated across the area shown in (A). **(C)** The radius was chosen to be large enough to integrate the intensities from both sister chromatids, even when they are spatially separated and distinguishable.

**Figure 3–Figure supplement 1.**
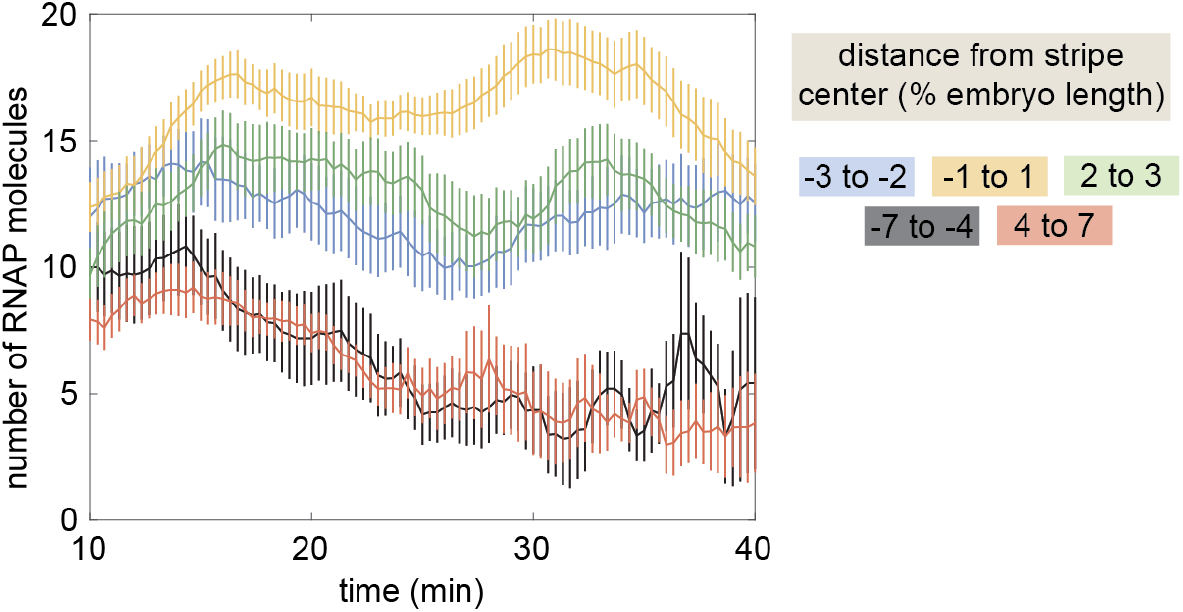
Mean transcriptional activity. Mean transcriptional activity as a function of time for different positions along the stripe. (Average over 11 embryos, error bars indicate bootstrap estimate of the standard error of the mean. See Materials and Methods).

**Figure 3–Figure supplement 2.**
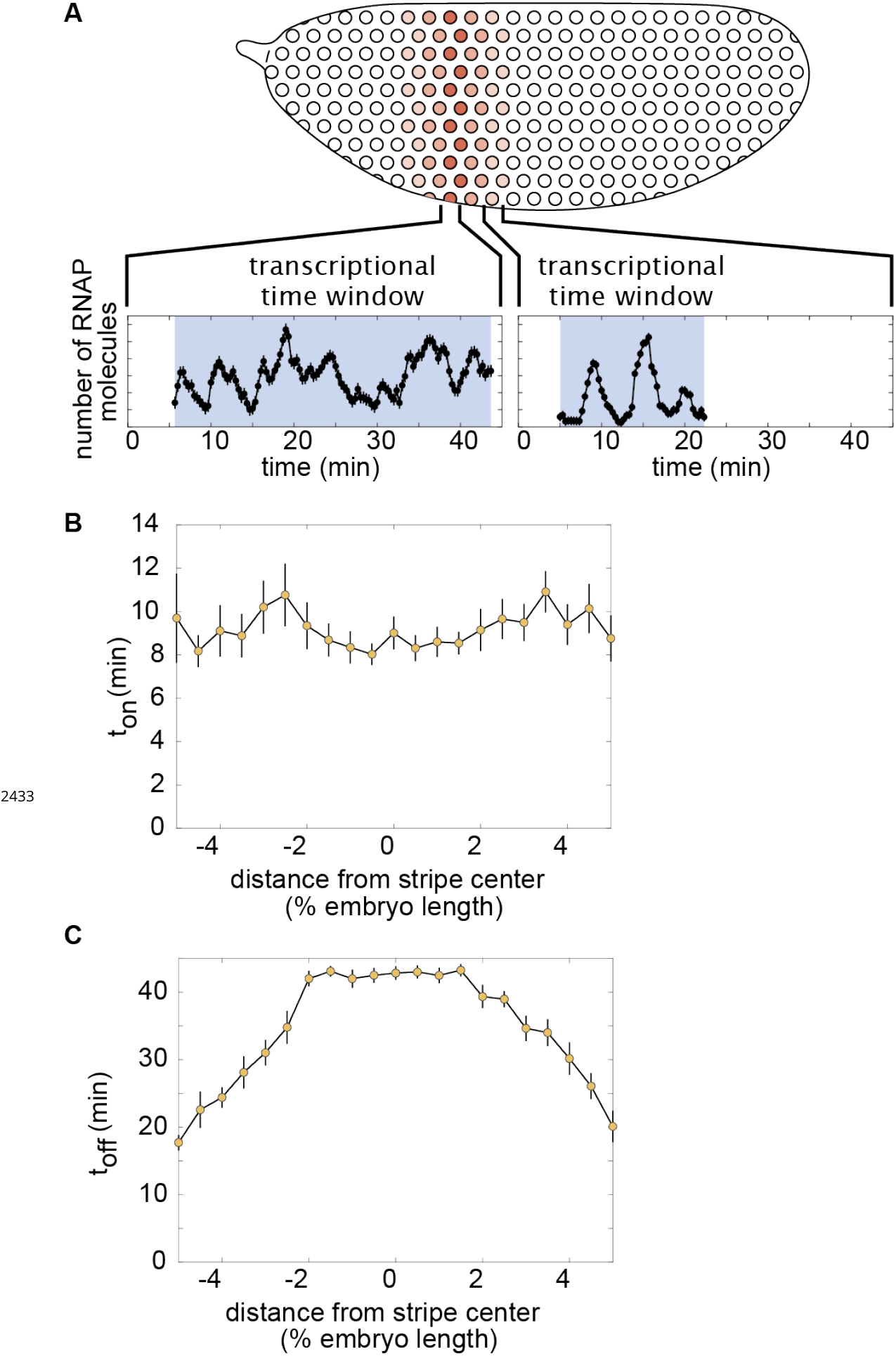
Regulation of the transcriptional time window. **(A)** Singlenucleus measurements reveal that the duration of transcription is modulated along the stripe and that nuclei transcribe in a burst-like fashion. **(B)** Time for nuclei to activate transcription after mitosis, *t*_on_, as a function of position along the stripe. **(C)** Time for nuclei to enter the quiescent transcriptional state, *t*_off_. (B,C, average over 11 embryos, error bars indicate bootstrap estimate of the standard error of the mean. See Materials and Methods).

**Figure 3–Figure supplement 3.**
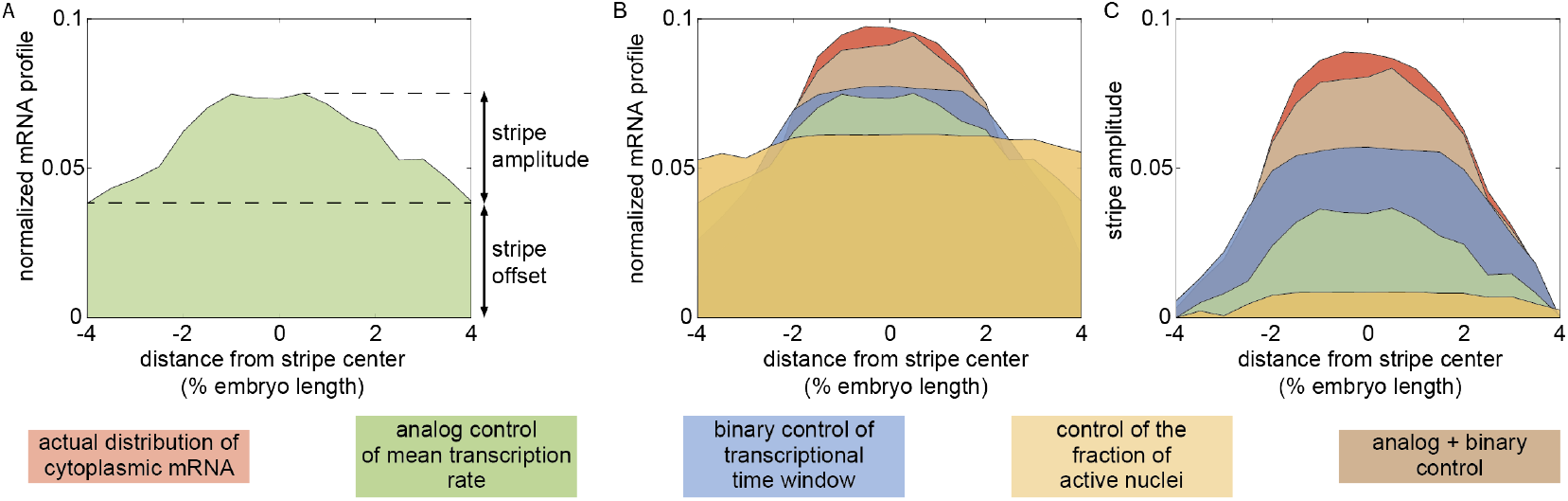
Definition of stripe amplitude. **(A)** The normalized mRNA profile for the stripe can be separated into an offset and an amplitude. **(B)** Normalized mRNA profiles and **(C)** stripe amplitude for the cytoplasmic pattern of mRNA as well as for the contributions from the various regulatory strategies.

**Figure 3–Figure supplement 4.**
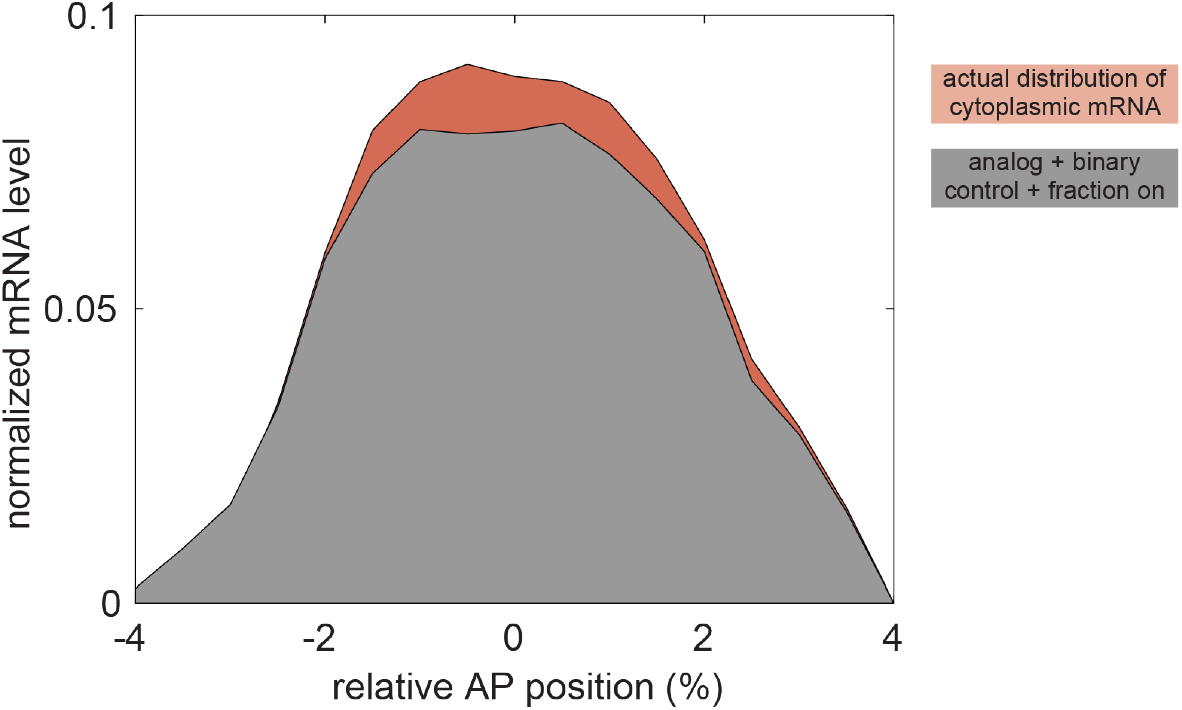
Joint effect of mean rate, binary control, and fraction of active nuclei. Including of the predicted effect of anterior-posterior-dependent modulation of the fraction of active nuclei has little effect on the predicted cytoplasmic mRNA profile (compare brown profile in Figure 1G, gray profile above). The remaining difference between the full profile (red) and the gray profile can be attributed the effects of temporal variations in the mean rate of transcription.

**Figure 5–Figure supplement 1.**
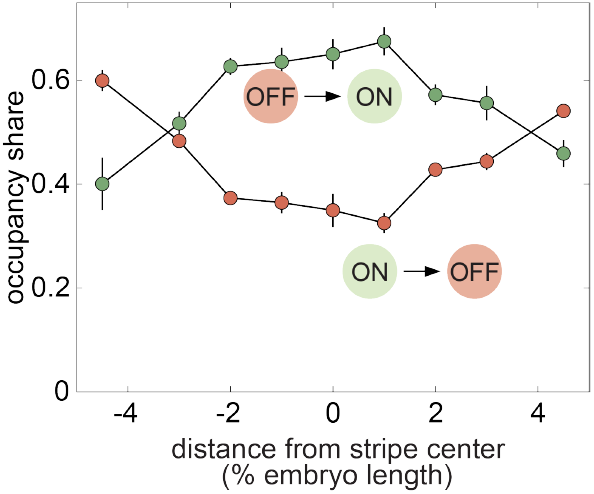
Fraction of time spent in each transcriptional state. Fraction of time spent in the ON and OFF states as a function of the position along the stripe. (Error bars indicate the magnitude of the difference between the first and third quartiles of mHMM inference results for bootstrap samples of experimental data. See Materials and Methods for details.)

